# Morphing Cholinesterase Inhibitor Amiridine into Multipotent Drugs for the Treatment of Alzheimer’s Disease

**DOI:** 10.1101/2024.02.15.580468

**Authors:** Eva Mezeiova, Lukas Prchal, Martina Hrabinova, Lubica Muckova, Lenka Pulkrabkova, Ondrej Soukup, Anna Misiachna, Jiri Janousek, Jakub Fibigar, Tomas Kucera, Martin Horak, Galina F. Makhaeva, Jan Korabecny

**Author notes:** corresponding authors: Martin Horak,; Galina F. Makhaeva,; Jan Korabecny.

## Abstract

The search for novel drugs to address the medical needs of Alzheimer’s disease (AD) is an ongoing process relying on the discovery of disease-modifying agents. Given the complexity of the disease, such an aim can be pursued by developing so-called multi-target directed ligands (MTDLs) that will impact the disease pathophysiology more comprehensively. Herewith, we contemplated the therapeutic efficacy of an amiridine drug acting as a cholinesterase inhibitor by converting it into a novel class of novel MTDLs. Applying the linking approach, we have paired amiridine as a core building block with memantine/adamantylamine, trolox, and substituted benzothiazole moieties to generate novel MTDLs endowed with additional properties like *N*-methyl-D-aspartate (NMDA) receptor affinity, antioxidant capacity, and anti-amyloid properties, respectively. The top-ranked amiridine-based compound **5d** was also inspected by *in silico* to reveal the butyrylcholinesterase binding differences with its close structural analogue **5b**. Our study provides insight into the discovery of novel amiridine-based drugs by broadening their target-engaged profile from cholinesterase inhibitors towards MTDLs with potential implications in AD therapy.

## Introduction

Alzheimer’s disease (AD) is a neurodegenerative condition primarily dominating the population over 65 years [1], being associated with cognitive disturbances and severe progression [2]. As novel drug candidates for AD keep failing in different phases of clinical trials, it can be claimed that the field of AD drug discovery reached its limit. The management of AD symptoms accounts for acetylcholinesterase (AChE, E.C. 3.1.1.7) inhibitors represented by rivastigmine, galantamine, and donepezil (Fig. 1), providing symptomatic relief by targeting cholinergic deficit typical for AD pathology [3]. Memantine (Fig. 2) is used for patients with moderate-to-severe AD, acting as an antagonist of *N*-methyl-D-aspartate receptors (NMDARs) [4]. The blockade of extrasynaptic NMDARs by memantine prevents excessive glutamate overstimulation, thus mitigating excitotoxicity [5]. Recently, two new drugs, aducanumab and lecanemab, were approved by the United States Food and Drug Administration agency in 2021 and 2023, respectively. These drugs act as monoclonal antibodies directed toward amyloid beta (A*β*) [6,7]. Abundance of A*β* plagues (extracellular deposits of A*β* proteins) together with neurofibrillary tangles (aggregates of hyperphosphorylated tau proteins) are characteristic features of AD [8]. Sodium oligomannate (GV-971) is a mixture of marine-derived oligosaccharides conditionally approved in the People’s Republic of China in 2019 for AD treatment [9]. This perspective for AD treatment is very interesting because of the recently discovered relationship of the gut microbiota-to-brain microglia (macrophage cells that are the main form of active immune defense in the central nervous system (CNS)), which means that dietary intake influences learning and memory as well as the health of the nervous system. The mechanism of action of GV-971 is unclear; however, it was proposed that stimulating and reconditioning of gut microbiota suppresses neuroinflammation [10].

**Fig. 1.**
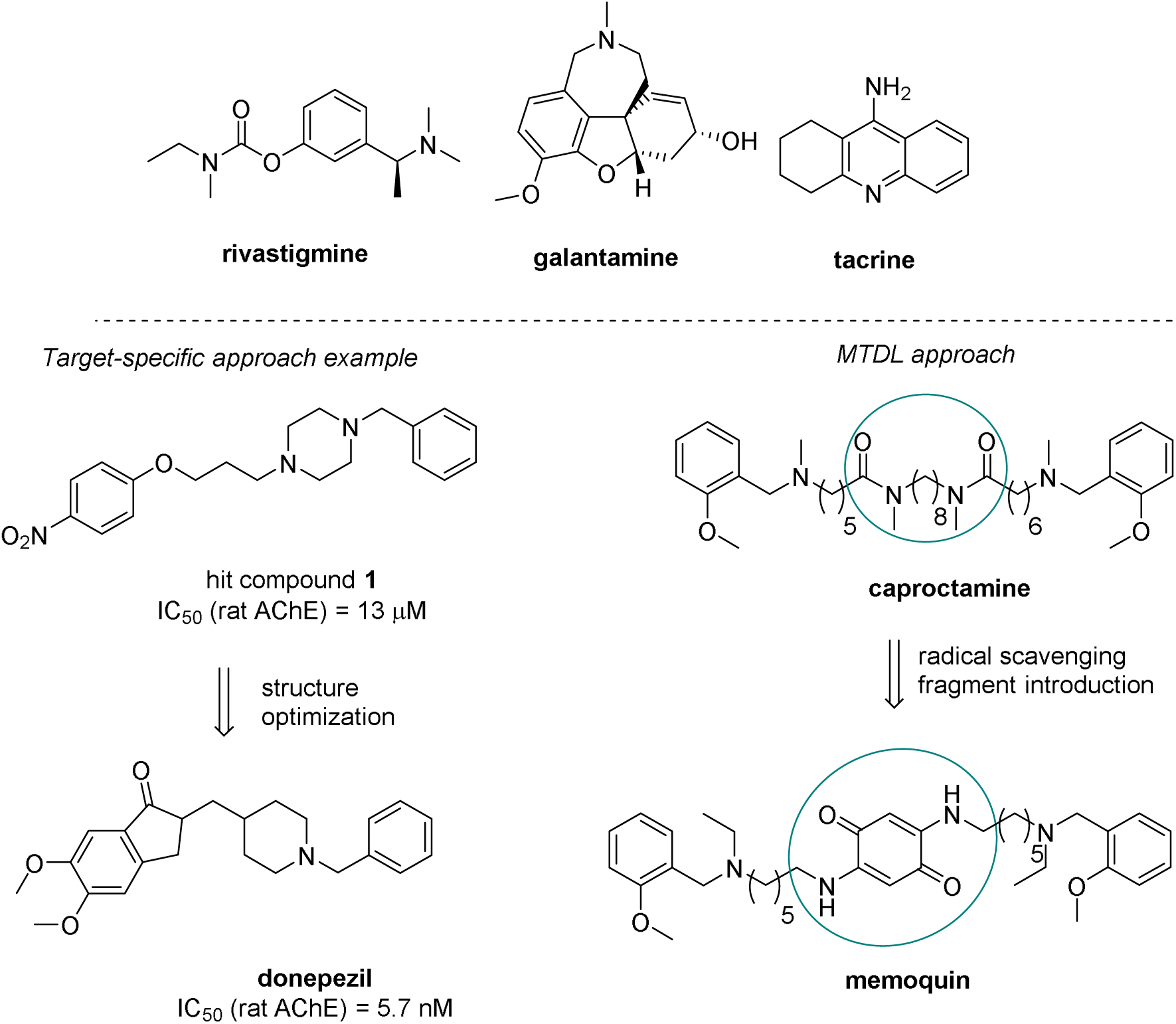
Chemical structures of rivastigmine, galantamine, and tacrine as representatives of cholinesterase inhibitors. Approaches to novel drugs for AD treatment on the selected candidates are displayed.

**Fig. 2.**
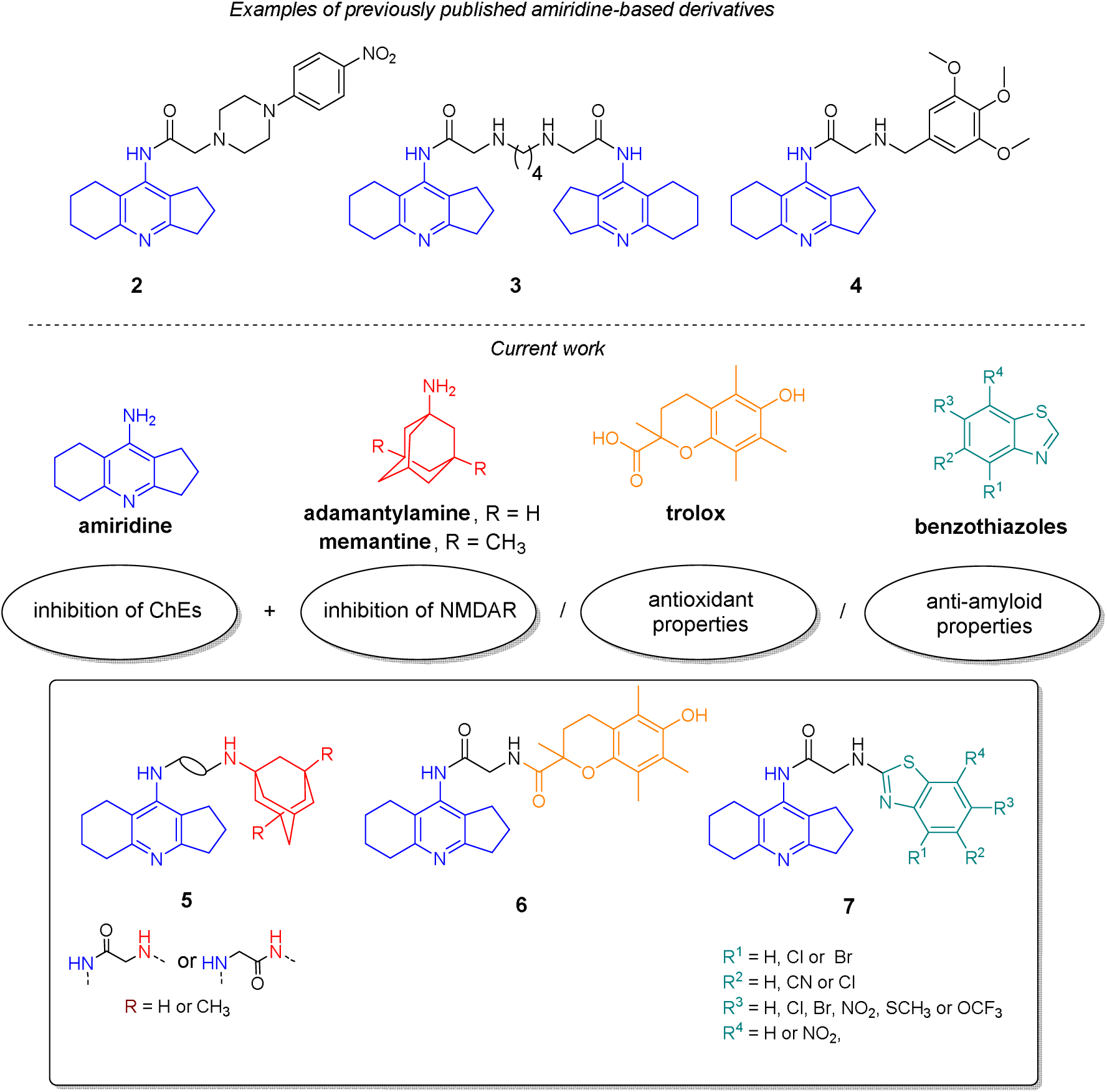
Examples of previously published amiridine-based derivatives and design strategy applied in the current study below, using various pharmacophores.

The pathophysiology of this disease is intricately linked to distinct biochemical processes involving various neurotransmission systems. These encompass aberrant protein processing, neuroinflammation, mitochondrial dysfunction, biometal dyshomeostasis, and oxidative stress. The complexity of AD neuropathology is the culprit, making disease understanding elusive. Another factor is the prolonged lifespan during which AD continuously evolves. These factors collectively underscore the ongoing pursuit of novel drugs for AD treatment, aimed at devising disease-modifying strategies while unraveling their underlying mechanisms and origins. Approaches to drug development can broadly be categorized into two main groups based on the number of targeted pathways. An illustrative example of the classical target-specific approach is discovering the well-known AD drug, donepezil (Fig. 1) [11]. The journey began in 1983 through random screening, during which the *N*-benzylpiperazine derivative **1** (Fig. 1), originally synthesized for the study of anti-arterial sclerosis, was identified as an AChE inhibitor. Its unique structure served as a starting point for further refinements, with a focus on enhancing anti-AChE properties and bioavailability [11,12]. Therapeutics that simultaneously address at least two disease-related targets, which may belong to different sub-pathologies, are often referred to as multi-target drugs (MTDs) or multi-target-directed ligands (MTDLs). Caproctamine (Fig. 1), for instance, was developed to enhance cholinergic activity by not only inhibiting AChE (thereby reducing acetylcholine hydrolysis rates) but also increasing ACh release in the synaptic cleft, achieved by blocking presynaptic muscarinic ACh autoreceptors [13]. Further refinements of the caproctamine scaffold, such as substituting the linear alkyl chain with a radical scavenger moiety, led to the development of memoquin (Fig. 1). This compound exhibited additional properties of interest for AD treatment [13].

The aforementioned approaches have paved the way for the development of several promising compounds aligned with the implied strategies for treating AD, a condition characterized by its complex nature. Many newly discovered compounds rely on well-established pharmacophores with known properties. For instance, pharmacophores exhibiting inhibitory activity against both cholinesterases (ChEs) are usually employed in fragment-based drug design, such as tacrine (Fig. 1), *N-*benzylpiperidine (a part of donepezil), or benzylalkylamine (a part of caproctamine or memoquin). These fragments can be integrated with complementary substructures to confer additional desired properties *via* the so-called ’linking approach,’ wherein the respective pharmacophores are interconnected, mostly using carbon-based linkers. The research presented herein exploits similar development approach, wherein the amiridine scaffold (Fig. 2) served as the core with anti-ChEs activities, combined with supplementary structures. Our primary objective was to specifically explore the potential of amiridine as such a pharmacophore and to diversify the pool of amiridine-based hybrid compounds.

## 1. Design

Amiridine (9-amino-2,3,5,6,7,8-hexahydro-1*H*-cyclopenta[*b*]quinoline; Fig. 2) has been identified as a reversible inhibitor of both ChEs, with a particular preference for butyrylcholinesterase (BChE; E.C. 3.1.1.8) [14]. Its structural features resemble that of tacrine (9-amino-1,2,3,4-tetrahydroacridine) (Fig. 1), a well-known ChEs inhibitor exhibiting greater selectivity towards BChE. Tacrine was the first drug from the ChE inhibitor family to be approved for AD treatment. However, it was withdrawn from the market in 2013 due to serious side effects. The absence of hepatotoxicity in amiridine in comparison to tacrine represents its major advantage [14–16]. On the other hand, from the medicinal chemistry standpoint, amiridine displays lower reactivity than tacrine, with fewer amiridine-based derivatives available in the literature. Recently, Makhaeva *et al.* synthesized several such compounds, including amiridine-piperazine hybrids (exemplified by **2**; Fig. 2) [17], bis-amiridines (represented by **3**; Fig. 2) [18], and thiourea-containing amiridine derivatives (exemplified by **4**; Fig. 2) [19]. In all the cases, the amiridine core was functionalized at the endocyclic nitrogen with either *N*-acyl- or *N*-thiourea-based spacers.

To expand the family of amiridine-based hybrids, we combined this pharmacophore with structures known for their respective biological activities related to AD pathophysiology. Specifically, we used memantine, 1-adamantylamine, trolox, and various unsubstituted/substituted benzothiazoles to enhance the anti-ChEs activity of amiridine, introducing NMDAR antagonist activity, antioxidant, and anti-amyloid properties, respectively (Fig. 2). Similar to previously published hybrids, we substituted the amiridine skeleton at the endocyclic nitrogen with *N*-acyl-based spacers. In the case of derivatives containing either adamantylamine or memantine, we prepared pairs that differed in the configuration of the amide linker to assess how this structural change impact biological activity. We were also interested in whether the hybrid combining amiridine and trolox would retain antioxidant properties. The last group of compounds were benzothiazole-based derivatives. Benzothiazole moiety has been utilized in the structures of numerous fluorescent probes for the detection of amyloid-beta proteins, also exerting anti-amyloid properties [20,21]. Thus, amiridine-benzothiazole hybrids were prepared with unsubstituted or substituted (Br, Cl, CN, NO_2_, SCH_3_, or OCF_3_) benzothiazole moiety.

## 2. Results and Discussion

### 2.1. Synthesis of Amiridine-based Heterodimers

Compounds **5a-d** were synthesized using a two-step procedure (Scheme 1). In the first step, acyclic nitrogens in amiridine, 1-adamantylamine (**9**), and memantine (**10**) were functionalized with chloroacetic acid. The acylation reactions were conducted employing a slightly modified method previously reported by Makhaeva *et al.* [17]. Subsequently, the chlorine atom in the resulting amides **8**, **11**, and **12** was replaced with an amine group originating from **9**, **10**, or amiridine, respectively. This nucleophilic substitution was carried out in CH_3_CN in the presence of K_2_CO_3_ and KI. The final derivatives, denoted as **5a-d**, were isolated in good yields, ranging from 41% to 54%.

**Scheme 1.**
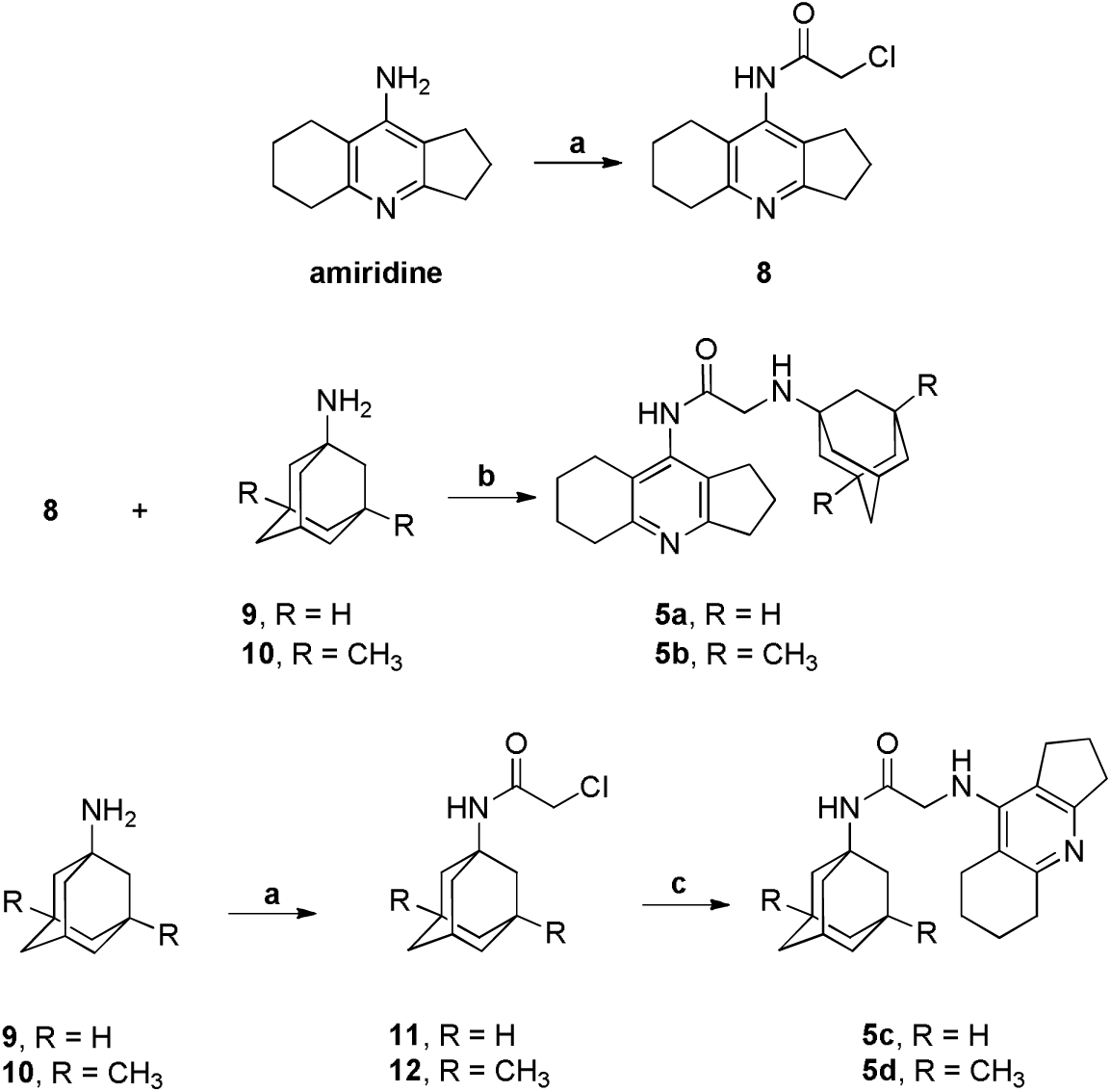
Preparation of the amiridine-based compounds **5a-d**. Reagents and conditions: a) 2-chloroacetyl chloride (4 eq.), CHCl_3_, 90 °C, overnight, **8**, 54%, **11**, 90%, **12**, 52 %; b) CH_3_CN, K_2_CO_3_, KI, reflux, 3 h, **5a**, 54%, **5b**, 54%; c) amiridine (1.1 eq), CH_3_CN, K_2_CO_3_, KI, reflux, overnight, **5c**, 48%, **5d**, 41%.

Intermediate **8** underwent transformation into compound **13** through a method known as the Gabriel amine synthesis. In this process, the phthalimide anion (derived from potassium phthalimide) reacted with compound **8**, and subsequent hydrolysis with KOH yielded the desired amine **13** in a 57% yield (Scheme 2). The substitution of the chlorine atom in **8** to the amino group in **13** was confirmed through LC-MS analysis (Supporting Information). Intermediate **13** was employed in the synthesis of the amiridine-trolox hybrid **6** (Scheme 2) and also in the preparation of amiridine-benzothiazoles **7a-m** (Scheme 3).

The activation of trolox was achieved using benzotriazol-1-yloxytris(dimethylamino)phosphonium hexafluorophosphate (BOP) in the presence of triethylamine (TEA) in dimethylformamide (DMF), resulting in the corresponding activated ester as an intermediate. Subsequent coupling between trolox and **13** afforded the amiridine-trolox hybrid **6** in an overall yield of 86% (Scheme 2).

For the *N*-alkylation of intermediate **13**, different 2-chlorobenzothiazoles **14** were used as alkylating agents (Scheme 3). The unsubstituted amiridine-benzothiazole hybrid, **7a**, was prepared under neat conditions (no solvent and no base addition). The remaining amiridine-benzothiazoles, **7b-m**, introduced a halogen atom (chlorine, bromine) or various functional groups (nitrile, nitro, methyl sulfide, or methoxy group). These were synthesized in the presence of nine equivalents of *N*,*N*-diisopropylethylamine (DIPEA). **7a-m** were obtained with satisfactory yields ranging from 21% to 77%. Notably, all the amiridine-benzothiazole hybrids were endowed with low solubility in common organic solvents, with the exception of DMSO.

**Scheme 2.**
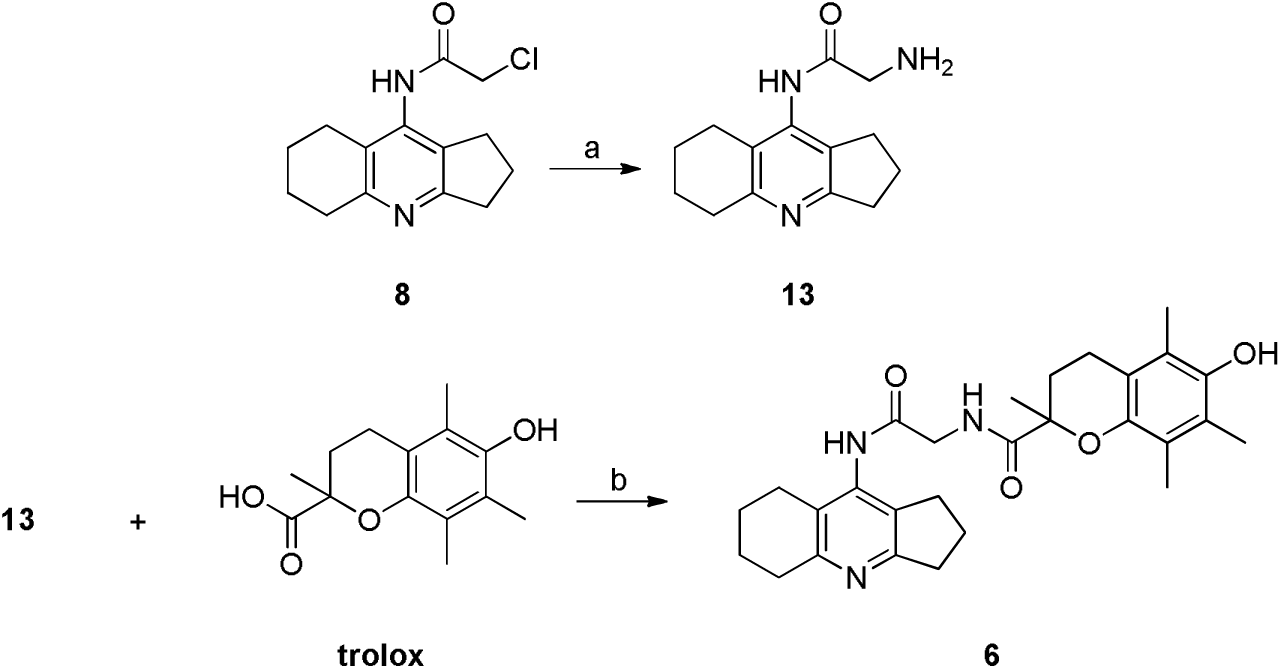
Preparation of intermediate **13** and final compound **6**. Reagents and conditions: a) potassium phthalimide, CH_3_CN, reflux, 3 h, then an excess of NH_2_NH_2_.H_2_O, reflux, overnight, 57%; b) DMF, TEA, BOP, room temperature, 2 days, 86%.

**Scheme 3.**
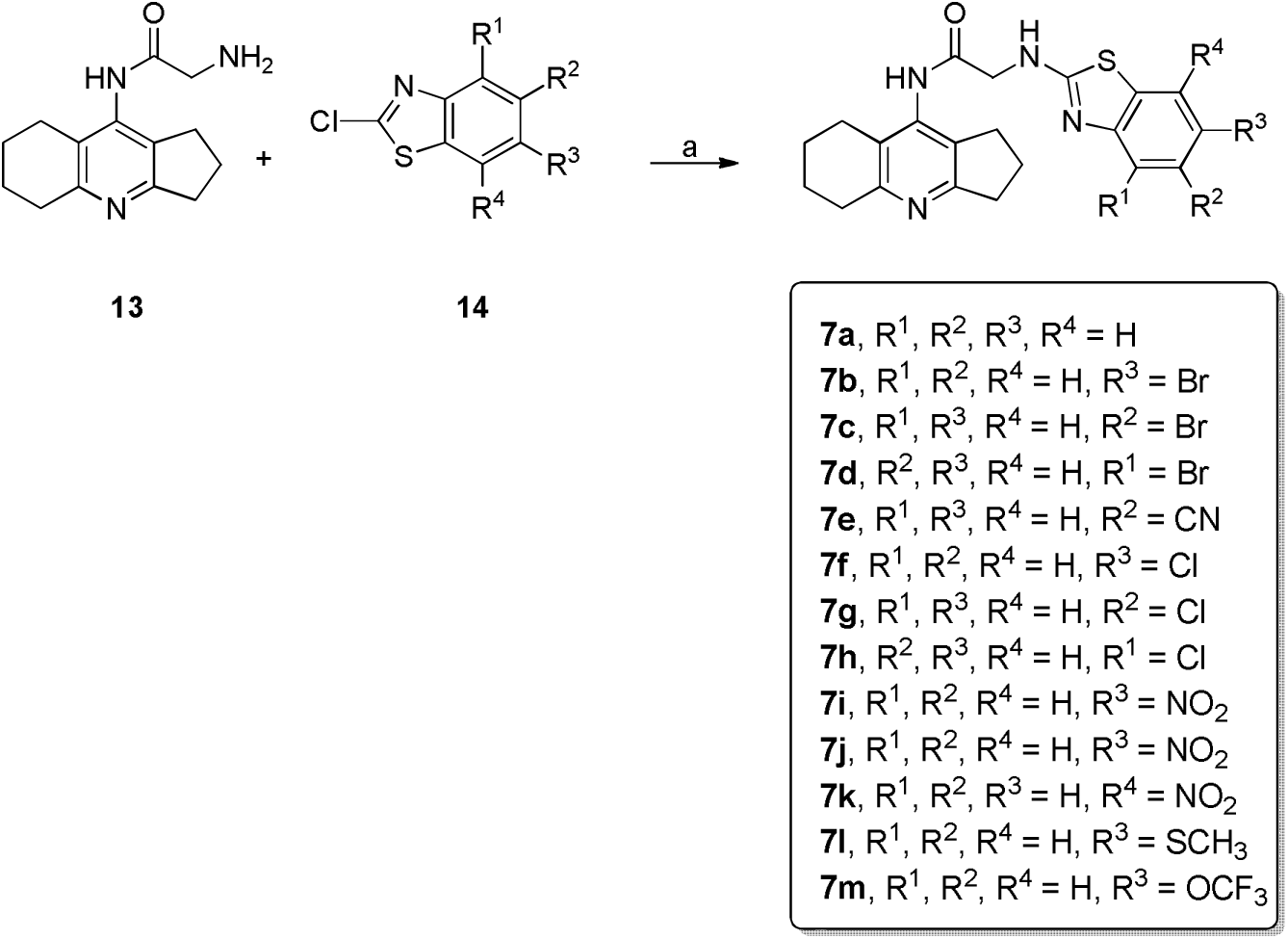
Preparation of amiridine-benzothiazole derivatives **7a-m**. Reagents and conditions: a) for **7a**: 2-chlorobenzothiazole, 110 °C, overnight, 32%; for **7b-m**: corresponding 2-chlorobenzothiazole, DIPEA, 100 °C, overnight, 21-77%.

### 2.2. Evaluation of Cholinesterase Inhibition Activity

ChEs are enzymes responsible for the hydrolysis and metabolism of various endogenous and exogenous substrates [22], that are broadly studied in association with disease conditions like myasthenia gravis or AD. The primary attention in the development of AD drugs has been paid to AChE inhibitors. Nevertheless, there is growing interest in nonselective or BChE-selective inhibitors due to the compensatory role of BChE when AChE levels are depleted during the progression of AD [23]. Selective BChE inhibitors may be beneficial for patients in moderate-to-severe stages of AD [23,24]. Given the presence of the amiridine scaffold in the structures of the novel compounds, we hypothesized that derivatives would exhibit activity against ChEs with selectivity towards BChE. Accordingly, we assessed the inhibition activities of these novel compounds against human recombinant AChE (*h*AChE) and human serum BChE (*h*BChE) following a modified spectrophotometric protocol developed by Ellman *et al.* [25]. Initial screening of compounds **5–7** was performed at a concentration of 1 µM, and subsequently, for the compounds exerting inhibition ≥ 25%, IC_50_ values were determined. Results for four compounds (**5c-d**, **7c**, and **7g**) are summarized in Table 1. Complete data are available in the Supplementary Information (Table S1), including the inhibition percentages for all derivatives. Amiridine hydrochloride (amiridine.HCl) and tacrine (THA) were used as reference compounds. Compounds **5–7** displayed weak inhibition potency against *h*AChE (1 – 18% at 1 µM concentration). The most active compound was **7b**, bearing a bromine atom at position 6 of the benzothiazole moiety, followed by **7c** with a bromine atom at position 5. Only a superficial structure-activity relationship for AChE inhibition can be drawn from the obtained data. Analogues containing the memantine moiety (**5b** and **5d**) exhibited greater activity than their counterparts with the adamantylamine group (**5a** and **5c**). The presence of any substituent on the benzothiazole core enhanced anti-AChE activity, with the best-tolerated position for the substituent being 6 for derivatives with halogen atoms (**7b-d** with bromine or **7g-h** with chlorine). *h*BChE inhibition properties of **5**–**7** ranged widely (0.9 – 95% at 1 µM concentration). The most potent inhibitors were derivatives **5c-d**, containing the adamantylamine and memantine moieties, for which IC_50_ values were determined (0.6 and 0.1 µM, respectively). Similar to AChE inhibition, analogues containing the memantine moiety (**5b** and **5d**) displayed more profound activity against BChE than their counterparts with the adamantylamine group (**5a** and **5c**). Additionally, the nature of the linker also influenced inhibition properties, with the linker featuring an amide part on the adamantylamine or memantine side being more favorable for BChE inhibition properties. Among the benzothiazole-containing derivatives, the most effective inhibitors were **7c** and **7g** (25% and 26% inhibition at 1 µM concentration, respectively), with IC_50_ values also determined (4.9 and 11 µM, respectively). Notably, the presence of bromine (**7c**) or chlorine (**7g**) atoms as substituents at position 5 of the benzothiazole segment enhanced anti-BChE properties. It is worth mentioning that most final derivatives exhibited better inhibition properties against *h*BChE than *h*AChE. **5c-d** displayed IC_50_ values for BChE inhibition in the submicromolar range and were equally potent as amiridine (IC_50_ = 0.18 µM) and tacrine (IC_50_ = 0.08 µM) [17,26].

**Table 1.** hBChE inhibitory activities of **5c-d**, **7c** and **7g** and reference compounds (amiridine hydrochloride and THA); their cytotoxicity profile on SH-SY5Y cell line, and predictions of BBB penetration.

| Compound | hBChE<br>$IC_{50} \pm SEM$<br>($\mu M$)<sup>a</sup> | Cytotoxicity<br>$IC_{50} \pm SEM$ ($\mu M$)<sup>a</sup> | | BBB permeation estimation | | |
| --- | --- | --- | --- | --- | --- | --- |
| | | SH-SY5Y | HepG2 | $P_e \pm SEM$<br>($10^{-6} cm.s^{-1}$)<sup>b</sup> | CNS<br>(+/-)<sup>c</sup> | BBB<br>score<sup>f</sup> |
| **5c** | $0.60 \pm 0.02$ | $\geq 200$ | $190 \pm 23$ | $1.6 \pm 0.2$ | CNS - | 4.9 |
| **5d** | $0.109 \pm 0.004$ | $110.0 \pm 1.5$ | $42 \pm 6.1$ | $2.20 \pm 0.05$ | CNS<br>+/- | 4.8 |
| **7c** | $4.9 \pm 0.4$ | $18.0 \pm 2.3$ | $22.0 \pm 0.2$ | $2.1 \pm 1.0$ | CNS<br>+/- | 4.1 |
| **7g** | $11.0 \pm 1.2$ | $98.0 \pm 5.7$ | $\geq 200$ | $2.8 \pm 0.8$ | CNS<br>+/- | 4.1 |
| Amiridine.HCl<sup>d</sup> | $0.18 \pm 0.01$ | $600 \pm 25$ | n.d.<sup>g</sup> | $5.8 \pm 0.5$ | CNS + | n.d.<sup>g</sup> |
| THA<sup>e</sup> | $0.080 \pm 0.001$ | $120.0 \pm 1.6$ | $19.4 \pm 0.8^h$ | $6.0 \pm 0.6$ | CNS + | n.d.<sup>g</sup> |

<sup>a</sup> Results are expressed as the mean of at least three experiments; <sup>b</sup> results are expressed as the mean of at least two experiments; <sup>c</sup> classification of the prediction to cross the BBB, CNS+: high BBB permeation predicted with $P_e$ ( $10^{-6} cm.s^{-1}$ ) $> 4.0$ , CNS-: low BBB permeation predicted with $P_e$ ( $10^{-6} cm.s^{-1}$ ) $< 2.0$ , CNS+/-: BBB permeation uncertain with $P_e$ ( $10^{-6} cm.s^{-1}$ ) from 4.0 to 2.0; <sup>d</sup> taken from reference [17]; <sup>e</sup> taken from references [26,27]; <sup>f</sup> calculated according to reference [28]; <sup>g</sup> n.d. = not determined; <sup>h</sup> taken from reference [29].

### 2.3. Docking Studies

To gain insight into the structural requirements responsible for the top-ranked BChE inhibitor **5d** activity, we performed docking studies in tandem with molecular dynamics (MM/MD). As **5d** is structurally related to derivative **5b**, which turned out to be BChE inactive, we use this compound for comparative purposes and to highlight essential interactions driving the ligand binding. Template structure of human BChE complexed with THA (PDB ID: 4BDS) was obtained from Protein Data Bank, meeting the following conditions: *i)* high-resolution of the solved protein-ligand complex (2.1 Å) being the prerequisite for reliable predictive results, and *ii)* the fact that the complex was elucidated for THA which is a close amiridine analogue, the core scaffold of compounds **5b** and **5d**.

Initially, we validated our MM/MD protocol by comparing complexes of predicted and crystal-embedded THA, resulting in a very high model reliability with RMSD value of 0.938 Å^2^ (Fig. S46). In the crystal structure of THA complexed with human BChE, the ligand binds to the catalytic site, revealing several crucial interactions with the protein [30]. Specifically, THA demonstrates parallel π-π stacking to W82 and hydrogen bond interactions between positively charged aromatic nitrogen *N*7 and H438 from the catalytic triad. Crystal structure of THA in human BChE also revealed a hydrogen-bond web mediated via water molecules to exocyclic amino group *N*15, contacting D70, S79 and T120. As our MM/MD is somewhat limited in terms of determining the water molecules involved in interactions between protein and THA, docked THA (Fig. S46B) also harbored a slightly different pose in the enzyme active site, resulting in the loss of hydrogen bond to H438; however, still facing towards W82 and contacting D70 and S79.

Top-ranked human BChE inhibitor **5d** revealed a very close orientation of amiridine moiety to THA within the enzyme active site (Fig. 3B). Indeed, the amiridine part of the molecule **5d** faces W82 via aromatic stacking. This interaction can be considered less productive as amiridine lacks one aromatic moiety compared to THA. Notably, other hydrogen bond interactions are preserved, counting the contact between charged *N*4 and the amide backbone of H438, and water-mediated connections between amide moiety of **5d** with S79 and D70. Memantine moiety of **5d** is implicated in hydrophobic interactions in the region defined by T120, Q119, S287 and Y332, contributing to ligand anchoring. On the contrary, amiridine-memantine compound **5b**, differing in the position of amide moiety, that resides in the vicinity of the amiridine core, yielded unproductive binding mode (Fig. 3A). Indeed, the compound **5b** is completely distorted in the human BChE active site, making opposite anchoring compared to compound **5d**. The top-scored docking pose of **5b** placed the amiridine moiety in the proximity of D70, T120, S287, F329 and G117. Memantine moiety of **5b** displayed hydrophobic interactions close to W82. The loss of hydrogen bonds that are typically observed in the crystal structure between THA and human BChE (Fig. S46A) or between compound **5d** and human BChE (Fig. 3B), is the culprit for such ligand orientation. It can be speculated that the origin of such ligand accommodation might stem from the decrease in *N*4 *p*K_a_, dropping from 9.62 (estimated for **5d**) to 7.30 (predicted for **5b**; predicted by Chemaxon tool v. 20.15; https://www.chemaxon.com), resulting lower capacity to be protonated and less electron density in the amiridine aromatic region of **5b**. As a consequence, the pivotal interaction between amiridine moiety of **5b** and W82 is less probable.

**Figure 3.**
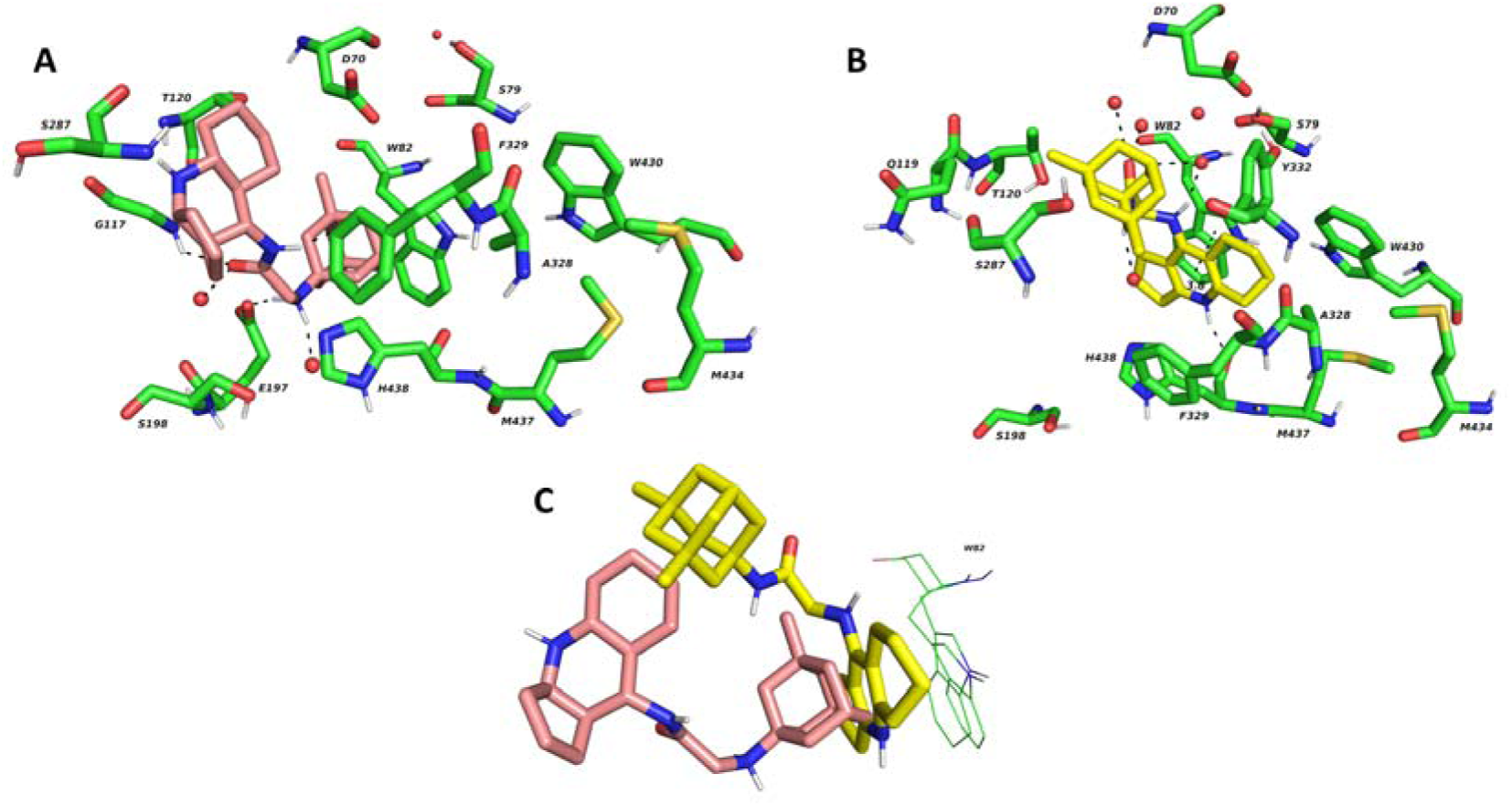
Top scored docking pose of **5b** (A) and **5d** (B) highlighting the key findings responsible for compound activity/inactivity. For the sake of clarity, superimposed ligands are aligned in the Figure C with respect to key amino acid residue W82 to demonstrate the binding difference. Compounds **5b** and **5d** are colored in salmon and yellow, respectively. Essential amino acid residues responsible for ligand anchoring are rendered in green. Important interactions of different origin are displayed with dashed black lines. The figure was created with The PyMOL Molecular Graphics System, v. 2.5.2.

### 2.4. Cytotoxicity

The cytotoxic profile of **5**–**7** was carried through the use of the MTT (3-(4,5-dimethylthiazol-2-yl)-2,5-diphenyltetrazolium bromide) assay on human neuroblastoma (SH-SY5Y) and hepatocellular carcinoma (HepG2) cell lines, following established protocols [31]. The cytotoxicity was assessed on SH-SY5Y cells in undifferentiated and in differentiated form. The data obtained from the measurements on the undifferentiated SH-SY5Y and HepG2 cells are summarized in Table 1 for **5c-d**, **7c** and **7g**, as well as for the reference compounds amiridine hydrochloride and THA. Additional data can be found in Supplementary Information (Table S2 and Fig. S1).

Compounds **5**–**7** were initially tested on SH-SY5Y undifferentiated cells. The determined IC_50_ values exhibited a relatively broad scale for derivatives endowed with either an adamantylamine or memantine scaffold, ranging from 31 to over 200 µM. A similar trend was observed among benzothiazole-based hybrids, reaching IC_50_ values spanning from 9.5 to over 200 µM. Notably, compounds containing adamantylamine moieties (**5a** and **5c**) displayed approximately twice lower toxicity compared to their memantine counterparts. Furthermore, the nature of the linker also influenced the cytotoxicity profile. Specifically, **5c** and **5d**, featuring an amide linker closer to the adamantylamine or memantine moieties, exhibited IC_50_ values approximately three times higher than **5a** and **5b**, respectively. The disparity in IC_50_ values for both BChE inhibition and cytotoxicity against undifferentiated SH-SY5Y cells between **5c**-**d**, and the less toxic **5a**-**b** indicated a low-cytotoxic profile for the former compounds.

Among the amiridine-benzothiazoles, **7a-m**, the substituent character and position notably influenced their cytotoxicity profile. Compound **7b**, bearing a bromine atom at position 6, exhibited the lowest toxicity, with an IC_50_ value ≥ 200 µM. Conversely, its analogs, **7c** and **7d**, possessing bromine atoms at positions 5 and 4, respectively, displayed cytotoxicity twice and more than ten times higher. A similar trend was observed for derivatives containing chlorine substituents, where **7f** (chlorine atom at position 6) was the least toxic, while **7g** and **7h** (chlorine atom at positions 5 and 4, respectively) exhibited approximately 1.5-fold higher cytotoxicity. The IC_50_ values for BChE inhibition and cytotoxicity against undifferentiated SH-SY5Y for **7c** and **7g** lie in the same range, proposing dosing with care.

Similar pattern was observed for HepG2 cell line with IC_50_ values spanning from 42 to 190 µM for derivatives with either an adamantylamine or memantine scaffold, while benzothiazole-based heterodimers exerted cytotoxicity with IC_50_ values between 14 and 200 µM. The only exception in cytotoxicity pattern was found for **7d**, reaching IC_50_ value eight times lower for HepG2 cell line than for SH-SY5Y. Most importantly, all the compounds turned out to be less cytotoxic than tacrine as reference drug.

To expand our understanding of the cytotoxic profiles of **5**–**7**, all compounds were also assessed on SH-SY5Y differentiated cells. For these evaluations, concentrations corresponding to the IC_50_ values obtained for undifferentiated SH-SY5Y cells were applied to each derivative. As depicted in Figure S1, the viability of differentiated cells treated with most tested compounds was either comparable or improved, suggesting similar or reduced toxicity compared to undifferentiated cells. Notably, compound **5b** was minutely inspected, displaying an IC_50_ value of 12 µM.

It is important to note that these results must be considered with precaution, representing preliminary insight into toxicity. To gain a more comprehensive understanding of the cytotoxic profiles of the synthesized compounds, additional investigations are warranted.

### 2.5. *In Vitro* Blood-Brain Barrier Permeation and Druglikeness Predictions

To investigate the potential of compounds **5**–**7** to cross the blood-brain barrier (BBB) and potentially exhibit CNS activity, we employed two assessment tools: the Parallel Artificial Membrane Permeation Assay (PAMPA) and an algorithm referred to as the "BBB score" [32,33,28]. The results for selected compounds (**5c**-**d**, **7c**, and **7g**), along with reference compounds (amiridine and THA), are presented in Table 1, while the complete dataset can be found in the Supplementary Information (Tables S2 and S3). For analogues **5a-d**, the nature of the linker had a significant impact on permeability *via* passive diffusion. Based on permeability (*P*_e_) values for these derivatives, linkers with an amide moiety in proximity to the amiridine side exhibited more favorable outcomes for BBB penetration compared to those positioned on the adamantylamine or memantine side. Specifically, *P*_e_ values for **5a**-**b** fell within the range indicative of high BBB permeation predictions. On the contrary, **5c**-**d** were predicted to possess low to uncertain properties to cross the BBB. Compound **7a** (lacking any substituent on the benzothiazole moiety) exhibited the highest *P*_e_ value within the benzothiazole-based family. Most benzothiazole-substituted derivatives displayed low or uncertain abilities to cross the BBB passively, with the exceptions of **7e** (bearing a nitrile group at position 5) and **7h** (containing a chlorine atom at position 4), indicating high propensity to cross the BBB. BBB penetration was also predicted by the "BBB score", an algorithm proposed by Gupta *et al.* This algorithm relies on five physicochemical descriptors: the number of aromatic rings, heavy atoms, topological polar surface area, p*K*_a_, and a descriptor composed of molecular weight, hydrogen bond donors, and acceptors (MWHBN) [28]. To classify a compound as CNS-active, its BBB score should typically fall within the range of [4–6]. In general, amiridine heterodimers BBB score values fell into two categories: [2–4] and [4, 5]. Out of them, **5a-d** exhibited high BBB scores (4.8 and 4.9, respectively). Among the benzothiazole-based derivatives, compounds such as **7a** (no substitution), **7b-d** (with bromine atoms), and **7f-h** (with chlorine atoms) displayed BBB scores of 4.1, still sufficient for potential BBB permeation. The BBB scores for the rest benzothiazole-based compounds were below 4, signifying a non-CNS active profile.

Additionally, we also assessed the drug-like properties of our compounds using the SwissADME website tool (http://www.swissadme.ch/index.php) [34,35]. Gastrointestinal absorption (GIA), adherence to Lipinski’s rule of five, bioavailability, and Pan Assay Interference Structures (PAINS) violation were all taken into account to estimate the drug-like properties. The comprehensive results are presented in the Supplementary Information (Table S3), along with bioavailability radar charts for all final derivatives (Figure S2). As demonstrated in Table S3, a majority of the amiridine compounds exhibited favorable properties for GIA, with no violations against Lipinski’s rule of five or PAINS. However, the bioavailability radar charts revealed that most of the derivatives were associated with relatively high lipophilicity and poor solubility, for the latter consistently observed during *in vitro* experiments.

### 2.6. NMDA receptor antagonist properties

Excessive activation of NMDARs can lead to neuronal death, generally characterized as excitotoxicity [36]. Ongoing debate centers on which subtypes of NMDARs exhibit involvement in the pathophysiological cascade of AD. To this end, there is a strong evidence that GluN2B-NMDARs are involved in neurodegeneration and Aβ-induced synaptic dysfunction and synapse loss in AD [5,36].

Antagonists targeting GluN2B-NMDARs hold considerable promise as potential therapeutic agents, offering the prospect of neuroprotection and the amelioration of cognitive deficits in individuals afflicted with AD. THA is known to modulate several neurotransmitter pathways, thus exerting a complex mechanism of action [37]. As cholinergic and glutamatergic pathways are the most profound systems in the context of symptom mitigation, in line with the THA dual action on cholinesterase and NMDA receptors [38], we investigated selected newly discovered amiridine derivatives for their activity against GluN1/GluN2B NMDA receptor. Specifically, the compounds possessing adamantylamine and memantine moieties (i.e. **5c-d**) and one benzothiazole derivative (**7m**) were inspected at 30 µM, given their known pharmacological properties at NMDA receptors. Data are expressed as relative inhibitions (RIs) to glutamate control response at membrane potentials of -60 mV, and +40 mV (Table 2). In line with the prerequisite of attaching NMDA receptor antagonistic scaffolds to amiridine core, all the tested compounds exhibited clear voltage-dependent antagonistic activity, although not reaching the activity of memantine.

**Table 2.** Relative inhibitions (RIs) of **5c**-**d** and **7m** and reference compound memantine at recombinant human GluN1/GluN2B NMDA receptor expressed in HEK293 cells

|  | -60 mV |  | +40 mV |  |
| --- | --- | --- | --- | --- |
| Compound | RI<sup>a</sup> (%) ± SEM | n<sup>b</sup> | RI<sup>a</sup> (%) ± SEM | n<sup>b</sup> |
| **5c** | 35.0 ± 2.3 | 5 | 14.0 ± 2.0 | 4 |
| **5d** | 68.0 ± 1.4 | 5 | 37.0 ± 2.1 | 4 |
| **7m** | 63.0 ± 2.6 | 5 | 32.0 ± 2.5 | 4 |
| **Memantine** | 99.0 ± 0.6 | 5 | 53.0 ± 1.0 | 7 |

<sup>a</sup> Relative inhibition (RI) measured at 30 μM of each compound in the indicated membrane potential is presented in percentages with standard error mean (SEM) by dividing inhibition of a compound to a glutamate control response; <sup>b</sup> n = number of recorded cells.

### 2.7. Antioxidant Properties

The compound **6** was assessed for its antioxidant properties *via* the DPPH (diphenyl-1-picrylhydrazyl) free radical assay. The rationale for determining the antioxidant properties in **6** stems from the fact that trolox, a water-soluble analog of vitamin E, is a well-known free radical scavenging pharmacophore [39]. Antioxidant activity was quantified as EC_50_, representing the concentration of the compound necessary to induce a 50% reduction in DPPH activity, with trolox serving as the standard. More detailed data are presented in Supplementary Information (Table S4). **6** exhibited limited radical scavenging capabilities, as evidenced by its high EC_50_ value of 114 µM, being sevenfold less potent than trolox (16 µM). This reduction in activity was an expected result, aligning with observations from prior studies involving THA-trolox heterodimers [26], where derivatives characterized by a short carbon linker (consisting of 2 to 3 methylene units) between the THA and trolox moieties demonstrated diminished antioxidant capacity. Nonetheless, THA-trolox still preserved activity against both ChEs.

### 2.8. Anti-amyloid Properties

**7c** and **7m** were tested to determine their inhibitory activities against Aβ_1-42_ self-induced aggregation. The selection of the compounds was dictated by the fact that both compounds are endowed with benzothiazole moieties, known to exert anti-amyloid properties via specific binding to Aβ [20,40]. The results for these selected compounds, alongside doxycycline as a positive control, are presented in Supplementary Information (Table S5 and Figure S3). Contrary to our expectations, neither of the chosen derivatives displayed anti-amyloid potency compared to doxycycline.

## 3. Conclusions

The primary goal of the study was to broaden the pharmacological profile of amiridine, a known cholinesterase inhibitor, by tethering to moieties of various pharmacological properties. The final derivatives, denoted as **5**–**7**, were synthesized to investigate the potential versatility of amiridine as an AChE-inhibition pharmacophore and to broaden the armamentarium of amiridine-based hybrid compounds with diverse biological activities. Structural scaffolds, including 1-adamantylamine, memantine, trolox, and various substituted/unsubstituted benzothiazoles, were introduced to augment amiridine’s AChE inhibition properties while conferring pharmacological properties against NMDA receptors, antioxidant capacity, and anti-amyloid characteristics. Comprehensive characterization of these novel compounds encompassed cytotoxicity assessments on both undifferentiated and differentiated SH-SY5Y cells, as well as prediction of BBB permeation *in silico* and experimentally via PAMPA assay.

Among the synthesized derivatives, **5c-d**, **7c**, and **7g** exhibited inhibitory activity against BChE in the sub-micromolar to low micromolar range. The most pronounced activities were determined for **5c-d** featured 1-adamantylamine and memantine scaffolds, respectively. It is pertinent to highlight that the IC_50_ values for BChE inhibition and cytotoxicity against SH-SY5Y cells for these compounds exhibited a substantial three-order difference, indicative of their low-toxic profiles. Moreover, **5c-d** demonstrated high BBB permeability scores, as indicated by the SwissADME prediction tool. Additionally, these compounds displayed promising predictions for GIA with no violations of Lipinski’s rule of five. Last but not least, the compounds successfully passed the PAINS filter. The assessment of their NMDAR inhibition potential warrants further investigation.

Further studies should focus on **5c-d** to even improve their BBB penetration and enhance their solubility.

Regrettably, **7c** and **7g**, possessing substituted benzothiazole fragments, exhibited relatively high cytotoxicity against undifferentiated SH-SY5Y cells, with IC_50_ values in the same range as determined for BChE. Contrary to the expectations, **7c** failed to demonstrate any inhibitory effect on Aβ_1-42_ self-induced aggregation.

In summary, our findings underscore the potential of the amiridine scaffold as a valuable pharmacophore for the development of novel therapeutics with a primary intention to treat AD. The synthesized compounds, particularly **5c-d**, represent promising lead structures for AD treatment. Nevertheless, further structural refinements are needed, with a focus on adjusting physicochemical properties, such as solubility and BBB permeability, to facilitate their progression in drug development endeavors.

## 4. Materials and Methods

### 4.1. Synthesis of amiridine-based heterodimers

#### 4.1.1. General Chemistry Methods

All reagents and solvents were purchased from Sigma-Aldrich (Prague, Czech Republic) or Fluorochem - Doug Discovery (United Kingdom) and were used in the highest available purity without further purification. The reactions were monitored by thin layer chromatography (TLC) performed on aluminium sheets precoated with silica gel 60 F254 (Merck, Prague, Czech Republic). The spots were visualized by ultraviolet light (at wavelength 254 nm). If necessary, the spots on TLCs were visualized using a solution of phosphomolybdic acid. Purifications of crude products were performed by column chromatography on silica gel 100 (particle size 0.063–0.200 mm, 70–230 mesh ASTM, Fluka, Prague, Czech Republic). Nuclear magnetic resonance (NMR) spectra were recorded in deuterated chloroform (CDCl_3_), methanol (CD_3_OD) or dimethyl sulfoxide (DMSO-d_6_) on a Varian S500 spectrometer operating at 500 MHz for ^1^H and 126 MHz for ^13^C. Chemical shifts (*δ*) are reported in parts per millions (ppm) and spin multiplicities are given as singlet (s), doublet (d), triplet (t), quintet or pentet (p), doublet of doublets (dd), quartet of triplets (qt), doublet of doublet of triplets (ddt), doublet or triplet of doublets (dtd), or multiplet (m). Coupling constants (J) are reported in Hz. The synthesized compounds were analyzed by LC-MS system consisting of UHLPC Dionex Ultimate 3000 coupled with Q Exactive Plus Orbital mass spectrometer to obtain high-resolution mass spectra (Thermo Fisher Scientific, Waltham, MA, USA). The purity of the final compounds established by LC-UV (254 nm) analysis was higher than 95%. Melting points were measured using a M-565 automated melting point recorder (Büchi, Flawil, Switzerland).

#### 4.1.2. Synthesis of intermediates 8, 11 – 13

Derivatives **8**, **11** and **12** were prepared by a slightly modified method adapted from [17]. ^1^H and ^13^C NMR spectra were in good agreement with previously published data [17,41]. The procedure for preparation and NMR spectra can be found in Supplementary Information.

*2-Amino-N-{1H,2H,3H,5H,6H,7H,8H-cyclopenta[b]quinolin-9-yl}acetamide* (**13**): To a suspension of **8** (1.52 g, 5.74 mmol) in dry CH_3_CN was added potassium phthalimide (1.6 g, 8.6 mmol). The mixture was stirred at reflux for 3 h, then excess of NH_2_NH_2_.H_2_O (3 mL) was added. The resulting mixture was stirred at reflux overnight. The reaction was monitored by TLC with mobile phase DCM/MeOH/NH_3_ (20/1/0.1). After cooling to room temperature the precipitate was filtered off and solvent was evaporated under reduced pressure. Crude product was purified by column chromatography. **13** was isolated as light brown solid (803 mg, 57%), decomposition from 150 °C. ^1^H NMR (CDCl_3_): *δ* 9.07 (s, 1H), 3.46 (s, 2H), 2.91 (t, *J* = 7.6 Hz, 2H), 2.83 (t, *J* = 6.3 Hz, 2H), 2.77 (t, *J* = 7.4 Hz, 2H), 2.53 (t, *J* = 6.2 Hz, 2H), 2.00 (p, *J* = 7.6 Hz, 2H), 1.71–1.79 (m, 6H) ppm. ^13^C NMR (CDCl_3_): *δ* 170.2, 163.9, 156.0, 139.0, 129.5, 123.2, 45.0, 34.5, 32.8, 30.2, 24.5, 23.1, 22.8, 22.6 ppm. HRMS [M+H]^+^: 246.15929 (calculated for [C_14_H_20_N_3_O]^+^: 246.16009).

#### 4.1.3. Synthesis of Final Derivatives

##### General procedure A

To a solution of corresponding chloroacetamine derivative **8**, **11** or **12** (1 eq) in dry CH_3_CN (0.05 M) was added amine **9**, **10** or amiridine (1.1 eq), then K_2_CO_3_ (12 eq) and KI (2.8 eq). The resulting mixture was stirred under reflux. The reaction was monitored by TLC with mobile phases DCM/MeOH/NH_3_ (20/1/0.1) for **5a-b**, PE/EA (5/1) for **5c-d** (the spots on TLCs of **11** and **12** were visualized using a solution of phosphomolybdic acid). After cooling to room temperature, the solid part was filtered of and the filtrate was concentrated under reduced pressure. Crude product was purified by column chromatography.

##### General procedure B

**13** (1 eq) and corresponding 2-chlorobenzothiazole (1 eq) was put into 10 mL flask and DIPEA (9 eq) was added. The resulting suspense was stirred at 100 °C overnight. The reaction was monitored by TLC with mobile phase DCM/MeOH/NH_3_ (20/1/0.1). After cooling to room temperature the mixture was filtrated through glass frit filter and the precipitate was washed with DCM and MeOH.

*2-[(adamantan-1-yl)amino]-N-{1H,2H,3H,5H,6H,7H,8H-cyclopenta[b]napthtalen-4-yl}acetamide* (**5a**): Prepared from **8** and **9** by general procedure **A**. **5a** was isolated as white solid (51 mg, 54%), decomposition from 155 °C. ^1^H NMR (CD_3_OD): *δ* 3.44–3.43 (m, 2H), 2.93 (t, *J* = 7.6 Hz, 2H), 2.87–2.84 (m, 2H), 2.81 (t, *J* = 7.5 Hz, 2H), 2.64 (t, *J* = 6.2 Hz, 2H), 2.12–2.06 (m, 5H), 1.88–1.79 (m, 4H), 1.74–1.65 (m, 12H) ppm. ^13^C NMR (CD_3_OD): *δ* 171.9, 163.0, 155.3, 140.2, 130.9, 124.9, 50.5, 43.2, 41.9, 36.2, 33.5, 31.7, 29.6, 29.4, 24.1, 22.6, 22.4, 22.2 ppm. HRMS [M+H]^+^: 380.26920 (calculated for [C_24_H_34_N_3_O]^+^: 380.26964).

*N-{1H,2H,3H,5H,6H,7H,8H-cyclopenta[b]napthalen-4-yl}-2-[(3,5-dimethyladamantan-1-yl)amino]acetamide* (**5b**): Prepared from **8** and **10** by general procedure **A**. **5b** was isolated as colorless oil (45 mg, 54%). ^1^H NMR (CD_3_OD): *δ* 3.43 (m, 2H), 2.94 (t, *J* = 7.8 Hz, 2H), 2.86 (t, *J* = 6.4 Hz, 2H), 2.81 (t, *J* = 7.5 Hz, 2H), 2.64 (t, *J* = 6.3 Hz, 2H), 2.18–2.12 (m, 1H), 2.11–2.06 (m, 2H), 1.89–1.78 (m, 4H), 1.54–1.53 (m, 2H), 1.38–1.29 (m, 8H), 1.19–1.11 (m, 2H), 0.87 (s, 6H) ppm. ^13^C NMR (CD_3_OD): *δ* 171.9, 163.0, 155.3, 140.2, 131.0, 125.0, 60.1, 52.1, 50.5, 48.1, 43.4, 42.5, 40.4, 33.5, 32.0, 31.7, 30.3, 29.4, 24.1, 22.5, 22.4, 22.2 ppm. HRMS [M]^+^: 408.30029 (calculated for [C_26_H_38_N_3_O]^+^: 408.30094).

*N-(adamantan-1-yl)-2-({1H,2H,3H,5H,6H,7H,8H-cyclopenta[b]naphthalen-4-yl}amino)acetamide* (**5c**): Prepared from **11** and amiridine by general procedure **A** (51 mg, 48%). **5c** was isolated as light brown solid, after recrystallization from a mixture of solvents (DCM, MeOH) slightly yellow solid, decomposition from 260 °C. ^1^H NMR (CD_3_OD): *δ* 4.83 (s, 2H), 3.08 (t, *J* = 7.7 Hz, 2H), 2.89 (t, *J* = 7.6 Hz, 2H), 2.73 (t, *J* = 6.2 Hz, 2H), 2.51 (t, *J* = 6.2 Hz, 2H), 2.25 (p, *J* = 7.7 Hz, 2H), 2.08–2.04 (m, 9H), 1.92–1.82 (m, 4H), 1.76–1.69 (m, 6H) ppm. ^13^C NMR (CD_3_OD): *δ* 164.8, 156.2, 154.4, 148.7, 119.9, 116.3, 52.3, 52.2, 40.9, 36.0, 32.4, 29.5, 28.0, 26.5, 22.9, 21.3, 21.0, 20.4 ppm. HRMS [M+H]^+^: 380.26877 (calculated for [C_24_H_34_N_3_O]^+^: 380.26964).

*2-({1H,2H,3H,5H,6H,7H,8H-cyclopenta[b]naphthalen-4-yl}amino)-N-(3,5-dimethyladamantan-1-yl)acetamide* (**5d**): Prepared from **12** and amiridine by general procedure **A** (48 mg, 41%). **5d** was isolated as light brown solid, after recrystallization from a mixture of solvents (DCM, MeOH) slightly yellow solid, decomposition from 266 °C. ^1^H NMR (CD_3_OD): *δ* 4.82–4.81 (m, 2H), 3.07 (t, *J* = 7.7 Hz, 2H), 2.89 (t, *J* = 7.6 Hz, 2H), 2.72 (t, *J* = 6.3 Hz, 2H), 2.51 (t, *J* = 6.2 Hz, 2H), 2.25 (p, *J* = 7.7 Hz, 2H), 2.15 (p, *J* = 3.2 Hz, 1H), 1.92–1.83 (m, 6H), 1.70–1.63 (m, 4H), 1.42–1.29 (m, 4H), 1.17 (s, 2H), 0.86 (s, 6H) ppm. ^13^C NMR (CD_3_OD): *δ* 164.8, 156.1, 154.4, 148.7, 119.9, 116.3, 53.9, 52.1, 52.2, 46.8, 42.2, 39.3, 32.3, 31.9, 30.1, 29.2, 27.9, 26.5, 22.9, 21.3, 21.0, 20.4 ppm. HRMS [M+H]^+^: 408.30020 (calculated for [C_26_H_38_N_3_O]^+^: 408.30094).

*N-{1H,2H,3H,5H,6H,7H,8H-cyclopenta[b]quinolin-9-yl}-2-[(6-hydroxy-2,5,7,8-tetramethyl-3,4-dihydro-2H-1-benzopyran-2-yl)formamido]acetamide* (**6**): Trolox (77 mg, 0.31 mmol) was dissolved in dry DMF (1 mL) and TEA (128 µL, 0.92 mmol) was added. After 30 min BOP (135 mg, 0.31 mmol) was added to the mixture and after additional 1 h a solution of **13** (75 mg, 0.31 mmol) was also added. The resulting mixture was stirred at room temperature for 2 days. The reaction was monitored by TLC with mobile phase DCM/MeOH/NH_3_ (20/1/0.1). After completion of the reaction the mixture was diluted with DCM (15 mL) and washed with water (15 mL). Water was then washed with DCM (15 mL) and joined organic parts were concentrated under reduced pressure. Crude product was purified by column chromatography. **6** was isolated as light yellow amorphous solid (125 mg, 86%). ^1^H NMR (CD_3_OD): *δ* 4.10–4.07 (m, 1H), 4.03 – 3.99 (m, 1H), 2.99 (t, *J* = 7.7 Hz, 2H), 2.88 (t, *J* = 6.2 Hz, 2H), 2.73 (t, *J* = 7.5 Hz, 2H), 2.57–2.49 (m, 4H), 2.20 (s, 3H), 2.14–2.10 (m, 5H), 1.95 (s, 3H), 1.90–1.84 (m, 2H), 1.82–1.76 (m, 2H), 1.54 (s, 3H), 1.30 (m, 2H) ppm. ^13^C NMR (CD_3_OD): *δ* 176.1, 167.7, 161.8, 154.2, 146.0, 144.2, 132.2, 126.1, 123.2, 121.8, 120.5, 117.2, 77.8, 46.5, 42.2, 35.6, 35.6, 32.9, 30.7, 23.7, 23.6, 22.6, 21.9, 20.2, 11.4, 10.8, 10.3 ppm. HRMS [M+H]^+^: 478.26965 (calculated for [C_28_H_36_N_3_O_4_]^+^: 478.27003).

*2-[(1,3-nebothiazol-2-yl)amino]-N-{1H,2H,3H,5H,6H,7H,8H-cyclopenta[b]quinolin-9-yl}acetamide* (**7a**): **13** (107 mg, 0.44 mmol) and 2-chlorobenzothiazole (57 µL, 0.44 mmol) were stirred at 110 °C overnight. After cooling to room temperature the crude product was purified by column chromatography with mobile phase DCM/MeOH/NH_3_ (20/1/0.1). **7a** was isolated as light yellow solid (53 mg, 32%), decomposition from 191 °C. ^1^H NMR (CD_3_OD): *δ* 7.63 (d, *J* = 7.8 Hz, 1H) 7.48 (d, *J* = 8.1 Hz, 1H), 7.30–7.27 (m, 1H), 7.10 (t, *J* = 7.5 Hz, 1H), 4.31 (s, 2H). 2.92 (t, *J* = 7.6 Hz, 2H), 2.85–2.80 (m, 4H), 2.63 (t, *J* = 6.5 Hz, 2H), 2.07 (p, *J* = 7.2 Hz, 2H), 1.86–1.71 (m, 2H), 1.76–1.71 (m, 2H) ppm. ^13^C NMR (CD_3_OD): *δ* 168.7, 167.3, 162.9, 155.3, 151.7, 140.3, 131.6, 130.6, 125.7, 125.5, 121.7, 120.6, 118.2, 46.8, 33.4, 31.7, 29.1, 23.9, 22.5, 22.4, 22.1 ppm. HRMS [M+H]^+^: 379.15793 (calculated for [C_21_H_23_N_4_OS]^+^: 379.15871).

*2-[(6-bromo-1,3-benzothiazol-2-yl)amino]-N-{1H,2H,3H,5H,6H,7H,8H-cyclopenta[b]quinolin-9-yl}acetamide* (**7b**): Prepared from **13** and 6-bromo-2-chlorobenzothiazole by general method **B**. **7b** was isolated as slightly brown solid (45 mg, 77%), decomposition from 244 °C. ^1^H NMR (DMSO-d_6_): *δ* 9.69 (s, 1H), 8.56 (t, *J* = 5.8 Hz, 1H), 7.95 (d, *J* = 2.0 Hz, 1H), 7.38 (dd, *J* = 8.5, 2.1 Hz, 1H), 7.29 (d, *J* = 8.5 Hz, 1H), 4.24 (d, *J* = 5.5 Hz, 2H), 2.82 (t, *J* = 7.6 Hz, 2H), 2.76 (t, *J* = 6.4 Hz, 2H), 2.69 (t, *J* = 7.5 Hz, 2H), 2.59 (t, *J* = 6.4 Hz, 2H), 1.95 (p, *J* = 7.6 Hz, 2H), 1.77–1.72 (m, 2H), 1.70–1.65 (m, 2H) ppm. ^13^C NMR (DMSO-d_6_): *δ* 167.4, 167.2, 163.0, 155.6, 151.9, 140.3, 133.5, 130.7, 128.9, 124.9, 124.0, 119.9, 113.0, 47.2, 34.1, 32.6, 29.7, 24.3, 23.0, 23.0, 22.7 ppm. HRMS [M+H]^+^: 457.07550/459.07333 (calculated for [C_21_H_22_BrN_4_OS]^+^: 457.06922).

*2-[(5-bromo-1,3-benzothiazol-2-yl)amino]-N-{1H,2H,3H,5H,6H,7H,8H-cyclopenta[b]quinolin-9-yl}acetamide* (**7c**): Prepared from **13** and 5-bromo-2-chlorobenzothiazole by general method **B**. **7c** was isolated as light brown solid (25 mg, 21%), decomposition from 224 °C. ^1^H NMR (DMSO-d_6_): *δ* 9.69 (s, 1H), 8.64 (t, *J* = 5.7 Hz, 1H), 7.67 (d, *J* = 8.3 Hz, 1H), 7.52 (d, *J* = 1.9 Hz, 1H), 7.20 (dd, *J* = 8.4, 2.0 Hz, 1H), 4.25 (d, *J* = 5.4 Hz, 2H), 2.83 (t, *J* = 7.6 Hz, 2H), 2.77 (t, *J* = 6.4 Hz, 2H), 2.70 (t, *J* = 7.4 Hz, 2H), 2.60 (t, *J* = 6.3 Hz, 2H), 1.96 (p, *J* = 7.5 Hz, 2H), 1.78–1.74 (m, 2H), 1.71–1.66 (m, 2H) ppm. ^13^C NMR (DMSO-d_6_): *δ* 168.1, 167.3, 163.1, 155.7, 154.2, 140.1, 130.6, 129.3, 124.8, 124.0, 123.2, 120.8, 118.8, 47.2, 34.2, 32.7, 29.6, 24.4, 23.0, 23.0, 22.7 ppm. HRMS [M+H]^+^: 457.06915/459.06686 (calculated for [C_21_H_22_BrN_4_OS]^+^: 457.06922).

*2-[(4-bromo-1,3-benzothiazol-2-yl)amino]-N-{1H,2H,3H,5H,6H,7H,8H-cyclopenta[b]quinolin-9-yl}acetamide* (**7d**): Prepared from **13** and 2-Chloro-4-bromobenzothiazole by general method **B**. **7d** was isolated as slightly brown solid (44 mg, 47%), decomposition from 255 °C. ^1^H NMR (DMSO-d_6_): *δ* 9.71 (s, 1H), 8.68 (t, *J* = 5.7 Hz, 1H), 7.72 (dd, *J* = 7.8, 1.1 Hz, 1H), 7.46 (dd, *J* = 7.8, 1.1 Hz, 1H), 6.96 (t, *J* = 7.8 Hz, 1H), 4.28 (d, *J* = 5.4 Hz, 2H), 2.83 (t, *J* = 7.6 Hz, 2H), 2.77 (t, *J* = 6.3 Hz, 2H), 2.72 (t, *J* = 7.5 Hz, 2H), 2.62 (t, *J* = 6.3 Hz, 2H), 1.96 (p, *J* = 7.5 Hz, 2H), 1.78–1.73 (m, 2H), 1.71–1.65 (m, 2H) ppm. ^13^C NMR (DMSO-d_6_): *δ* 167.1, 166.9, 163.1, 155.7, 150.6, 140.1, 132.0, 130.6, 129.2, 124.8, 122.8, 121.0, 111.3, 47.1, 34.2, 32.8, 29.7, 24.4, 23.1, 23.0, 22.7 ppm. HRMS [M+H]^+^: 457.07550/459.06631 (calculated for [C_21_H_22_BrN_4_OS]^+^: 457.06922).

*2-[(5-cyano-1,3-benzothiazol-2-yl)amino]-N-{1H,2H,3H,5H,6H,7H,8H-cyclopenta[b]quinolin-9-yl}acetamide* (**7e**): Prepared from **13** and 2-chlorobenzothiazole-5-carbonitrile by general method **B**. **7e** was isolated as slightly yellow solid (25 mg, 21%), decomposition from 234 °C. ^1^H NMR (DMSO-d_6_): *δ* 9.71 (s, 1H), 8.78 (t, *J* = 5.7 Hz, 1H), 7.94 (d, *J* = 8.0 Hz, 1H), 7.76 (d, *J* = 1.6 Hz, 1H), 7.45 (dd, *J* = 8.1, 1.7 Hz, 1H), 4.30 (d, *J* = 5.1 Hz, 2H), 2.83 (t, *J* = 7.6 Hz, 2H), 2.77 (t, *J* = 6.4 Hz, 2H), 2.70 (t, *J* = 7.5 Hz, 2H), 2.59 (t, *J* = 6.2 Hz, 2H), 1.96 (p, *J* = 7.8 Hz, 2H), 1.78–1.74 (m, 2H), 1.71–1.66 (m, 2H) ppm. ^13^C NMR (DMSO-d_6_): *δ* 168.2, 167.1, 163.1, 155.7, 152.8, 124.8, 124.7, 122.9, 121.2, 119.9, 108.5, 47.2, 34.2, 32.7, 29.7, 24.3, 23.0, 23.0, 22.7 ppm. HRMS [M+H]^+^: 404.15320 (calculated for [C_22_H_22_N_5_OS]^+^: 404.15396).

*2-[(6-chloro-1,3-benzothiazol-2-yl)amino]-N-{1H,2H,3H,5H,6H,7H,8H-cyclopenta[b]quinolin-9-yl}acetamide* (**7f**): Prepared from **13** and 2,6-dichlorobenzothiazole by general method **B**. **7f** was isolated as slightly brown solid (43 mg, 32%), decomposition from 231 °C. ^1^H NMR (DMSO-d_6_): *δ* 9.68 (s, 1H), 8.55 (t, *J* = 5.8 Hz, 1H), 7.83 (d, *J* = 2.3 Hz, 1H), 7.35 (d, *J* = 8.5 Hz, 1H), 7.27 (dd, *J* = 8.6, 2.3 Hz, 1H), 4.24 (d, *J* = 5.3 Hz, 2H), 2.82 (t, *J* = 7.6 Hz, 2H), 2.76 (t, *J* = 6.4 Hz, 2H), 2.70 (t, *J* = 7.4 Hz, 2H), 2.59 (t, *J* = 6.3 Hz, 2H), 1.96 (p, *J* = 7.6 Hz, 2H), 1.78–1.73 (m, 2H), 1.70–1.65 (m, 2H) ppm. ^13^C NMR (DMSO-d_6_): *δ* 167.4, 167.2, 163.1, 155.7, 151.6, 140.1, 133.0, 130.6, 126.2, 125.4, 124.8, 121.2, 119.4, 47.2, 34.2, 32.7, 29.6, 24.3, 23.0, 23.0, 22.7 ppm. HRMS [M+H]^+^: 413.12897 (calculated for [C_22_H_22_ClN_4_OS]^+^: 413.11973).

*2-[(5-chloro-1,3-benzothiazol-2-yl)amino]-N-{1H,2H,3H,5H,6H,7H,8H-cyclopenta[b]quinolin-9-yl}acetamide* (**7g**): Prepared from **13** and 2,5-dichlorobenzothiazole by general method **B**. **7g** was isolated as slightly brown solid (43 mg, 41%), decomposition from 232 °C. ^1^H NMR (DMSO-d_6_): *δ* 9.68 (s, 1H), 8.63 (t, *J* = 5.7 Hz, 1H), 7.73 (d, *J* = 8.4 Hz, 1H), 7.39 (d, *J* = 2.1 Hz, 1H), 7.08 (dd, *J* = 8.3, 2.1 Hz, 1H), 4.26 (d, *J* = 5.4 Hz, 2H), 2.83 (t, *J* = 7.6 Hz, 2H), 2.77 (t, *J* = 6.4 Hz, 2H), 2.70 (t, *J* = 7.5 Hz, 2H), 2.60 (t, *J* = 6.3 Hz, 2H), 1.96 (p, *J* = 7.5 Hz, 2H), 1.78–1.74 (m, 2H), 1.71–1.66 (m, 2H) ppm. ^13^C NMR (DMSO-d_6_): *δ* 168.3, 167.3, 163.1, 155.7, 153.9, 140.1, 130.7, 130.6, 130.1, 124.8, 122.8, 121.3, 117.9, 47.2, 34.2, 32.7, 29.6, 24.3, 23.0, 23.0, 22.7 ppm. HRMS [M+H]^+^: 413.11346 (calculated for [C_22_H_22_ClN_4_OS]^+^: 413.11973).

*2-[(4-chloro-1,3-benzothiazol-2-yl)amino]-N-{1H,2H,3H,5H,6H,7H,8H-cyclopenta[b]quinolin-9-yl}acetamide* (**7h**): Prepared from **13** and 2,4-dichlorobenzothiazole by general method **B**. **7h** was isolated as brown solid (35 mg, 34%), decomposition from 230 °C. ^1^H NMR (DMSO-d_6_): *δ* 9.72 (s, 1H), 8.69 (t, *J* = 5.7 Hz, 1H), 7.68 (dd, *J* = 7.8, 1.1 Hz, 1H), 7.32 (dd, *J* = 7.9, 1.1 Hz, 1H), 7.03 (t, *J* = 7.9 Hz, 1H), 4.29 (d, *J* = 5.6 Hz, 2H), 2.83 (t, *J* = 7.6 Hz, 2H), 2.77 (t, *J* = 6.3 Hz, 2H), 2.72 (t, *J* = 7.4 Hz, 2H), 2.62 (t, *J* = 6.3 Hz, 2H), 1.96 (p, *J* = 7.6 Hz, 2H), 1.78–1.73 (m, 2H), 1.71–1.65 (m, 2H) ppm. ^13^C NMR (DMSO-d_6_): *δ* 167.2, 163.0, 155.6, 149.3, 140.2, 132.6, 130.6, 126.2, 124.8, 122.3, 122.3, 120.5, 47.2, 34.2, 32.7, 29.7, 24.4, 23.0, 23.0, 22.7 ppm. HRMS [M+H]^+^: 413.11853 (calculated for [C_22_H_22_ClN_4_OS]^+^: 413.11973).

*N-{1H,2H,3H,5H,6H,7H,8H-cyclopenta[b]quinolin-9-yl}-2-[(6-nitro-1,3-benzothiazol-2-yl)amino]acetamide* (**7i**): Prepared from **13** and 2-chloro-6-nitro-benzothiazole by general method **B**. **7i** was isolated as light brown solid (33 mg, 34%), decomposition 235 °C. ^1^H NMR (DMSO-d_6_): *δ* 9.74 (s, 1H) 9.12 (s, 1H), 8.75 (d, *J* = 2.5 Hz, 1H), 8.14 (dd, *J* = 8.9, 2.5 Hz, 1H), 7.48 (d, *J* = 8.9 Hz, 1H), 4.34 (s, 2H), 2.83 (t, *J* = 7.6 Hz, 2H), 2.77 (t, *J* = 6.4 Hz, 2H), 2.70 (t, *J* = 7.4 Hz, 2H), 2.60 (t, *J* = 6.1 Hz, 2H), 1.96 (p, *J* = 8.0 Hz, 2H), 1.78–1.73 (m, 2H), 1.71–1.67 (m, 2H) ppm. ^13^C NMR (DMSO-d_6_): *δ* 171.4, 166.9, 163.1, 158.2, 155.8, 141.5, 140.0, 132.0, 130.6, 124.8, 122.6, 118.4, 117.8, 47.3, 34.2, 32.7, 29.6, 24.4, 23.0, 23.0, 22.7 ppm. HRMS [M+H]^+^: 424.14294 (calculated for [C_21_H_22_N_5_O_3_S]^+^: 424.14379).

*N-{1H,2H,3H,5H,6H,7H,8H-cyclopenta[b]quinolin-9-yl}-2-[(5-nitro-1,3-benzothiazol-2-yl)amino]acetamide* (**7j**): Prepared from **13** and 2-chloro-5-nitro-1,3-benzothiazole by general method **B**. **7j** was isolated as light yellow solid (53 mg, 49%), decomposition from 231 °C. ^1^H NMR (DMSO-d_6_): *δ* 9.75 (s, 1H), 8.88 (t, *J* = 5.7 Hz, 1H), 8.08 (d, *J* = 2.2. Hz, 1H), 8.01–7.99 (m, 1H), 7.94–7.91 (m, 1H), 4.32–4.31 (m, 2H), 8.83 (t, *J* = 7.6 Hz, 2H), 2.77 (t, *J* = 6.4 Hz, 2H), 2.71 (t, *J* = 7.4 Hz, 2H), 2.63 (t, *J* = 6.4 Hz, 2H), 1.97 (p, *J* = 7.3 Hz, 2H), 1.78–1.74 (m, 2H), 1.72–1.67 (m, 2H) ppm. ^13^C NMR (DMSO-d_6_): *δ* 169.0, 167.1, 163.1, 155.7, 153.0, 146.6, 140.1, 139.5, 130.6, 124.8, 122.3, 116.3, 112.2, 47.2, 34.2, 32.7, 29.6, 24.4, 23.0, 23.0, 22.7 ppm. HRMS [M+H]^+^: 424.14294 (calculated for [C_21_H_22_N_5_O_3_S]^+^: 424.14379).

*N-{1H,2H,3H,5H,6H,7H,8H-cyclopenta[b]quinolin-9-yl}-2-[(7-nitro-1,3-benzothiazol-2-yl)amino]acetamide* (**7k**): Prepared from **13** and 2-chloro-7-nitrobenzo[d]thiazole by general method **B**. **7k** was isolated as yellow solid (50 mg, 48%), decomposition from 231 °C. ^1^H NMR (DMSO-d_6_): *δ* 9.73 (s, 1H), 8.89 (d, *J* = 5.8 Hz, 1H), 8.03 (dd, *J* = 8.2, 0.9 Hz, 1H), 7.79 (dd, *J* = 7.8, 1.0 Hz, 1H), 7.54 (t, *J* = 8.1 Hz, 1H), 4.32 (d, *J* = 4.0 Hz, 2H), 2.83 (t, *J* = 7.6 Hz, 2H), 2.77 (t, *J* = 6.4 Hz, 2H), 2.71 (t, *J* = 7.4 Hz, 2H), 2.60 (t, *J* = 6.1 Hz, 2H), 1.96 (p, *J* = 7.5 Hz, 2H), 1.78–1.73 (m, 2H), 1.71–1.66 (m, 2H) ppm. ^13^C NMR (DMSO-d_6_): *δ* 169.2, 167.1, 163.1, 155.7, 155.1, 142.1, 140.1, 130.6, 127.5, 126.9, 124.5, 117.4, 47.2, 34.2, 32.7, 29.6, 24.4, 23.0, 23.0, 22.7 ppm. HRMS [M+H]^+^: 424.14337 (calculated for [C_21_H_22_N_5_O_3_S]^+^: 424.14379).

*N-{1H,2H,3H,5H,6H,7H,8H-cyclopenta[b]quinolin-9-yl}-2-{[6-(methylsulfanyl)-1,3-benzothiazol-2-yl]a mino}acetamide* (**7l**): Prepared from **13** and 2-chloro-6-(methylthio)benzo[d]thiazole by general method **B**. **7l** was isolated as light brown solid (22 mg, 25%), decomposition from 180 °C. ^1^H NMR (DMSO-d_6_): *δ* 9.71 (s, 1H), 8.48 (d, *J* = 5.8 Hz, 1H), 7.74 (d, *J* = 1.9 Hz, 1H), 7.37 (d, *J* = 8.4 Hz, 1H), 7.25 (dd, *J* = 8.4, 2.0 Hz, 1H), 4.28 (d, *J* = 5.6 Hz, 2H), 2.88 (t, *J* = 7.6 Hz, 2H), 2.82 (t, *J* = 6.4 Hz, 2H), 2.74– 2.77 (m, 5H), 2.65 (t, *J* = 6.4 Hz, 2H), 2.01 (p, *J* = 7.4 Hz, 2H), 1.84–1.78 (m, 2H), 1.76–1.71 (m, 2H) ppm. ^13^C NMR (DMSO-d_6_): *δ* 167.6, 166.4, 163.1, 155.7, 150.8, 140.2, 132.4, 130.6, 130.0, 125.9, 124.8, 120.3, 118.8 ppm. HRMS [M+H]^+^: 425.14621 (calculated for [C_22_H_25_N_4_OS_2_]^+^: 425.14643).

*N-{1H,2H,3H,5H,6H,7H,8H-cyclopenta[b]quinolin-9-yl}-2-{[6-(trifluoromethoxy)-1,3-benzothiazol-2-yl] amino}acetamide* (**7m**): Prepared from **13** and 2-chrolo-6-(trifuoromethoxy)benzo[d]thiazole by general method **B**. **7m** was isolated as white solid (52 mg, 34%), decomposition from 270 °C. ^1^H NMR (DMSO-d_6_): *δ* 9.68 (s, 1H), 8.60 (t, *J* = 5.8 Hz, 1H), 7.84 (d, *J* = 2.5 Hz, 1H), 7.43 (d, *J* = 8.7 Hz, 1H), 7.23 (dd, *J* = 8.7, 2.6 Hz, 1H), 4.25 (d, *J* = 5.5 Hz, 2H), 2.82 (t, *J* = 7.6 Hz, 2H), 2.76 (t, *J* = 6.4 Hz, 2H), 2.70 (t, *J* = 7.4 Hz, 2H), 2.59 (t, *J* = 6.2 Hz, 2H), 1.96 (t, *J* = 7.6 Hz, 2H), 1.78–1.73 (p, *J* = 7.6 Hz, 2H), 1.70–1.66 (m, 2H) ppm. ^13^C NMR (DMSO-d_6_): *δ* 167.8, 167.4, 163.1, 155.7, 151.8, 142.8, 140.1, 132.4, 130.6, 124.8, 121.8, 119.6, 118.8, 115.1, 47.2, 34.2, 32.7, 29.6, 24.3, 23.0, 23.0, 22.7 ppm. HRMS [M+H]^+^: 463.14050 (calculated for [C_22_H_22_F_3_N_4_O_2_S]^+^: 463.14101).

### 4.2. Cholinesterase Inhibition Assay

The inhibitory activities of **5**–**7** and all the reference compounds were determined using modified Ellman’s method [25,42,43] against human recombinant AChE (*h*AChE, E.C. 3.1.1.7, purchased from Sigma-Aldrich, Prague, Czech Republic) and human plasmatic BChE (*h*BChE, E.C. 3.1.1.8, purchased from Sigma-Aldrich, Prague, Czech Republic). The results are expressed as IC_50_ (the concentration of the compound that is required to reduce 50% of cholinesterase activity). Other used compounds – phosphate buffer solution (PBS, pH = 7.4), 5,5’-dithio-bis(2-nitrobenzoic) acid (Ellman’s reagent, DTNB), acetylthiocholine (ATCh) and butyrylthiocholine (BTCh) were also commercially available and were purchased from Sigma-Aldrich (Prague, Czech Republic). During the measurement, 96-well microplates from polystyrene (ThermoFisher Scientific, Waltham, MA, USA) were used. The solutions of corresponding enzyme in PBS were prepared up to final activity 0.002 U/µL. The assay medium (100 µL) consisted of cholinesterase (10 µL), DTNB (20 µL of 0.01 M solution) and PBS (40 µL of 0.1 M solution). The solutions of the tested compounds (10 µL of different concentrations) were pre-incubated for 5 minutes in the assay medium and then solution of substrate (20 µL of 0.01 M ATCh or BTCh iodide solution) was added to initiate the reaction. The increase of absorbance was measured at 412 nm using Multimode microplate reader Synergy 2 (BioTek Inc., Winooski, VT, USA). For the calculation of the resulting measured activity (the percentage of inhibition I) following formula was used:

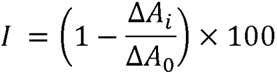

where Δ*A*_i_ indicates absorbance change provided by adequate enzyme exposed to corresponding inhibitor and Δ*A*_0_ indicates absorbance change when a solution of PBS was added instead of a solution of inhibitor. Software Microsoft Excel (Redmont, WA, USA) and GraphPad Prism version 5.02 for Windows (GraphPad Software, San Diego, CA, USA) were used for the statistical data evaluation.

### 4.3. *In silico* Studies

Molecular docking was used for binding pose calculations. The 3D structure of ligands was built by OpenBabel, v. 2.3.2 [44] and optimized by Avogadro, v. 1.2.0 using the force fields GAFF [45]. The ligands were converted into pdbqt-format by OpenBabel, v. 2.3.2. The human BChE was gained from the RCSB database (PDB ID: 4BDS, resolution 2.1 Å) and prepared for docking by the function DockPrep of the software Chimera, v. 1.14 [46] and by MGLTools, v. 1.5.4 [47]. The docking calculation was made by Vina, v. 1.1.2 as semi-flexible with flexible ligands and rigid receptor [48].

The docking pose of ligands was then improved by molecular dynamic (MD) simulation. The receptor structure was prepared using the software Chimera. The best-scored docking poses were taken as the starting point for MD. The force-field parameters for ligands were assessed by Antechamber[49] v. 20.0 using General Amber force-field 2 [50]. MD simulation was carried out by Gromacs, v. 2018.1 [51]. The complex receptor-ligand was solvated in the periodic water box using the TIP3P model. The system was neutralized by adding Na^+^ and Cl^−^ ions to a concentration of 10 nM. The system energy was minimalized and equilibrated in a 100-ps isothermal-isochoric NVT and then a 100-ps isothermal-isobaric NPT phase. Then, a 10-ns MD simulation was run at a temperature of 300 K. The molecular docking and MD results were 3D visualized by the PyMOL Molecular Graphics System, Version 2.5.2, Schrödinger, LLC.

### 4.4. Cytotoxicity evaluation

The cytotoxicity of tested compounds was assessed on the HepG2 and SH-SY5Y cells in both undifferentiated and differentiated form (ECACC, 94030304) using the MTT (Sigma-Aldrich, St. Louis, MO) as described previously[31].

The both cell lines were cultured according to ECACC recommended conditions and seeded into 96-well plates in 100 µL and density of 8000 per well. For evaluation of cytotoxicity of the differentiated form the SH-SY5Y cells also underwent 9-day differentiation protocol into mature neurons according to Riegerová *et al* . [52]. The tested compounds were dissolved in DMSO (Sigma Aldrich) or in phosphate buffer saline (PBS, Sigma Aldrich) and subsequently in the growth medium - high glucose Dulbecco’s Modified Eagle’s medium (DMEM; Sigma-Aldrich, D6429). The final concentration of DMSO did not exceed 1% (v/v). Then, the differentiated SH-SY5Y cells were treated with concentrations corresponding to IC_50_ values obtained from the cytotoxicity measurement using undifferentiated SH-SY5Y cells. Cells were exposed to a tested compound for 24 h. Then the medium was replaced by a medium containing 10 μM of MTT and cells were allowed to produce formazan for another approximately 3 h under surveillance. Thereafter, medium was aspirated and purple crystals of MTT formazan were dissolved in 100 μL DMSO under shaking. Cell viability was assessed spectrophotometrically by the amount of formazan produced. Absorbance was measured at 570 nm with 650 nm reference wavelength on Spark (Tecan Group Ltd, Switzerland). The IC_50_ value was then calculated from the control-subtracted triplicates using non-linear regression (four parameters) of GraphPad Prism 9 software. Final IC_50_ and SEM values were obtained as a mean of three independent measurements.

### 4.5. PAMPA assay

PAMPA (the parallel artificial membrane permeability assay) is a high-throughput screening tool applicable for prediction of the passive transport of potential drugs across the BBB [32,33]. In this study, it has been used as a non-cell-based *in vitro* assay carried out in a coated 96-well membrane filter. The filter membrane of the donor plate was coated with PBL (Polar Brain Lipid, Avanti, USA) in dodecane (4 µl of 20 mg/ml PBL in dodecane) and the acceptor well was filled with 300 µl of PBS (pH 7.4; ***V***_A_). The tested compounds were dissolved first in DMSO and subsequently dluted with PBS (pH 7.4) to final concentrations of 50–100 μM in the donor wells. Concentration of DMSO did not exceed 0.5% (v/v) in the donor solution. The final concentration of DMSO did not exceed 0.5% (v/v) in the donor solution. About 300 µL of the donor solution was added to the donor wells (*V*_D_) and the donor filter plate was carefully put on the acceptor plate so that coated membrane was "in touch" with both donor solution and acceptor buffer. In principle, test compound diffused from the donor well through the lipid membrane (*Area* = 0.28 cm^2^) to the acceptor well. The concentration of the drug in both donor and the acceptor wells were assessed after 3, 4, 5, and 6 h of incubation in quadruplicate using the UV plate reader Spark (Tecan Group Ltd, Switzerland) at the maximum absorption wavelength of each compound. Besides that, solution of theoretical compound concentration, simulating the equilibrium state established if the membrane were ideally permeable was prepared and assessed as well. Concentration of the compounds in the donor and acceptor well and equilibrium concentration were calculated from the standard curve and expressed as the permeability (*Pe*) according the equation:

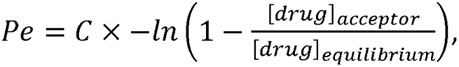

Where

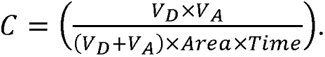

### 4.6. Assessment of NMDA Receptor Antagonistic Activity

Whole-cell patch-clamp recordings were performed using an Axopatch 200B amplifier (Molecular Devices, USA) at a holding membrane potential of -60 mV and +40 mV in the human embryonic kidney 293 (HEK293) cell line [53]. HEK293 cells were co-transfected with DNA vectors encoding human versions of the GluN1-4a (GluN1) and GluN2B subunits, together with GFP (green fluorescent protein) as described previously [54]. Electrophysiological measurements were performed ∼24-48 hours after completion of transfection. The extracellular recording solution (ECS) contained (in mM): 160 NaCl, 2.5 KCl, 10 HEPES, 10 D-glucose, 0.2 EDTA and 0.7 CaCl_2_ (pH 7.3 with NaOH). Borosilicate glass micropipettes (tip resistance 4-6 MΩ) were prepared using a P-1000 puller (Sutter Instruments, USA) and filled with an intracellular recording solution containing (in mM): 125 gluconic acid, 15 CsCl, 5 EGTA, 10 HEPES, 3 MgCl2, 0.5 CaCl_2_, and 2 ATP-Mg salts (pH 7.2 with CsOH). A microprocessor-controlled rapid perfusion system with a time constant for solution exchange around the cell of ∼20 ms was used. The co-agonist glycine (100 µM) was applied throughout the recording period, and the agonist L-glutamate was used at a concentration of 1 mM to elicit responses of the GluN1/GluN2B receptors. The test drugs were dissolved in dimethyl sulfoxide (DMSO) at a concentration of 10 mM and their inhibitory effect was tested by electrophysiology at a concentration of 30 µM. All described chemicals for electrophysiology were purchased from Merck (Czech Republic). The currents were low-pass filtered at 2 kHz using an eight-pole Bessel filter, digitized at 5 kHz and recorded using the pCLAMP 9 program (Molecular Devices, USA). The inhibitory effect of the studied compounds was analyzed at steady-state conditions using the Clampfit 10.7 program (Molecular Devices, USA).

### 4.7. Antioxidant Properties

DPPH, methanol, and trolox (as reference standard) were purchased from Sigma-Aldrich (Czech Republic). Polystyrene Nunc 96-well microplates with flat bottom shape (ThermoFisher Scientific, USA) were used for the measuring purposes. DPPH solution was prepared at 0.2 mM concentration. The assay medium (200 µL) consisted of 100 µL of DPPH solution and 100 µL of tested compound (10^-3^ – 10^-6^ M). The reaction time constituted 30 minutes. The antioxidant activity was determined by measuring the change in absorbance relative to DPPH solution at 517 nm at laboratory temperature using Multi-mode microplate reader Synergy 2 (Vermont, USA). Each concentration was tested in triplicate. Software GraphPad Prism version 5 for Windows (GraphPad Software, San Diego, CA, USA) was used for statistical data evaluation.

### 4.8. Inhibition of Amyloid-beta Formation

#### Preparation of samples

1 mg of Aβ_1-42_ (HFIP-treated, BACHEM) was dissolved in DMSO to obtain a stable stock solution, aliquoted and stored at -20 °C. For the assay, Aβ_1-42_ stock solution was then diluted to a final 50 µM concentration with 10 mM phosphate buffer (pH = 7.4) containing 150 mM NaCl by brief sonication and vortexing. 1 mg of thioflavin T (ThT) was dissolved in methanol to obtain a stock solution which was subsequently diluted in 50 mM glycine-NaOH (pH = 8.6) to 0.4 mM. Assay mixture contains 20 µM ThT. Stock solution of inhibitors were prepared in DMSO.

#### Aβ_1-42_ inhibition assay

Aβ_1-42_ self-aggregation was performed by incubating of 50 µM Aβ_1-42_ solution at room temperature in the assay conditions without any stirring in black, clear bottom 96-well plate (Greiner) by a multi plate reader (Synergy HT, Biotek, Winooski, Vermont, United States) using ThT fluorimetric assay. Final volume of assay mixture was 100 µL. The excitation and emission wavelengths were set at 440/30 and 485/20 nm, respectively. Inhibition experiments were monitored by incubating Aβ_1-42_ at the given conditions in the presence of 50 µM studied samples. As a positive control 50 µM doxycycline was used. Fluorescence data were recorded every 10 min during 72 h incubation time without any stirring. Each inhibitor was assayed in duplicates in at least three independent experiments and the presented values were averaged and are expressed as the mean ± SEM (the standard error of the mean). First the ratio was calculated after subtraction of fluorescence of unbound ThT according to equation 1:

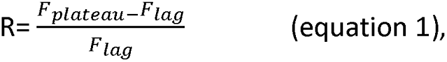

where F_lag_ and F_plateau_ are fluorescence intensities of Aβ_1-42_ in lag and plateau phase of aggregation kinetic at five time points every 30 min, respectively. After that, the percent inhibition of Aβ_1-42_aggregation was calculated as follow (equation 2):

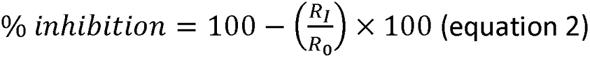

where R_I_ and R_0_ are ratios according to equation 1 with or without studied compounds respectively.

## Acknowledgments

We express our gratitude to Galina Makhaeva team for providing the sample of ipidacrine (amiride). We thank Barbora Hejtmankova for her contribution to the measurement of anti-ChEs properties of final compounds.

## Appendix. Supplementary Information

Supplementary data consists of NMR spectra of the final compounds, as well as LC-MS and LC-UV spectra, the data from measurements of antioxidant properties and inhibition activity toward Aβ aggregation and additional *in silico* experiments.

**Abbreviations**

