## Supplementary Information for "Morphing Cholinesterase Inhibitor Amiridine into Multipotent Drugs for the Treatment of Alzheimer’s Disease"

### 1. Synthesis of intermediates 8, 11 and 12

Derivatives **8**, **11** and **12** were prepared by a slightly modified method from [1]. **General procedure:** Solution of 2-chloroacetyl chloride (4 eq) in chloroform (0.25 M) was added dropwise to cooled (ice bath) solution of amiridine, 1-adamantylamine (**9**) or memantine (**10**) (1 eq) in chloroform (0.43 M). The reaction mixture was stirred at 90 °C overnight. The reaction was monitored by TLC with mobile phases DCM/MeOH/$NH_3$ (9/1/0.1) for **8**, DCM/MeOH/$NH_3$ (50/1/0.1) for **11** and **12** (the spots on TLCs were visualized using a solution of phosphomolybdic acid). After cooling to room temperature, the mixture was washed with 5% (w/v) aqueous solution of $NaHCO_3$ and then with brine. The organic solvent was dried over anhydrous $Na_2SO_4$ and evaporated under reduced pressure with the residual 2-chloroacetyl chloride. The crude product was purified by column chromatography. NMR spectra of **8**, **11** and **12** were in agreement with previously publish data.[1–3]

*2-Chloro-N-{1H,2H,3H,5H,6H,7H,8H-cyclopenta[b]quinolin-9-yl}acetamide* (**8**): Isolated as white solid (1.53 g, 54%). $^{1}$H NMR ($CDCl_3$): $\delta$ 7.92 (s, 1H), 4.16 (s, 2H), 2.93 (t, $J$ = 7.6 Hz, 2H), 2.84 (t, $J$ = 6.3 Hz, 2H), 2.75 (t, $J$ = 7.4 Hz, 2H), 2.53 (t, $J$ = 6.2 Hz, 2H), 2.03 (p, $J$ = 7.6 Hz, 2H), 1.72–1.81 (m, 4H) ppm. $^{13}$C NMR ($CDCl_3$): 164.2, 163.4, 156.5, 137.9, 130.1, 123.7, 42.8, 34.5, 32.7, 29.8, 24.3, 23.0, 22.7, 22.5 ppm. HRMS $[M+H]^+$: 265.11011 (calculated for $[C_{14}H_{18}ClN_2O]^+$: 265.11022).

*N-(adamantan-1-yl)-2-chloroacetamide* (**11**): Isolated as colorless crystals (542 mg, 90%). $^{1}$H NMR ($CDCl_3$): $\delta$ 6.15 (s, 1H), 3.86 (s, 2H), 2.03–2.04 (m, 3H), 1.93–1.96 (m, 6H), 1.60–1.63 (m, 6H) ppm. $^{13}$C NMR ($CDCl_3$): $\delta$ 164.6, 52.4, 42.9, 41.3, 36.2, 29.4 ppm.

*2-Chloro-N-(3,5-dimethyladamantan-1-yl)acetamide* (**12**): Isolated as brown oil (684 mg, 52%). $^{1}$H NMR ($CDCl_3$): $\delta$ 6.17 (s, 1H), 3.85 (s, 2H), 2.10 (p, J = 3.2 Hz, 1H), 1.79–1.80 (m, 2H), 1.62 (ddt, $J$ = 12.1, 2.4, 1.4 Hz, 2H), 1.57 (ddt, $J$ = 11.7, 2.4, 1.2 Hz, 2H), 1.31–1.35 (m, 2H), 1.24 (dtd, $J$ = 12.5, 2.5, 1.4 Hz, 2H), 1.10 (qt, $J$ = 12.4, 2.1 Hz, 2H) ppm. $^{13}$C NMR ($CDCl_3$): $\delta$ 164.7, 54.0, 50.5, 47.2, 42.5, 39.8, 32.4, 30.1, 30.0 ppm.

### 2. Inhibition of cholinesterases

The evaluation of inhibition activities of novel compounds against human recombinant AChE (*h*AChE; E.C. 3.1.1.7) and human serum BChE (*h*BChE, E.C. 3.1.1.8) was determined following the modified spectrophotometric protocol of Ellman *et al.*[4] Final compounds **5**–**7** were primary tested at concentration of 1 µM, then $IC_{50}$ values were assessed for selected derivatives. The percentages of inhibition and $IC_{50}$ values (of **5c-d**, **7c** and **7g**) are summarized in Table S1. Amiridine and THA were used as reference compounds.

*Table S1. Inhibition activities of **5**–**7** and reference compounds (amiridine hydrochloride and THA).*

| Name | Structure | ***h*AChE ± SEM**[a] **(%) c =1 x $10^{-6}$ M ($IC_{50}$)** | ***h*BChE ± SEM**[a] **(%) c =1 x $10^{-6}$ M ($IC_{50}$)** |
|---|---|---|---|
| **5a** | 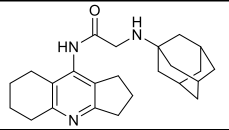 | 2.83 ± 0.49 | 14.01 ± 1.75 |
| **5b** | 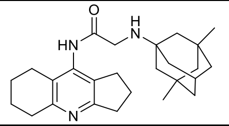 | 9.13 ± 0.48 | 18.61 ± 0.65 |
| **5c** | 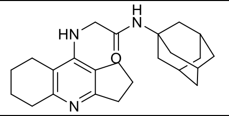 | 1.53 ± 0.38 | 62.66 ± 0.16 ($IC_{50}$ = 0.6 ± 0.02 µM) |
| **5d** | 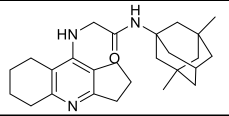 | 5.23 ± 0.25 | 95.16 ± 0.06 ($IC_{50}$ = 0.1 ± 0.004 µM) |
| **6** | 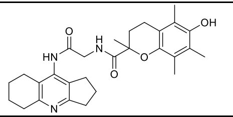 | 3.95 ± 0.25 | 21.52 ± 0.94 |
| **7a** | 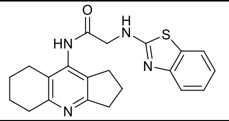 | 1.46 ± 0.17 | 7.19 ± 0.91 |
| **7b** | 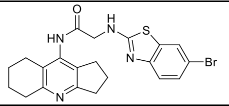 | 18.48 ± 0.37 | 10.83 ± 0.09 |
| **7c** | 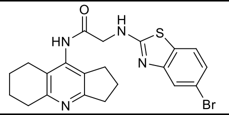 | 15.71 ± 1.17 | 25.00 ± 2.39 ($IC_{50}$ = 4.9 ± 0.35 µM) |
| **7d** | 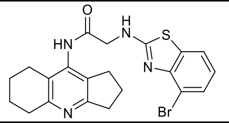 | 1.80 ± 0.33 | 6.73 ± 0.45 |
| **7e** | 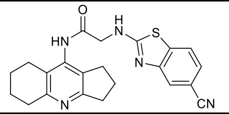 | 13.30 ± 0.57 | 13.92 ± 0.60 |
| **7f** | 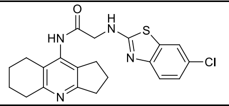 | 12.13 ± 1.21 | 12.96 ± 0.44 |
| **7g** | 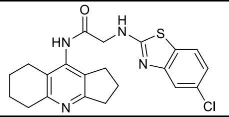 | 11.89 ± 0.74 | 26.46 ± 1.45 ($IC_{50}$ = 11 ± 1.2 µM) |
| **7h** | 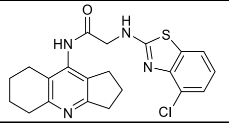 | 4.68 ± 1.48 | 9.26 ± 0.68 |
| **7i** | 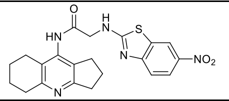 | 7.98 ± 0.99 | 0.93 ± 0.08 |

| | | | |
|---|---|---|---|
| **7j** | 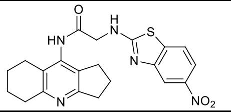 | 8.20 ± 0.41 | 8.03 ± 1.37 |
| **7k** | 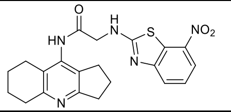 | 9.62 ± 0.41 | 4.47 ± 0.58 |
| **7l** | 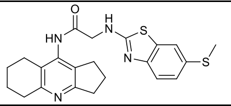 | 7.56 ± 0.38 | 6.05 ± 1.77 |
| **7m** | 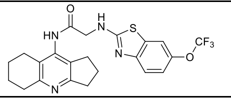 | 13.28 ± 0.71 | 1.50 ± 0.64 |
| **Amiridine.HCl**[b] | 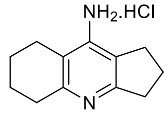 | ($IC_{50}$ = 4.44 ± 0.36 μM) | ($IC_{50}$ = 0.18 ± 0.01 μM) |
| **THA**[c] | 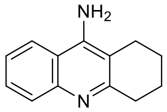 | ($IC_{50}$ = 0.32 ± 0.013 μM) | ($IC_{50}$ = 0.08 ± 0.001 μM) |

[a] Results are expressed as the mean of at least three experiments; [b] taken from reference [1]; [c] taken from reference [5].

#### 3. Cytotoxicity and BBB permeation estimation according to PAMPA

The cytotoxic profile of **5–7** was carried out by MTT (3-(4,5-dimethylthiazol-2-yl)-2,5-diphenyltetrazolium bromide) assay on human neuroblastoma (SH-SY5Y) and hepatocellular carcinoma (HepG2) cell lines as described previously.[6] The cytotoxicity of all the novel compounds was assessed on the SH-SY5Y cells in undifferentiated and in differentiated form. The data obtained from the measurements on the undifferentiated cells are summarized in Table S2, while the measurements on differentiated cells are shown on Figure S1.

To investigate the possibility of **5–7** to cross the BBB and thus potentially be CNS active drugs, we used two tools, a parallel artificial membrane permeation assay (PAMPA) and an algorithm called "BBB score".[7–9] The results of final derivatives and reference compounds (amiridine.HCl and THA) for PAMPA are shown in Table S2.

*Table S2. Cytotoxicity profiles of **5–7** on SH-SY5Y cell line and predictions to BBB penetration by PAMPA results.*

| **Name** | **Structure** | **Cytotoxicity $IC_{50}$ ± SEM (μM)** | | **BBB permeation estimation** | |
|---|---|---|---|---|---|
| | | **SH-SY5Y** | **HepG2** | **$P_e$ ± SEM ($10^{-6}$ cm.s$^{-1}$)**[a] | **CNS (+/-)**[b] |
| **5a** | 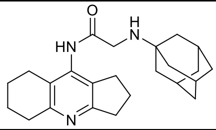 | 74.02 ± 2.80 | 67.89 ± 1.12 | 23.35 ± 1.86 | CNS + |

| | | | | | |
|---|---|---|---|---|---|
| **5b** | 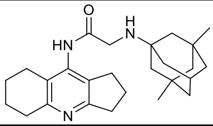 | 31.02 ± 2.76 | 19.54 ± 1.72 | 7.63 ± 0.09 | CNS + |
| **5c** | 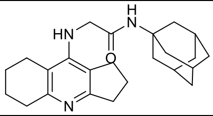 | ≥ 200 | 186.69 ± 11.66 | 1.63 ± 0.19 | CNS - |
| **5d** | 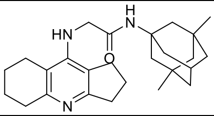 | 112.45 ± 1.45 | 41.57 ± 3.51 | 2.23 ± 0.05 | CNS +/- |
| **6** | 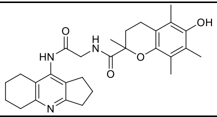 | ≥ 200 | n.d.[c] | 15.93 ± 0.38 | CNS + |
| **7a** | 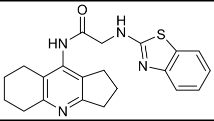 | 72.31 ± 6.89 | ≥ 200 | 20.64 ± 0.44 | CNS + |
| **7b** | 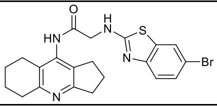 | ≥ 200 | ≥ 200 | 3.74 ± 0.36 | CNS +/- |
| **7c** | 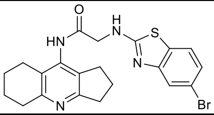 | 17.68 ± 2.34 | 21.61 ± 0.21 | 2.13 ± 1.04 | CNS +/- |
| **7d** | 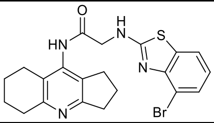 | 111.40 ± 5.60 | 14.47 ± 1.84 | 2.62 ± 0.00 | CNS +/- |
| **7e** | 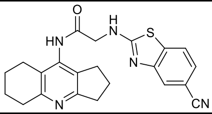 | 77.35 ± 4.54 | 26.41 ± 1.82 | 13.90 ± 1.26 | CNS + |
| **7f** | 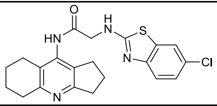 | 145.57 ± 14.48 | ≥ 200 | 2.17 ± 0.04 | CNS +/- |
| **7g** | 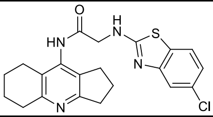 | 98.24 ± 5.74 | ≥ 200 | 2.80 ± 0.79 | CNS +/- |
| **7h** | 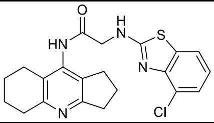 | 9.52 ± 1.85 | 31.27 ± 3.48 | 1.94 ± 0.37 | CNS - |
| **7i** | 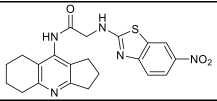 | 91.27 ± 11.02 | ≥ 200 | 4.89 ± 1.50 | CNS + |
| **7j** | 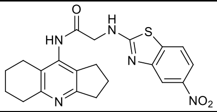 | 93.73 ± 5.38 | 25.83 ± 0.25 | 1.83 ± 0.54 | CNS - |
| **7k** | 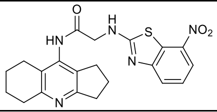 | 51.04 ± 8.56 | ≥ 200 | 1.12 ± 0.62 | CNS - |
| **7l** | 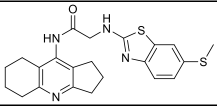 | 172.55 ± 17.55 | ≥ 200 | 2.65 ± 0.01 | CNS +/- |

| | | | | | |
|---|---|---|---|---|---|
| **7m** | 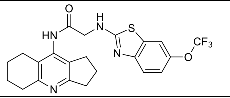 | 29.51 ± 2.30 | ≥ 200 | 2.54 ± 0.42 | CNS +/- |
| **Amiridine.HCl** | 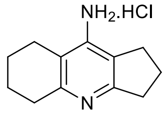 | 597.47 ± 25.49 | n.d.[c] | 5.8 ± 0.5 | CNS + |
| **THA** | 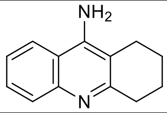 | 122.25 ± 1.59 | 168.47 ± 3.63[d] | 6.0 ± 0.6 | CNS + |

[a] Results are expressed as the mean of at least two experiments; [b] classification of the prediction to cross the BBB, CNS +: high BBB permeation predicted with $P_e$ ($10^{-6}$ cm.s$^{-1}$) > 4.0, CNS -: low BBB permeation predicted with $P_e$ ($10^{-6}$ cm.s$^{-1}$) < 2.0, CNS +/-: BBB permeation uncertain with $P_e$ ($10^{-6}$ cm.s$^{-1}$) from 4.0 to 2.0; [c] n.d. = not determined; [d] taken from reference .

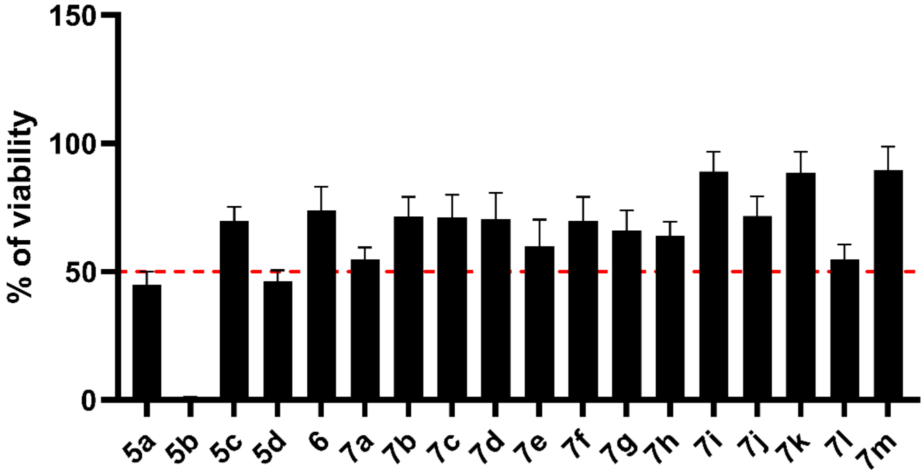

*Figure S1. The viability of differentiated SH-SY5Y cells treated with tested compounds in the concentration corresponding to the $IC_{50}$ values obtained on normal (undifferentiated) SH-SY5Y cells.*

#### 4. BBB Score and Drug-Likeness

To explore the drug-likeness of our compounds, we compute BBB score according to Gupta *et al.*, and used SwissADME website tool (http://www.swissadme.ch/index.php) to add other in silico pharmacokinetic and drug-like properties. Our focus was on gastrointestinal absorption (GIA), Lipinski's rule of five, bioavailability, and PAINS violations (Pan Assay Interference Structures). The results are shown in Table S3. The bioavailability radar charts of all final derivatives are displayed on Figure S2.

*Table S3. Summary of selected in silico pharmacokinetic properties and drug-likeness of* ***K2508–K2525*** *using various rules and prediction models.*

| Compound | GIA[a] | BBB score[b] | Lipinski[c] | Bio. Score[d] | PAINS[e] | Bioavailability Violations |
|---|---|---|---|---|---|---|
| **5a** | High | 4.8 | Yes | 0.55 | 0 alert | Solubility<br>Lipophilicity |

| | | | | | | |
|---|---|---|---|---|---|---|
| **5b** | High | 4.8 | Yes | 0.55 | 0 alert | Solubility Lipophilicity |
| **5c** | High | 4.9 | Yes | 0.55 | 0 alert | Solubility Lipophilicity |
| **5d** | High | 4.8 | Yes | 0.55 | 0 alert | Solubility Lipophilicity |
| **6** | High | 3.5 | Yes | 0.55 | 0 alert | Solubility Lipophilicity |
| **7a** | High | 4.1 | Yes | 0.55 | 0 alert | Solubility Lipophilicity |
| **7b** | High | 4.1 | Yes | 0.55 | 0 alert | Solubility Lipophilicity |
| **7c** | High | 4.1 | Yes | 0.55 | 0 alert | Solubility Lipophilicity |
| **7d** | High | 4.1 | Yes | 0.55 | 0 alert | Solubility Lipophilicity |
| **7e** | High | 3.5 | Yes | 0.55 | 0 alert | Solubility Lipophilicity |
| **7f** | High | 4.1 | Yes | 0.55 | 0 alert | Solubility Lipophilicity |
| **7g** | High | 4.1 | Yes | 0.55 | 0 alert | Solubility Lipophilicity |
| **7h** | High | 4.1 | Yes | 0.55 | 0 alert | Solubility Lipophilicity |
| **7i** | Low | 2.9 | Yes | 0.55 | 0 alert | Solubility Lipophilicity Nitro group[f] |
| **7j** | Low | 2.9 | Yes | 0.55 | 0 alert | Solubility Lipophilicity Nitro group[f] |
| **7k** | Low | 2.9 | Yes | 0.55 | 0 alert | Solubility Lipophilicity Nitro group[f] |
| **7l** | High | 3.7 | Yes | 0.55 | 0 alert | Solubility Lipophilicity |
| **7m** | Low | 2.9 | Yes | 0.55 | 0 alert | Solubility Lipophilicity |

[a] GIA = gastrointestinal absorption [10,11]; [b] BBB score = blood-brain barrier permeation prediction [9]; [c] All compounds showed 0 violation against Lipinski's rule of five [12]; [d] Bio. Score = Abbott bioavailability score [13]; [e] PAINS = Pan Assay Interference Structures [14]; [f] Structural alert according to [15].

5a

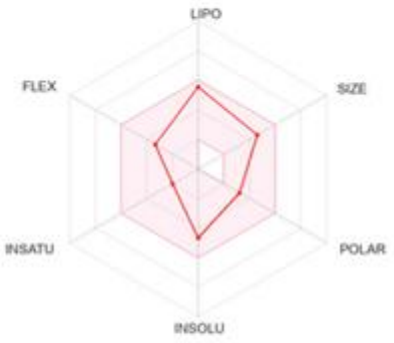

5b

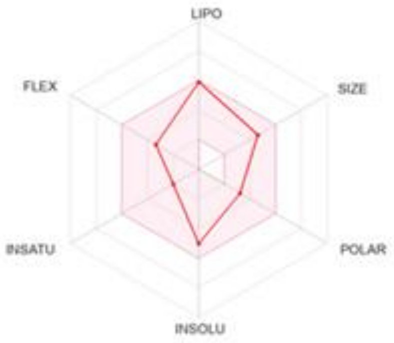

5c

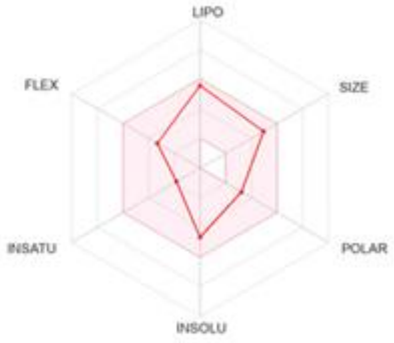

5d

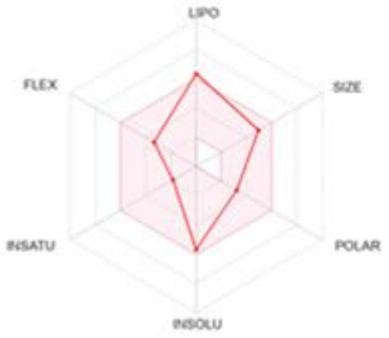

6

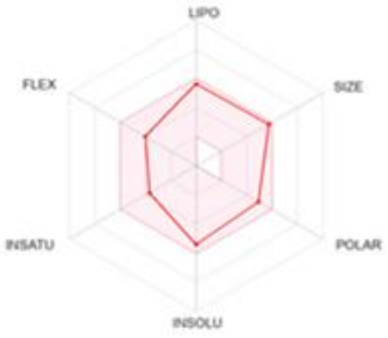

7a

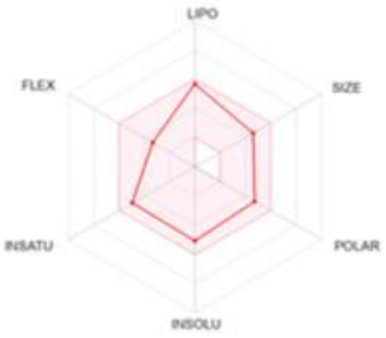

7b

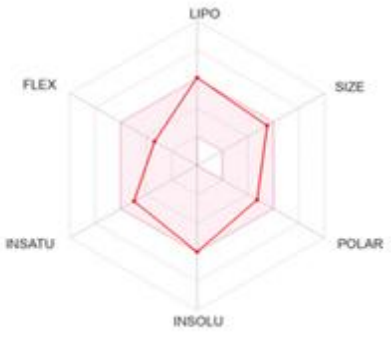

7c

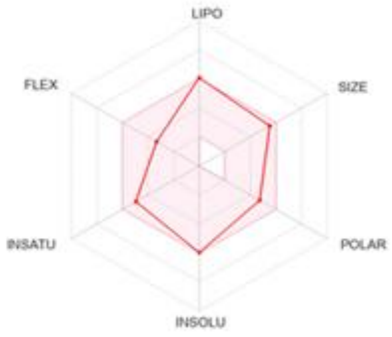

7d

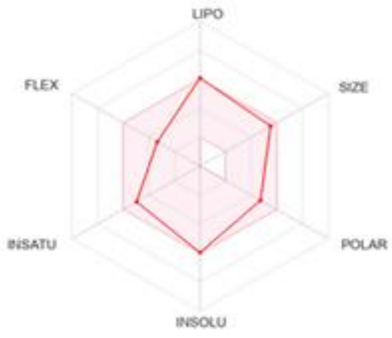

7e

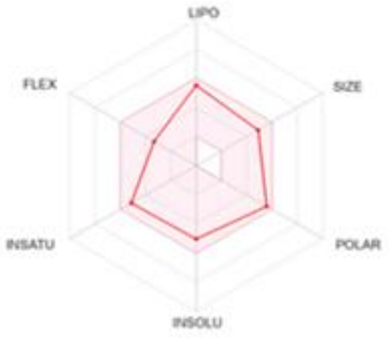

7f

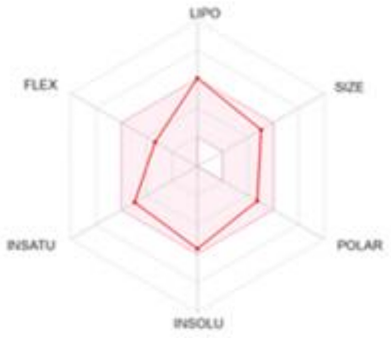

7g

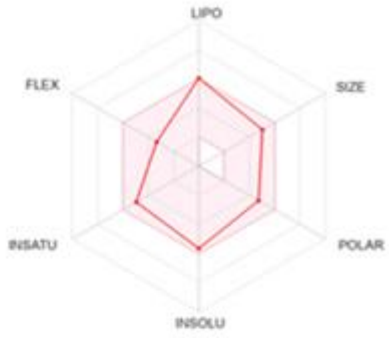

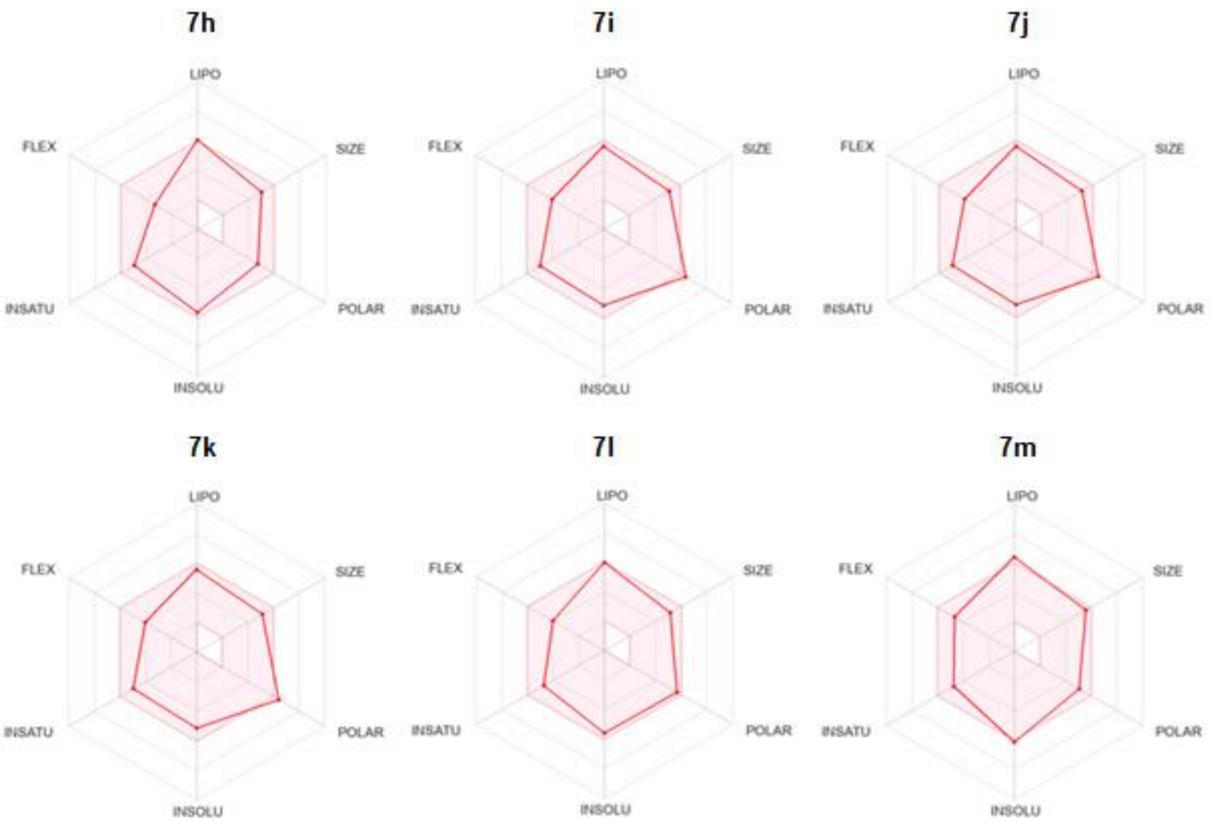

*Figure S2. The bioavailability radar charts of final derivatives* ***5**–**7**. The pink middle area represents the optimum range for bioavailability. It sums the lipophilicity (LIPO), size, polarity (POLAR), solubility (INSOLU), saturation, (INSATU), and flexibility (FLEX).*

### 5. Antioxidant properties

The compound **6** was assessed for its antioxidant properties *via* the DPPH (diphenyl-1-picrylhydrazyl) free radical assay. Antioxidant activity was quantified as $EC_{50}$, representing the concentration of the compound necessary to induce a 50% reduction in DPPH activity, with trolox serving as the standard. The data are detailed in Table S4.

*Table S4. Antioxidant efficiency of 6 and control compound trolox.*

| Name | Structure | Antioxidant efficiency (activity) $EC_{50}$ ± SEM (µM) |
|---|---|---|
| **6** | 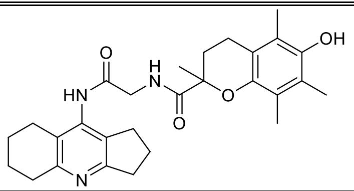 | 113.60 ± 3.40 |
| **Trolox** | 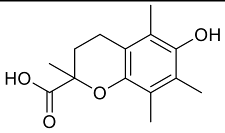 | 16.20 ± 0.42 |

#### 6. Inhibition of amyloid beta aggregation

**7c** and **7m** were tested to determine their inhibitory activities against $A\beta_{1\text{-}42}$ self-induced aggregation. The results for these selected compounds, alongside doxycycline as a positive control, are presented in Table S5 and Figure S3.

*Table S5.Inhibition activities of **7c**, **7m**, and a positive control doxycycline against self-induced $A\beta_{1\text{-}42}$ aggregation.*

| Name | Structure | **Inhibition of $A\beta_{1\text{-}42}$ aggregates ± SEM (%)*** |
|---|---|---|
| **7c** | 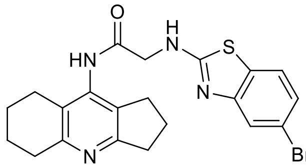 | ≈ 0 |
| **7m** | 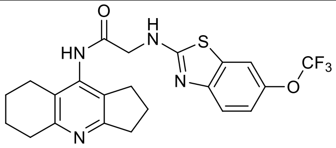 | ≈ 0 |
| **Doxycycline** | 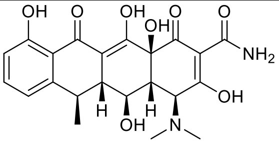 | ≈ 100 |

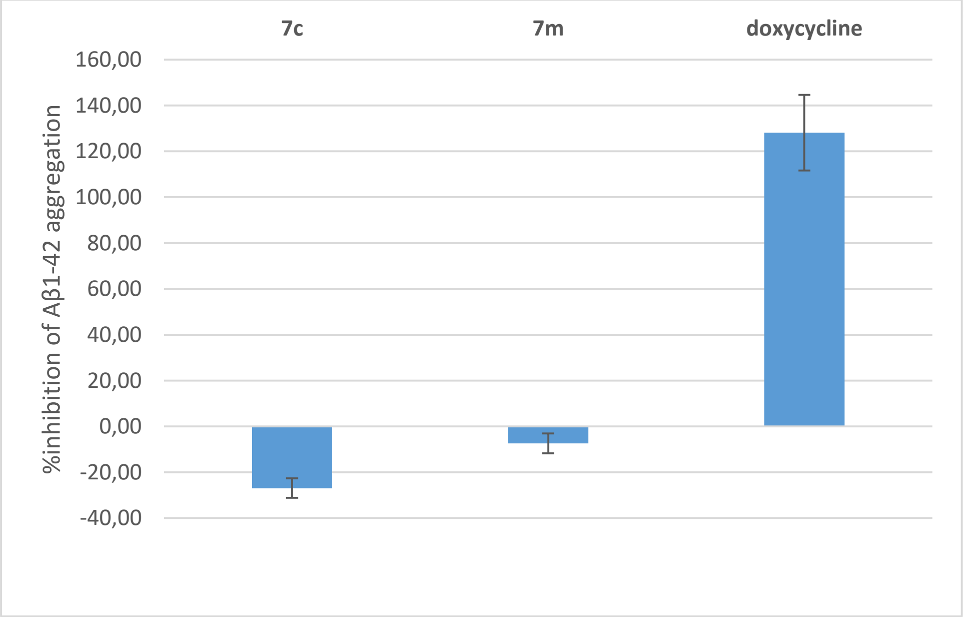

*Figure S3. Inhibition of $A\beta_{1\text{-}42}$ self-aggregation by **7c**, **7m**, and a positive control doxycycline.*

### 7. NMR spectra of prepared derivatives

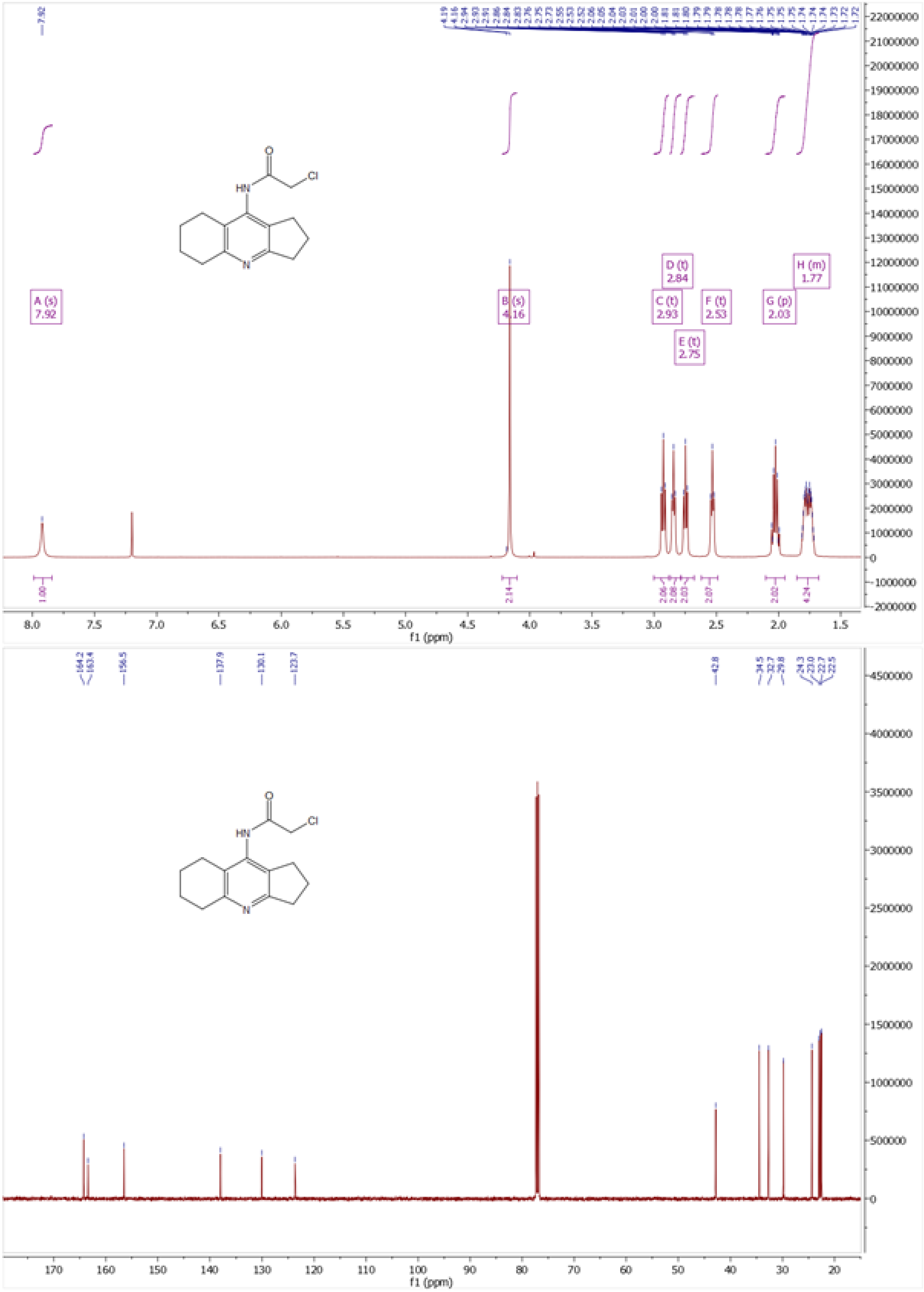

*Figure S4. $^{1}H$ and $^{13}C$ NMR spectra of compound* ***8***

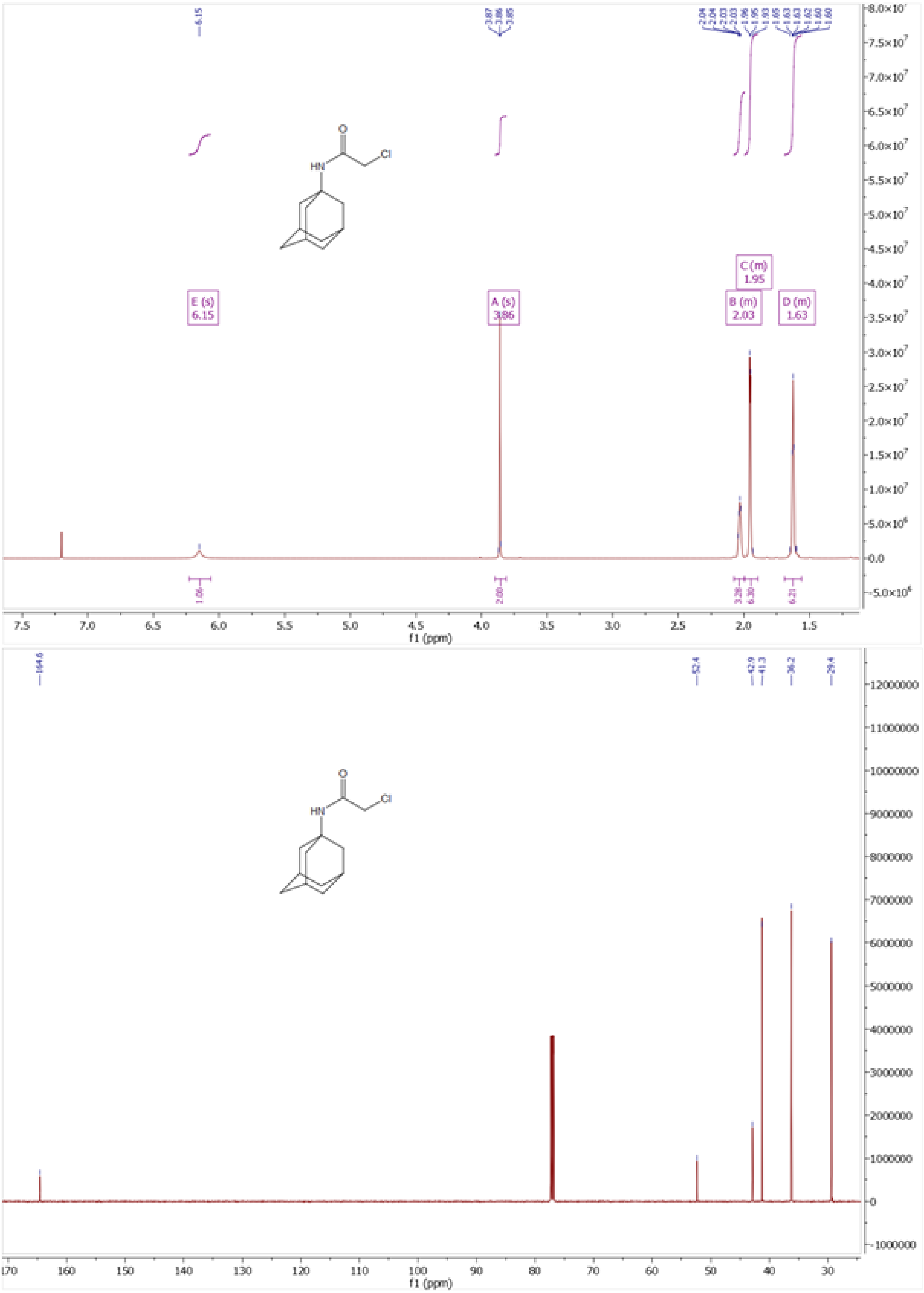

*Figure S5. $^{1}H$ and $^{13}C$ NMR of compound* ***11***

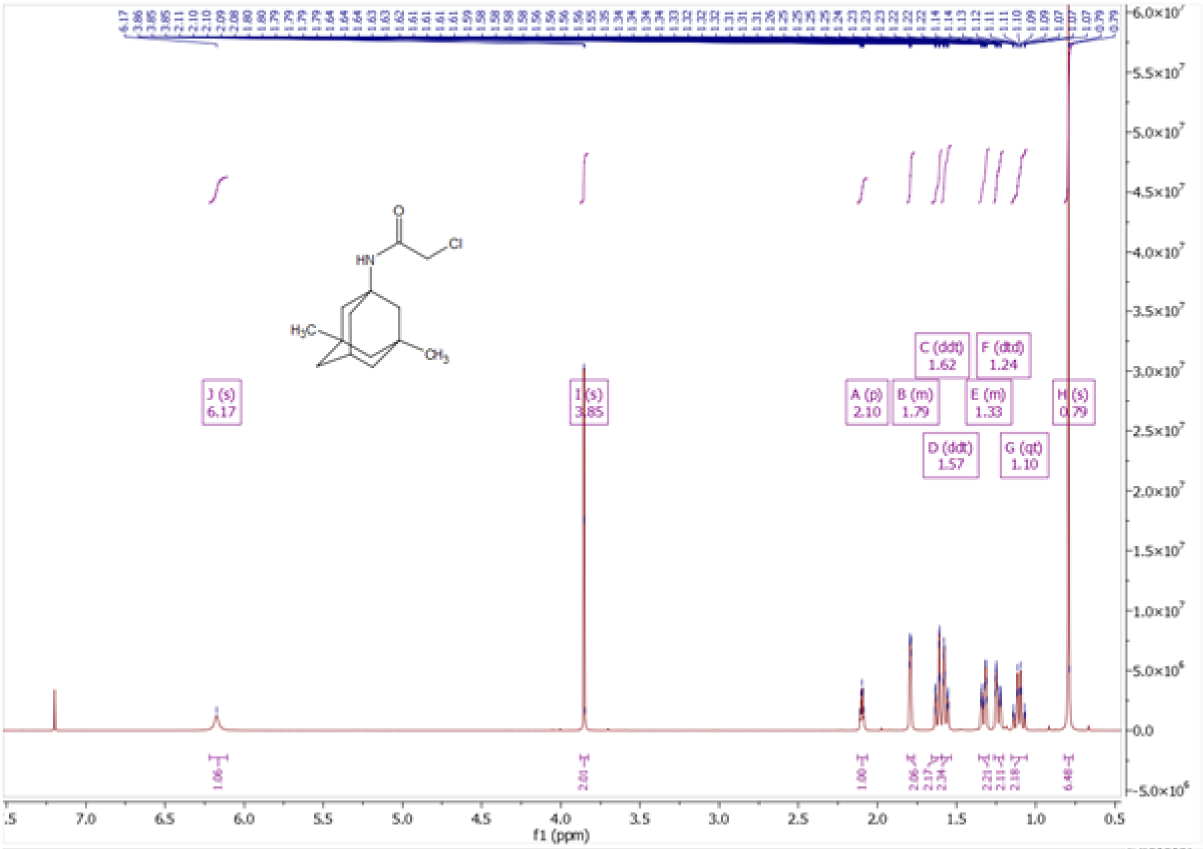

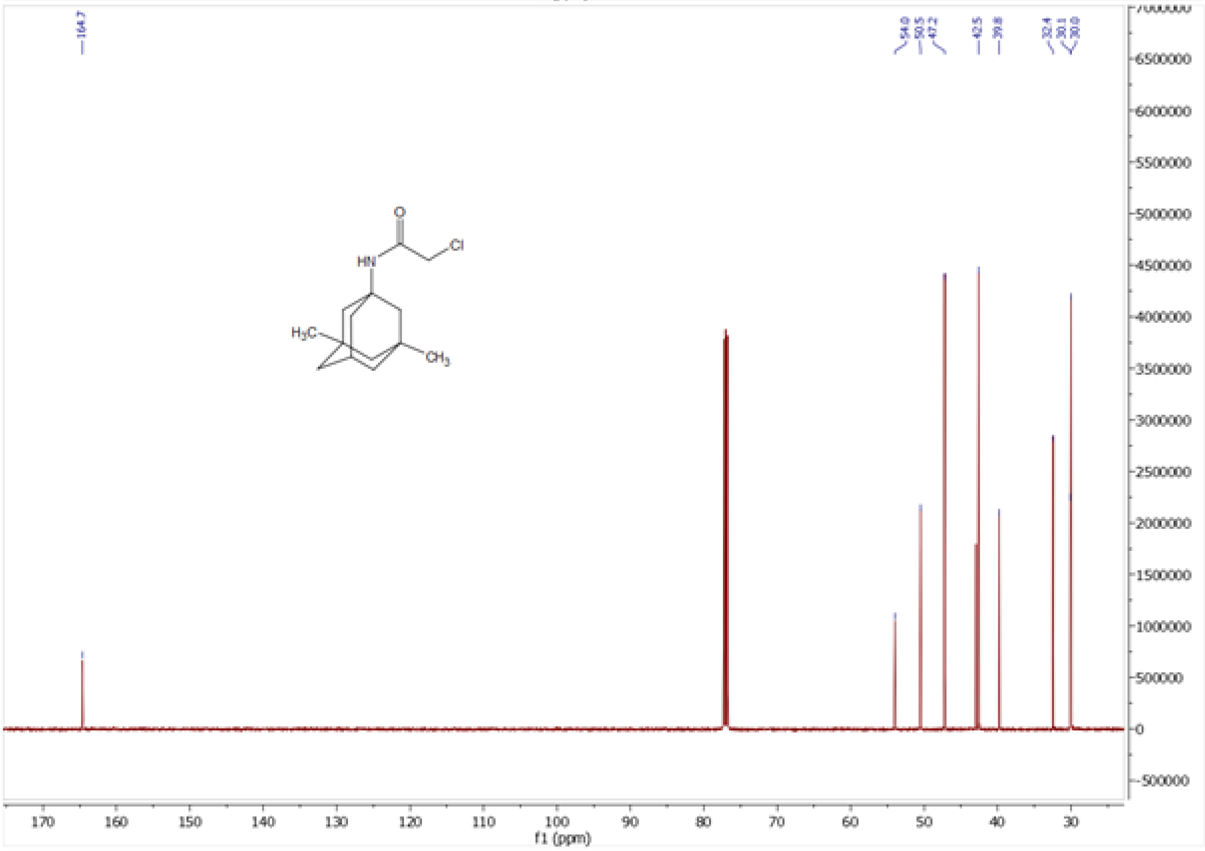

*Figure S6. [1]H and [13]C NMR spectra of compound **12***

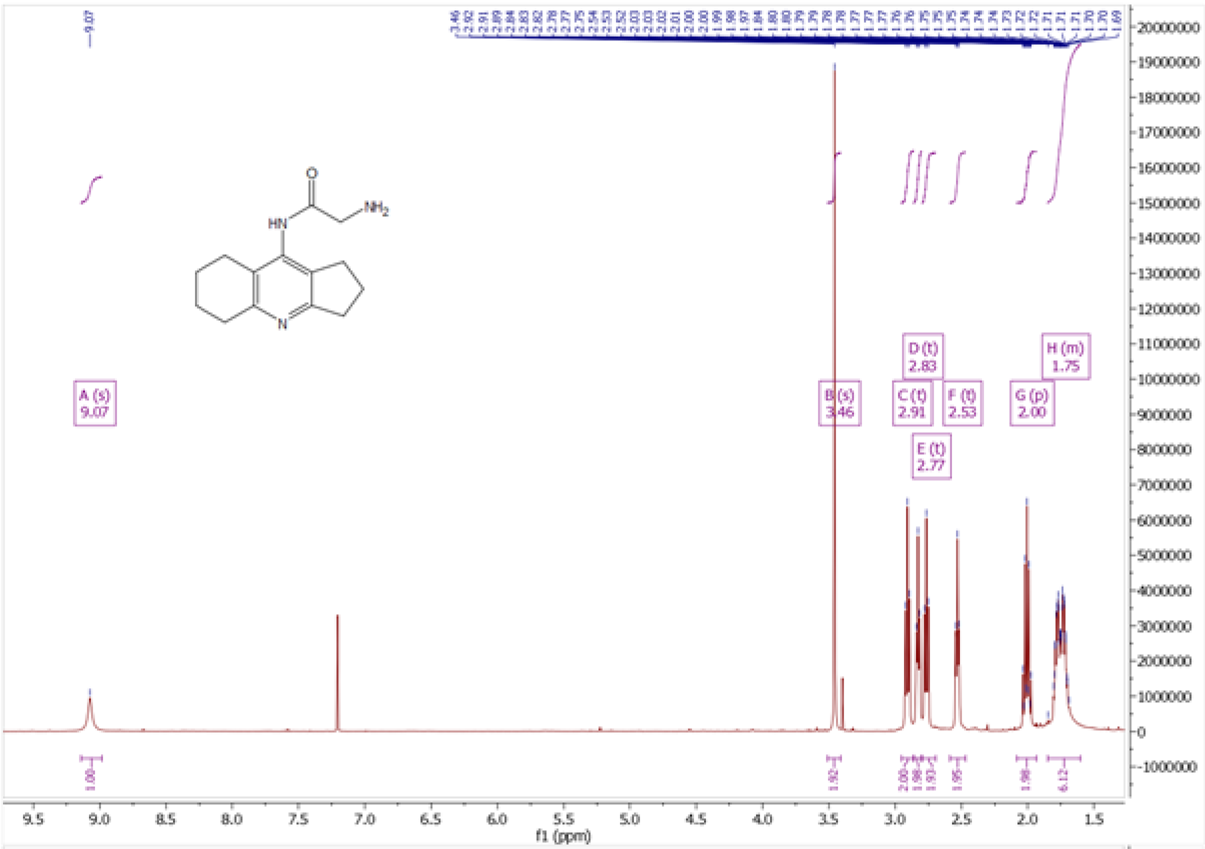

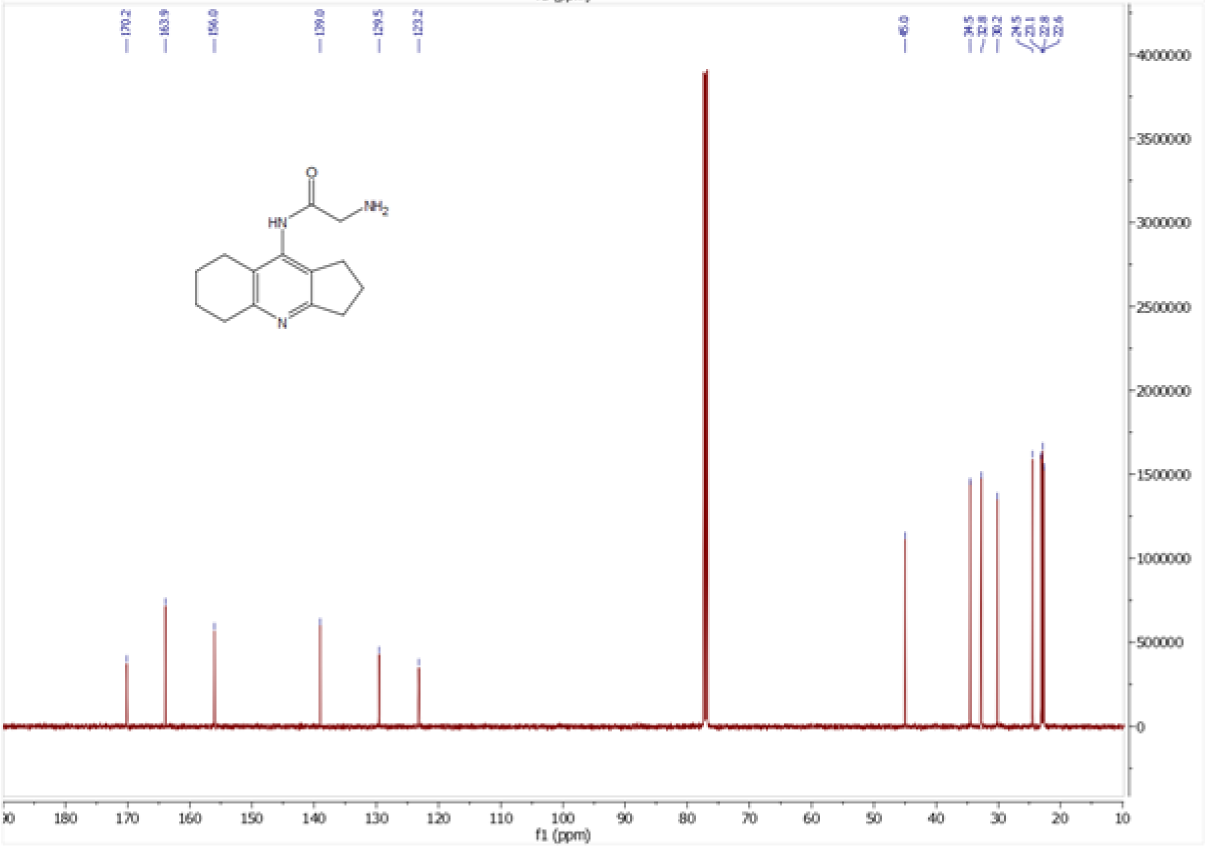

*Figure S7. ¹H and ¹³C NMR spectra of compound* ***13****.*

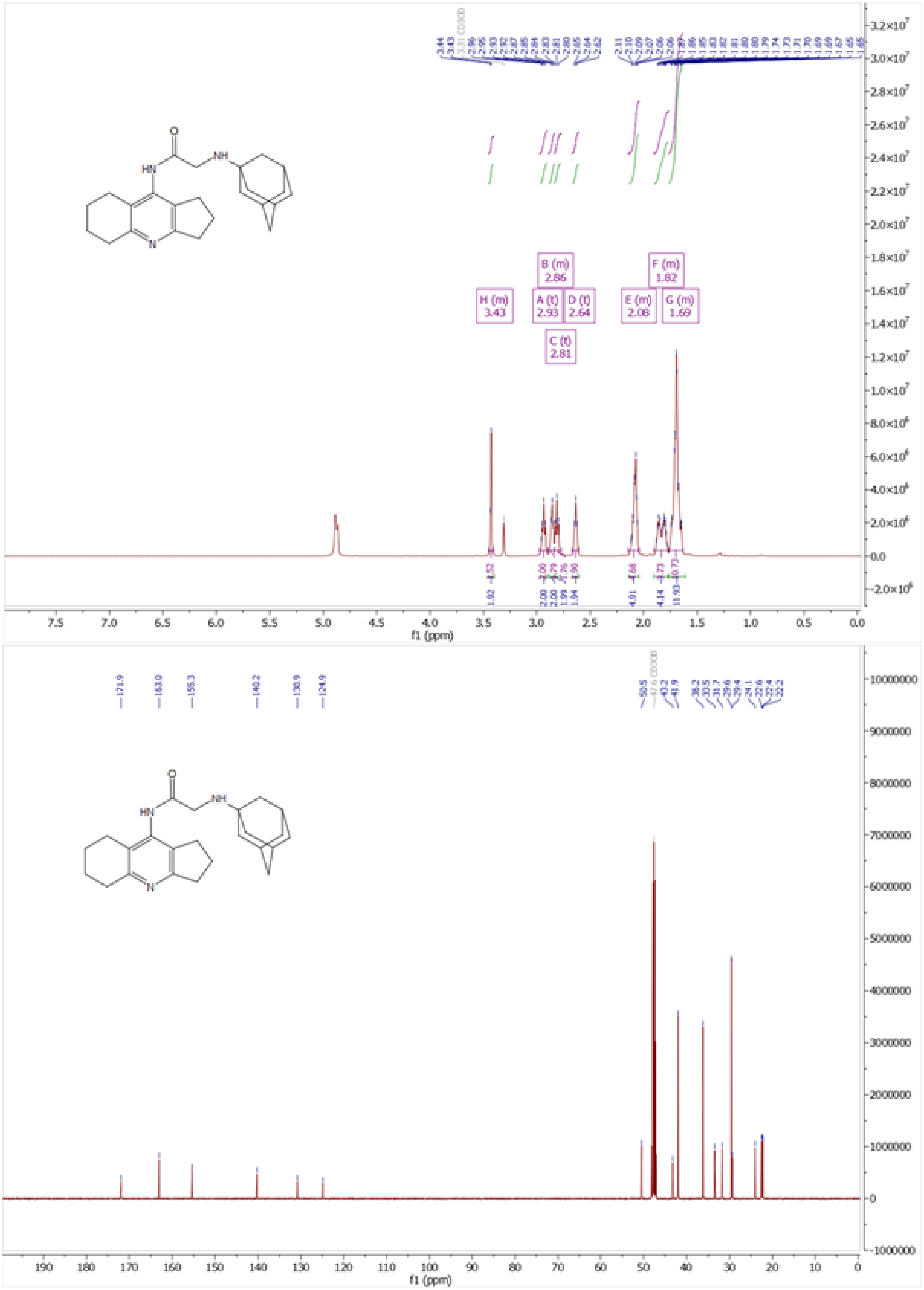

*Figure S8.* $^{1}H$ *and* $^{13}C$ *NMR spectra of* ***5a***

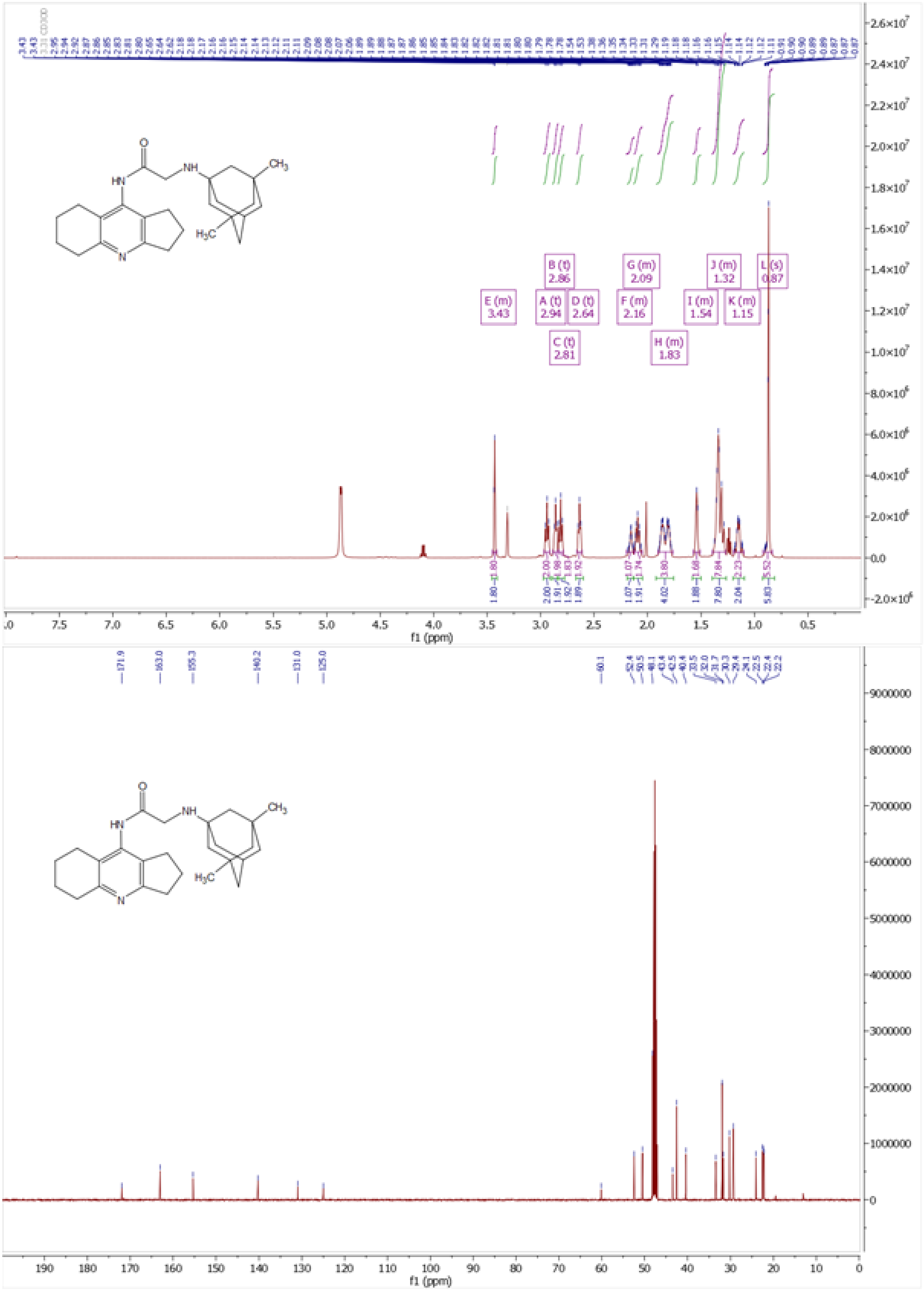

*Figure S9. [1]H and [13]C NMR spectra of **5b***

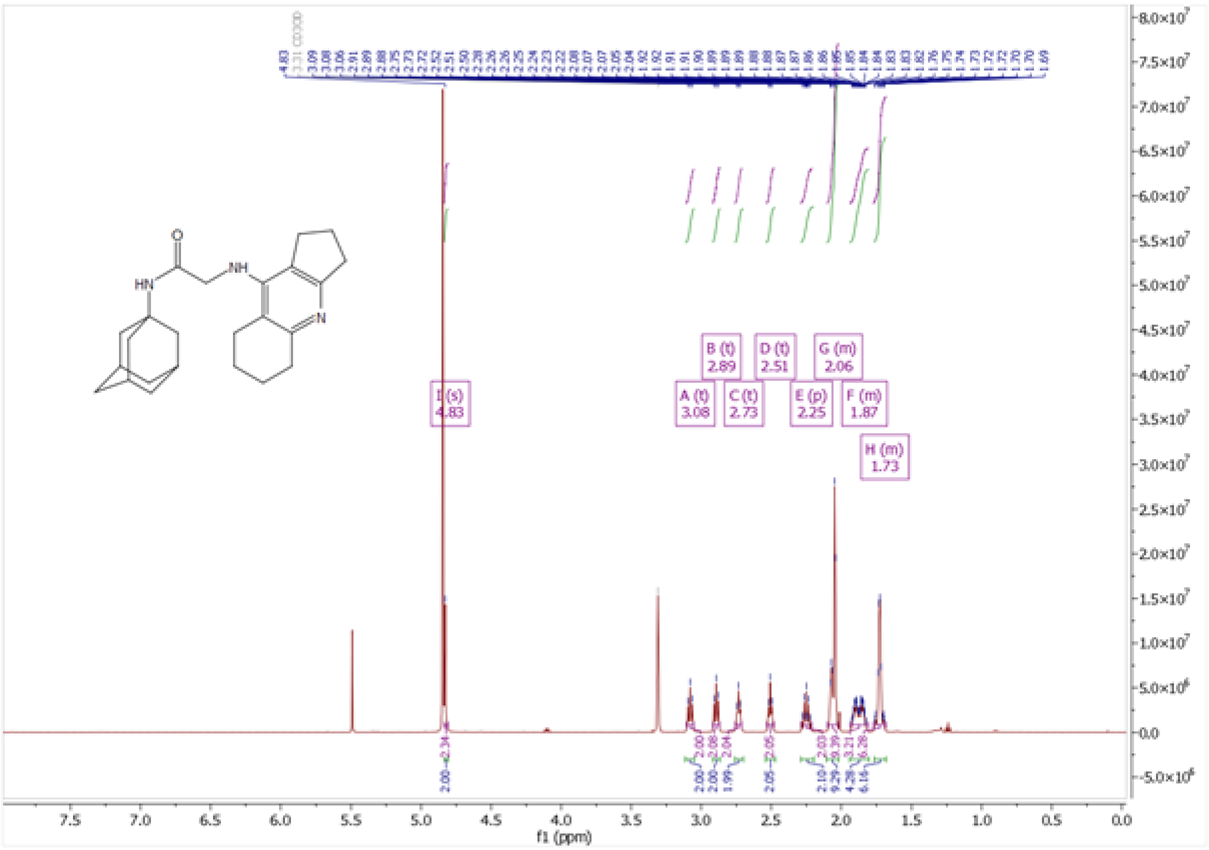

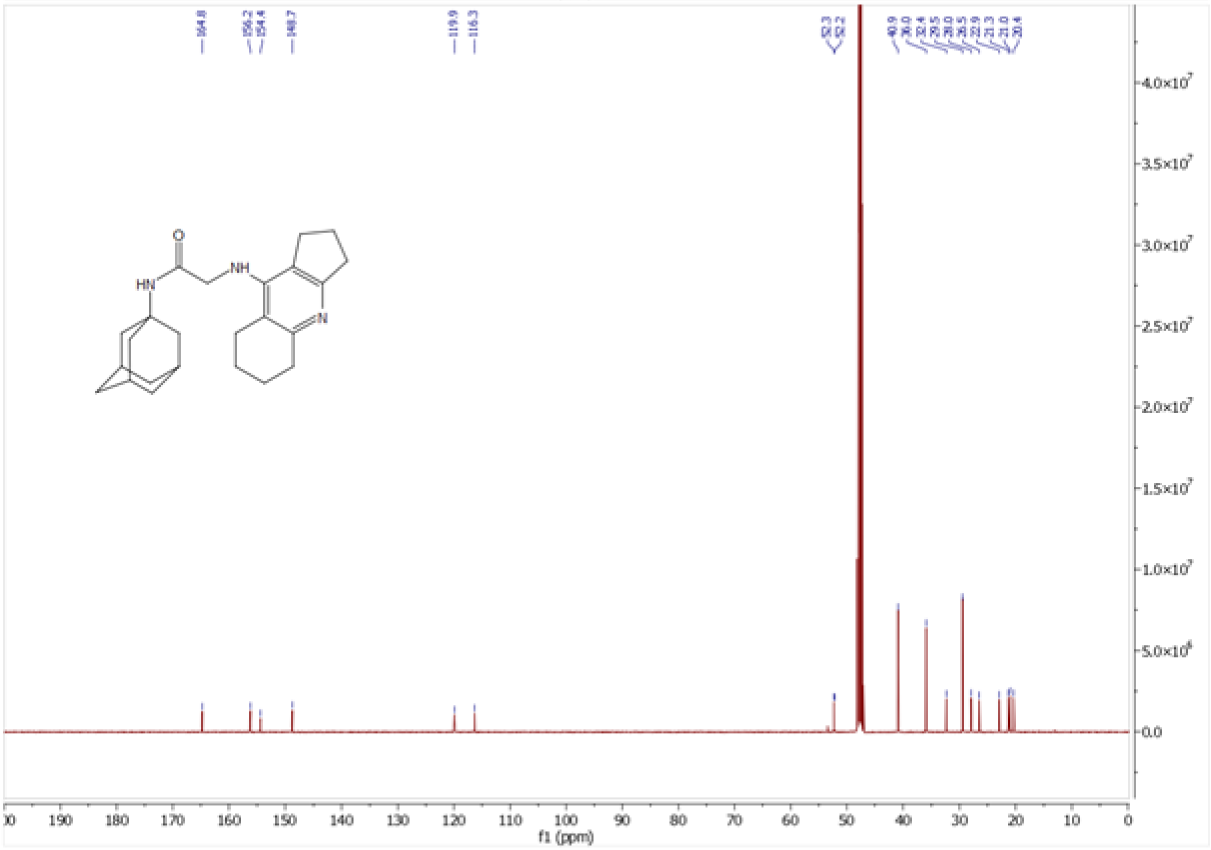

*Figure S10. [1]H and [13]C NMR spectra of **5c***

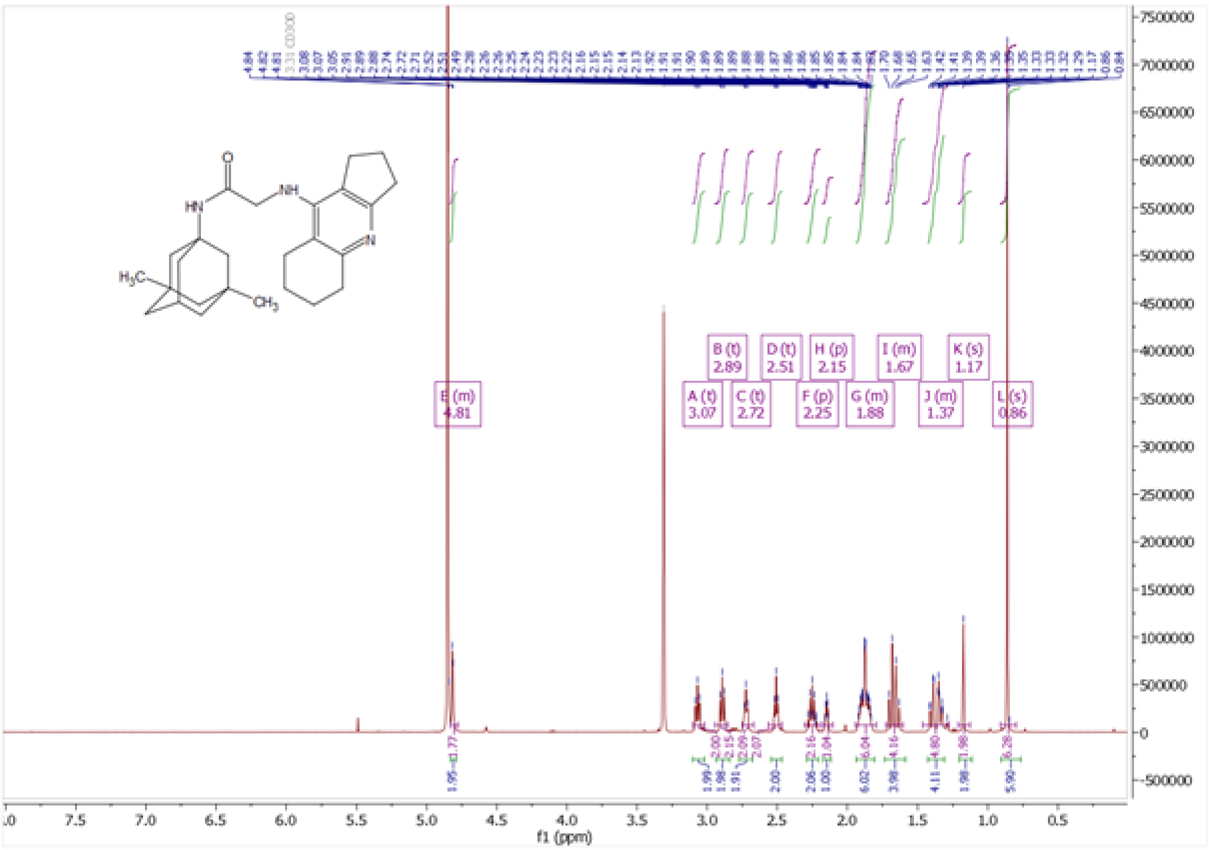

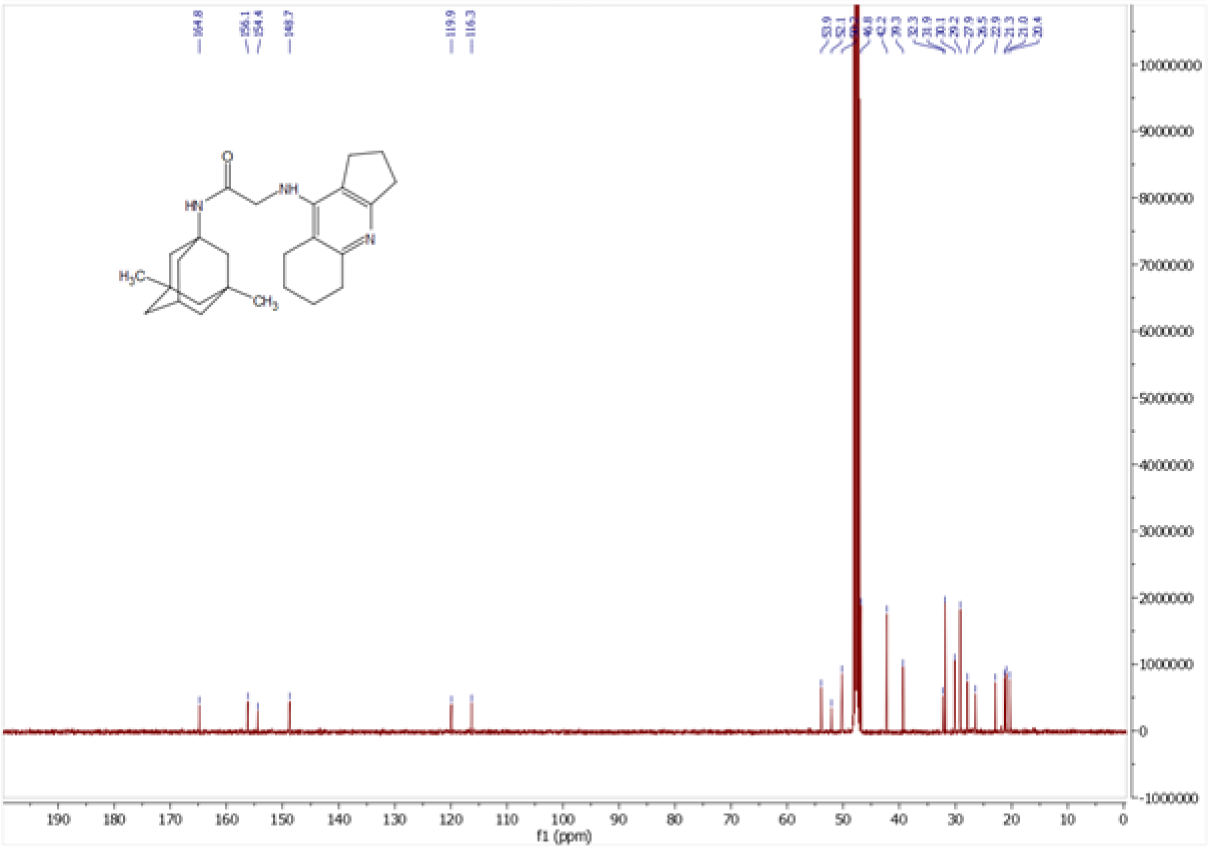

*Figure S11. [1]H and [13]C NMR spectra of* ***5d***

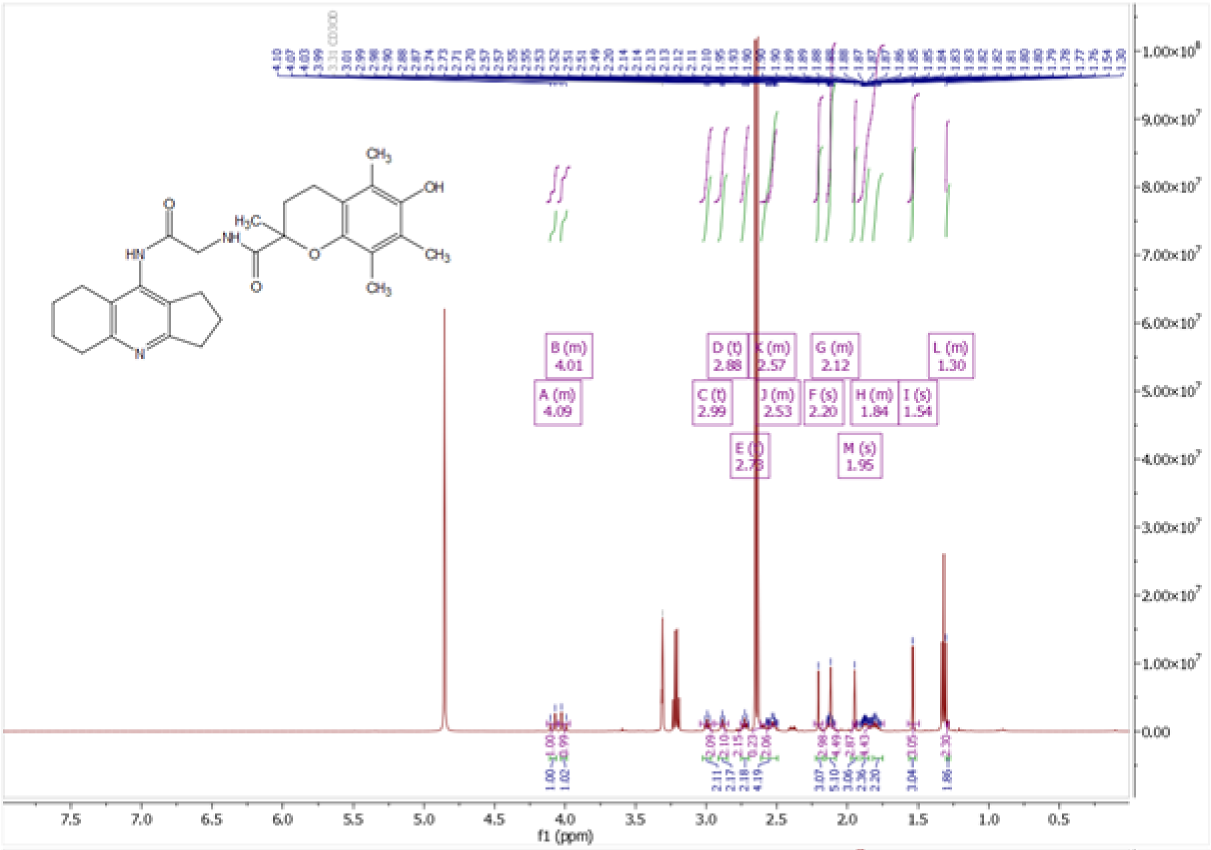

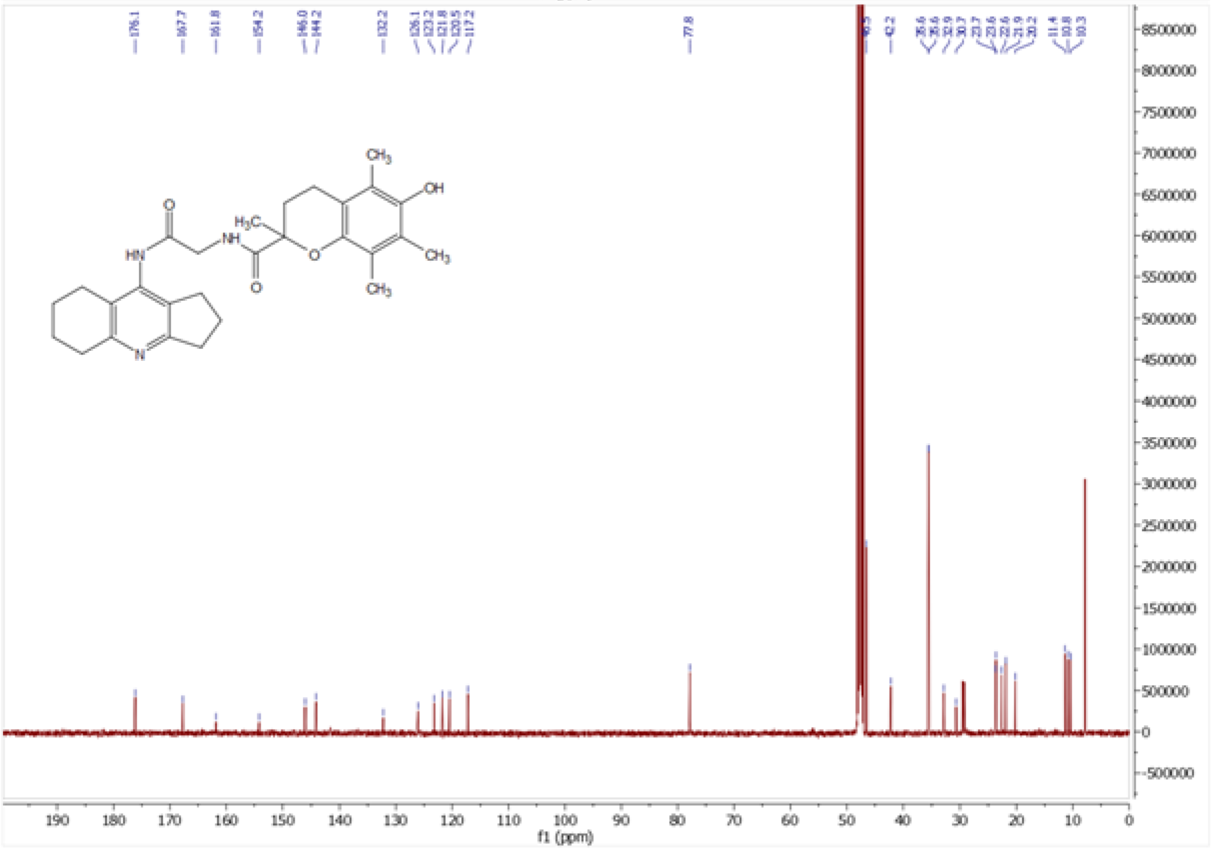

*Figure S12.* *$^{1}$H and $^{13}$C NMR spectra of* ***6***

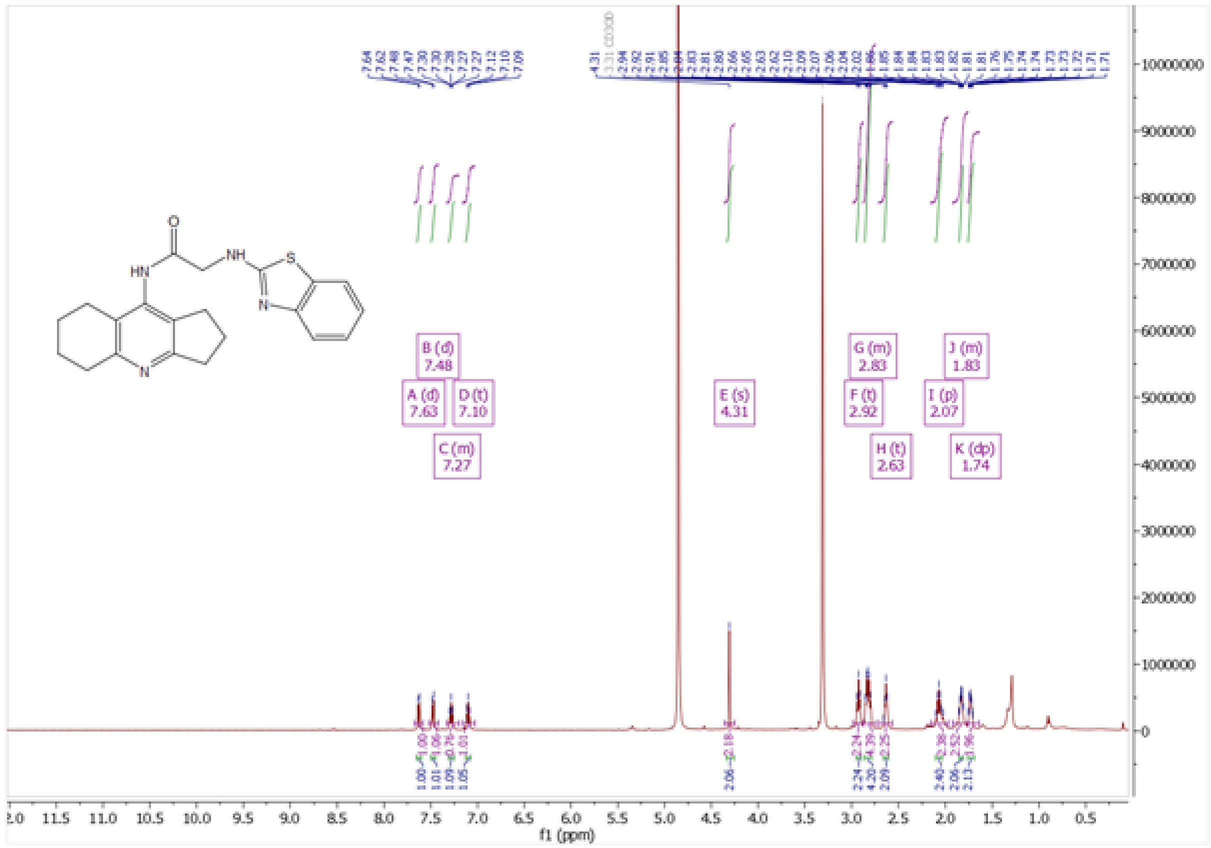

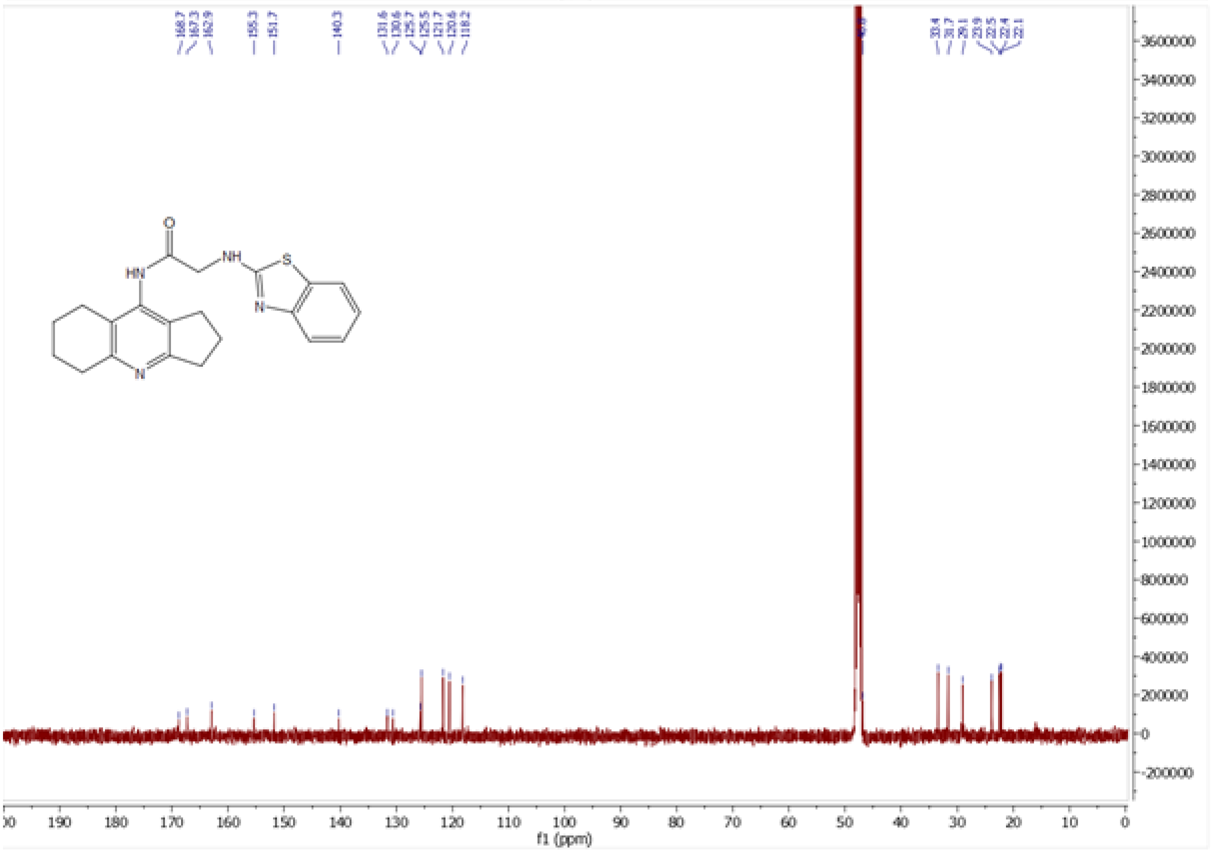

*Figure S13. ¹H and ¹³C NMR spectra of **7a***

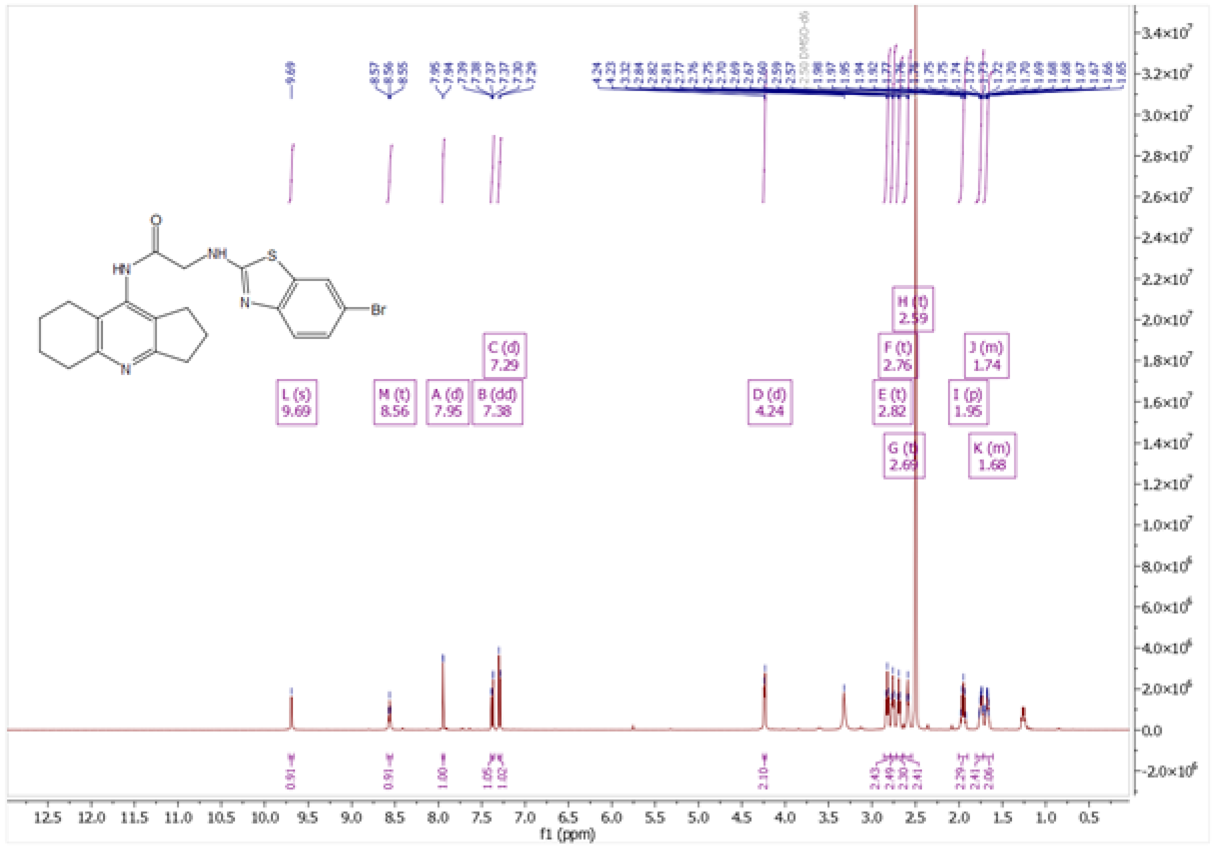

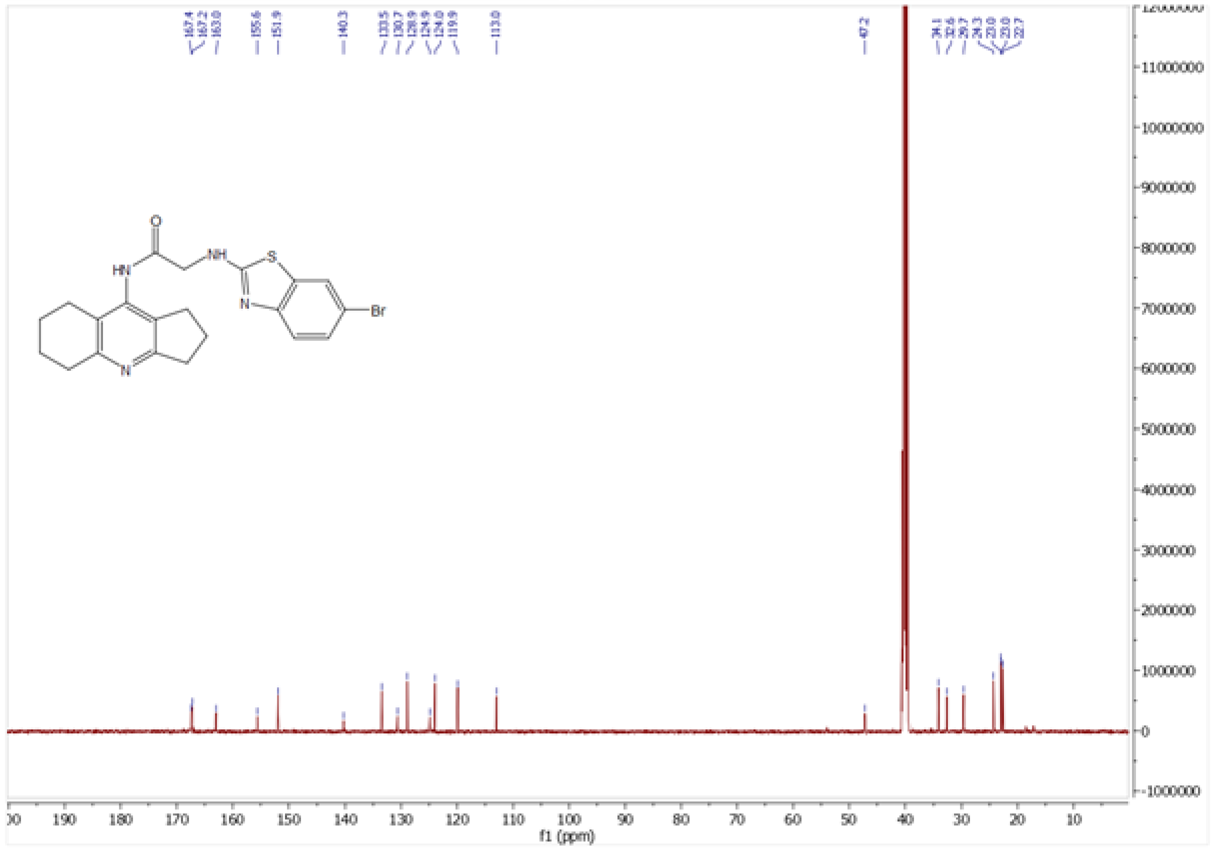

Figure S14. [1]H and [13]C NMR spectra of **7b**

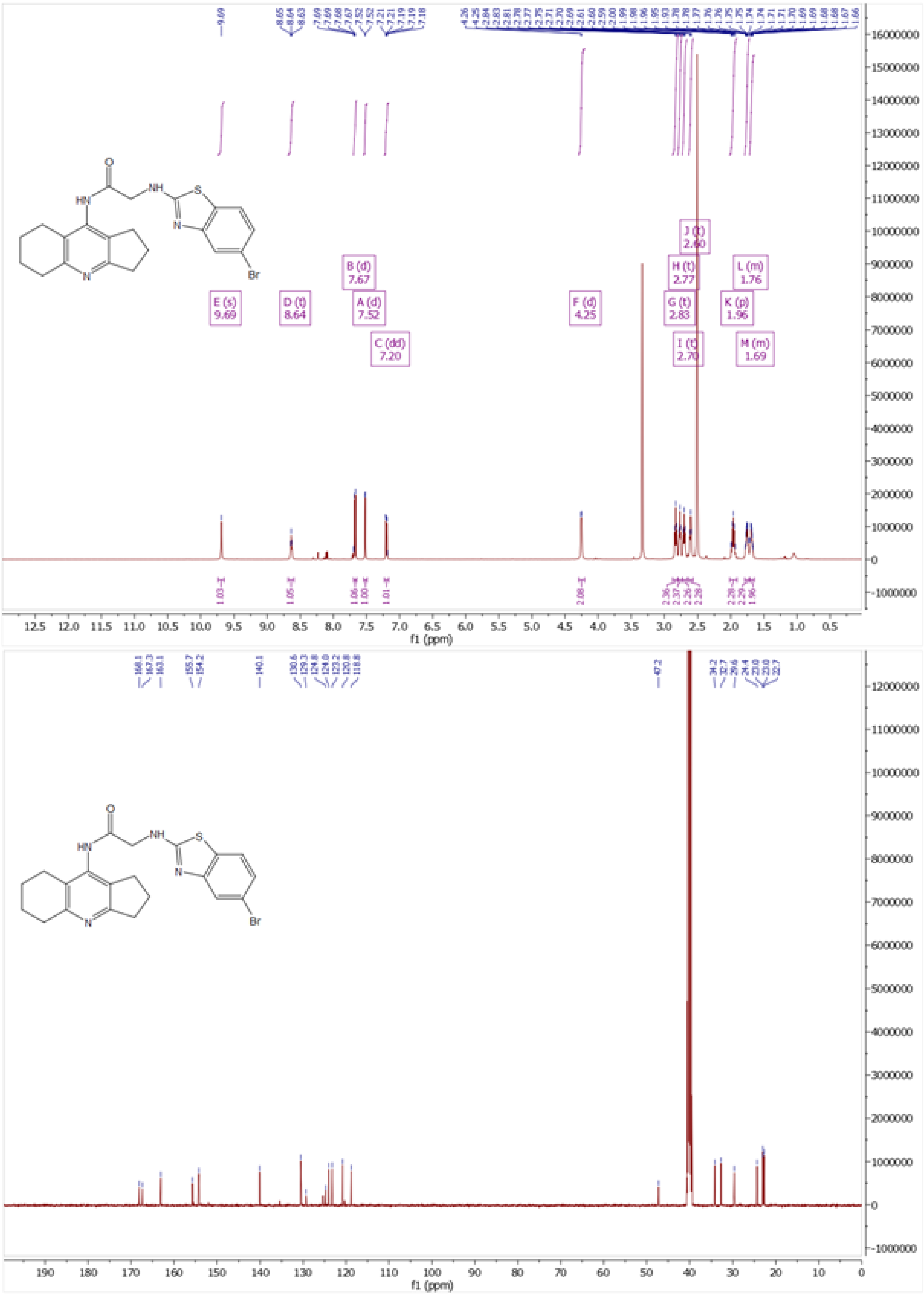

*Figure S15. $^1$H and $^{13}$C NMR spectra of **7c***

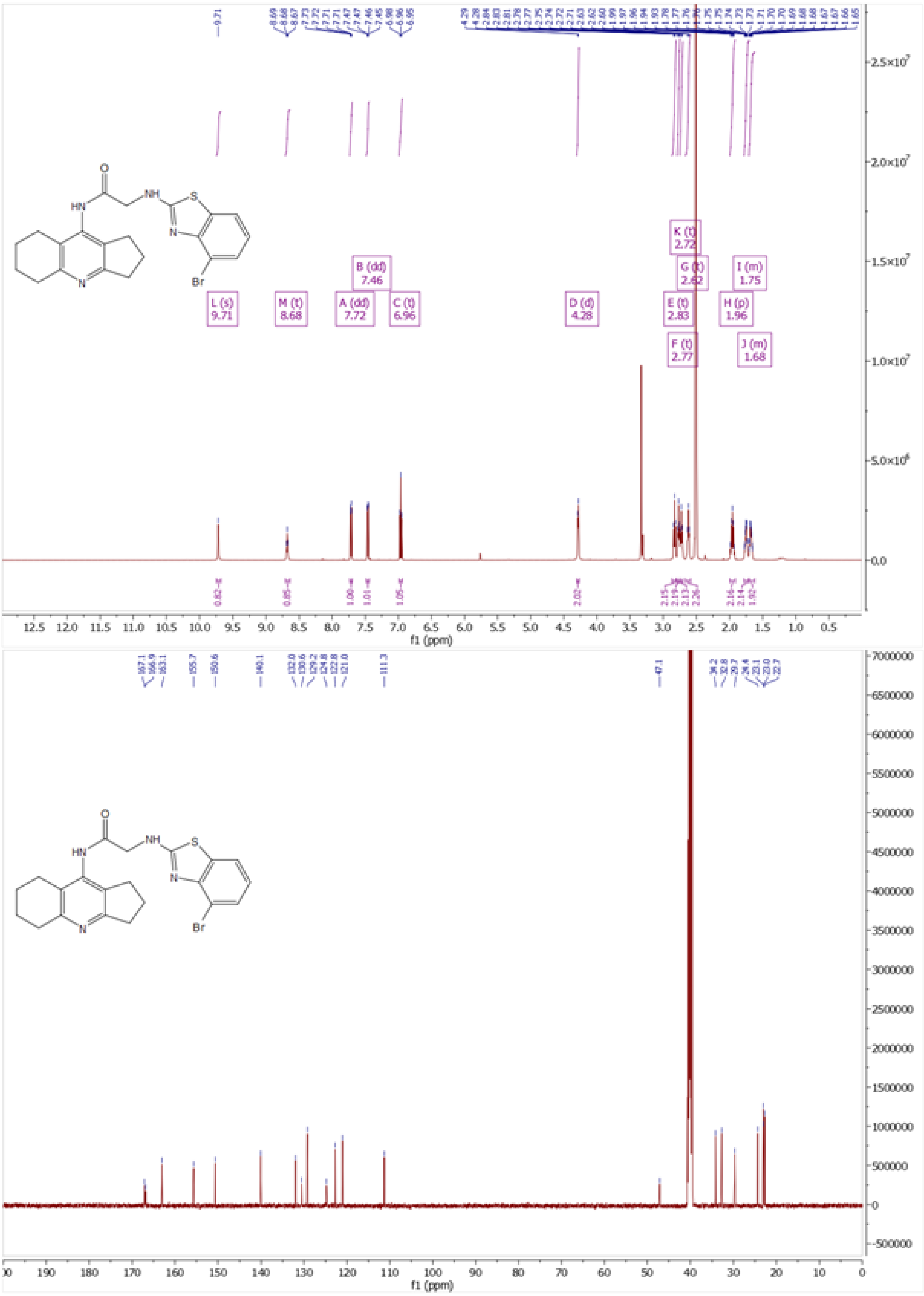

*Figure S16.* [1]*H and* [13]*C NMR spectra of* ***7d***

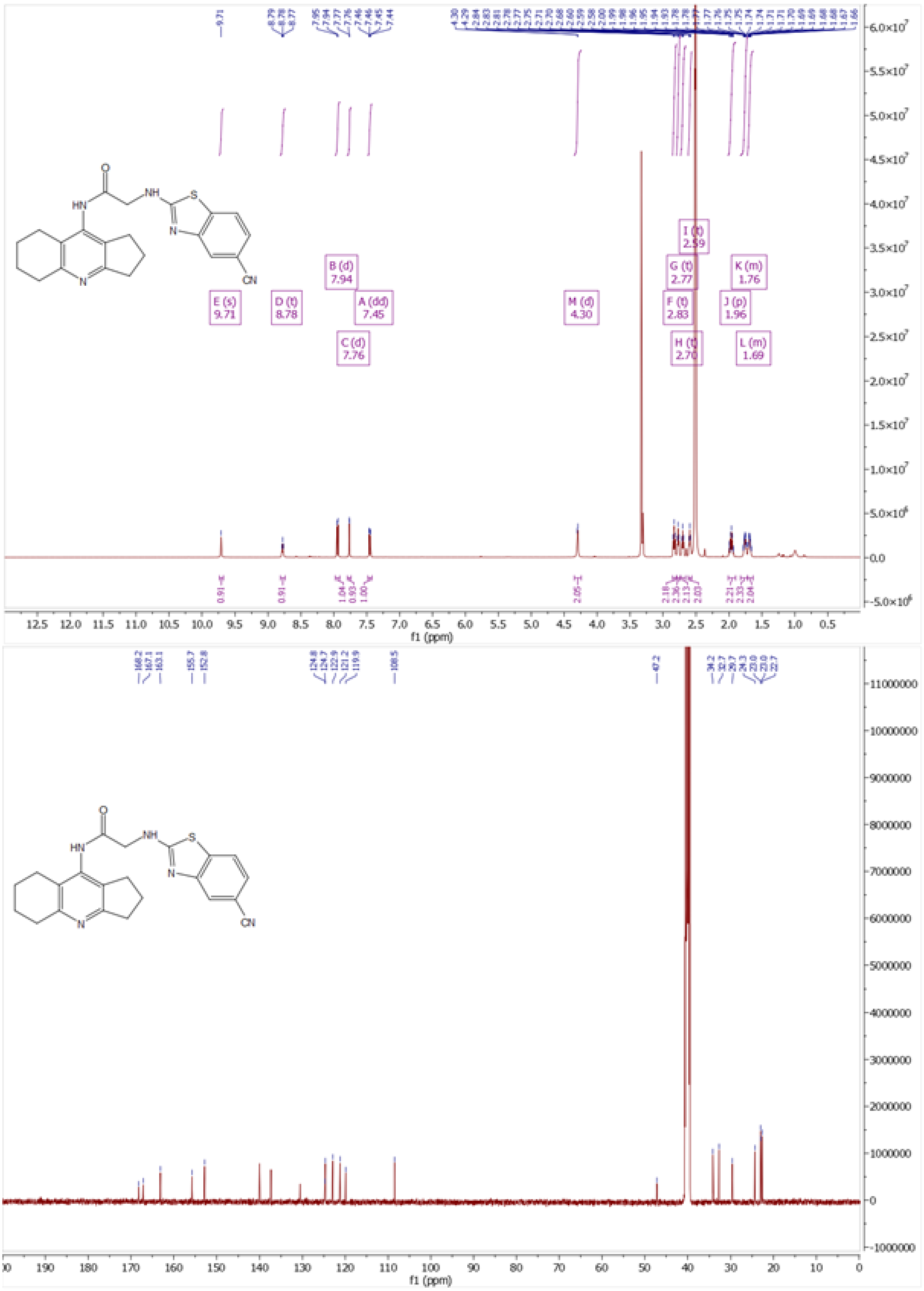

*Figure S17. [1]H and [13]C NMR spectra of **7e***

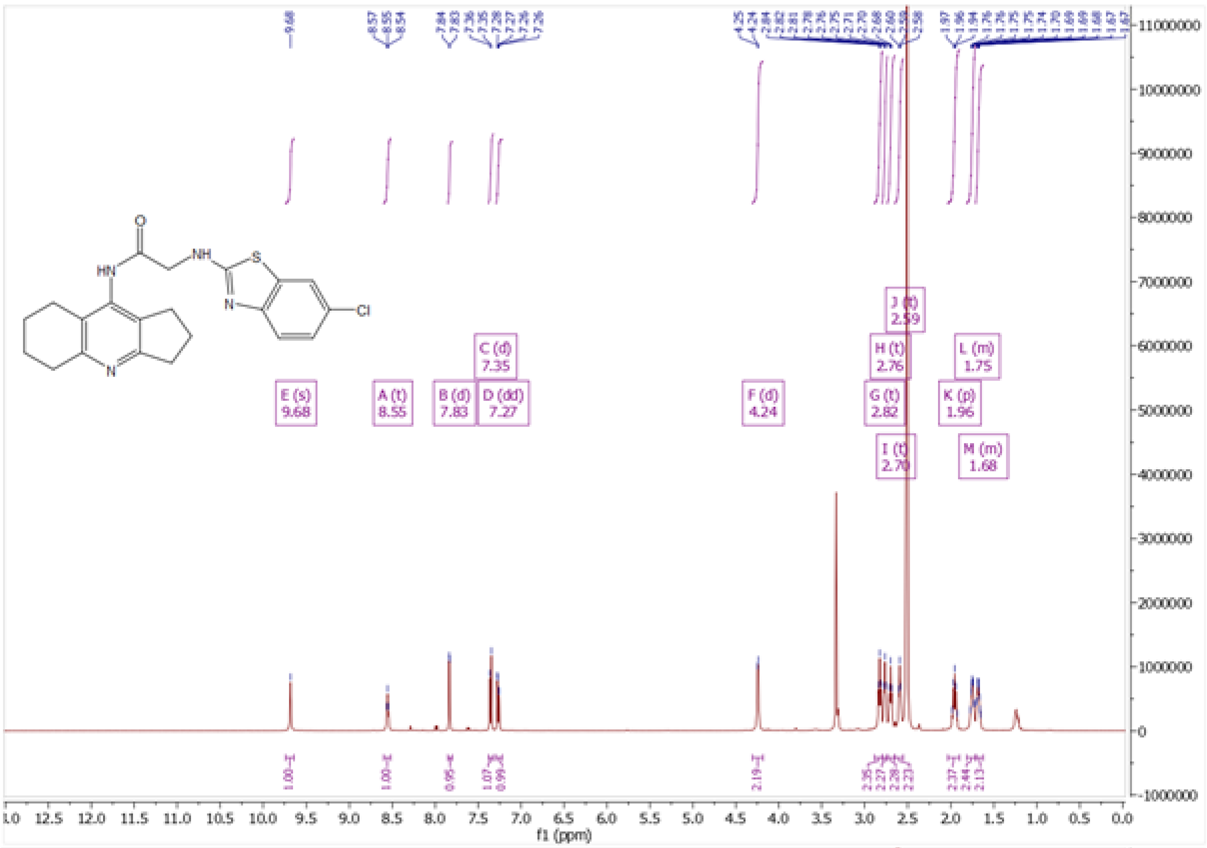

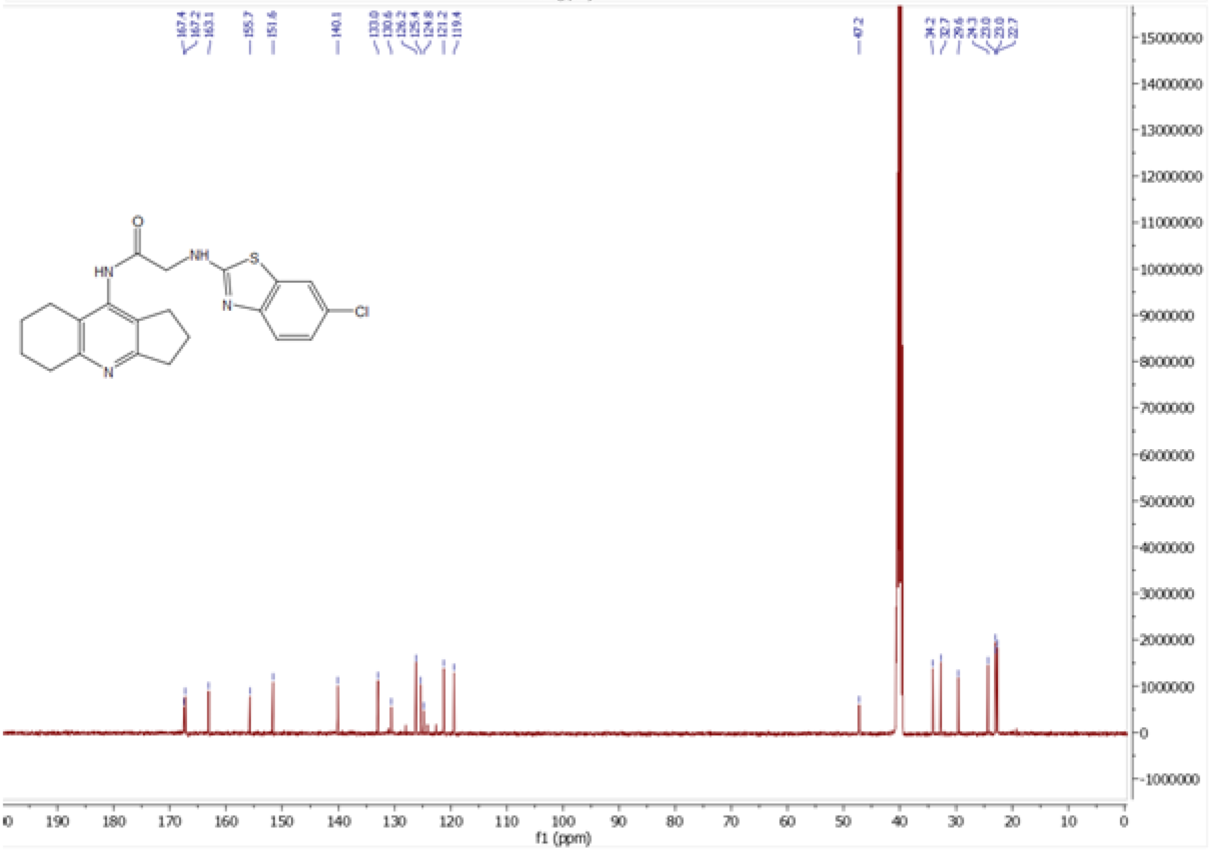

*Figure S18.* $^1$*H and* $^{13}$*C NMR spectra of* ***7f***

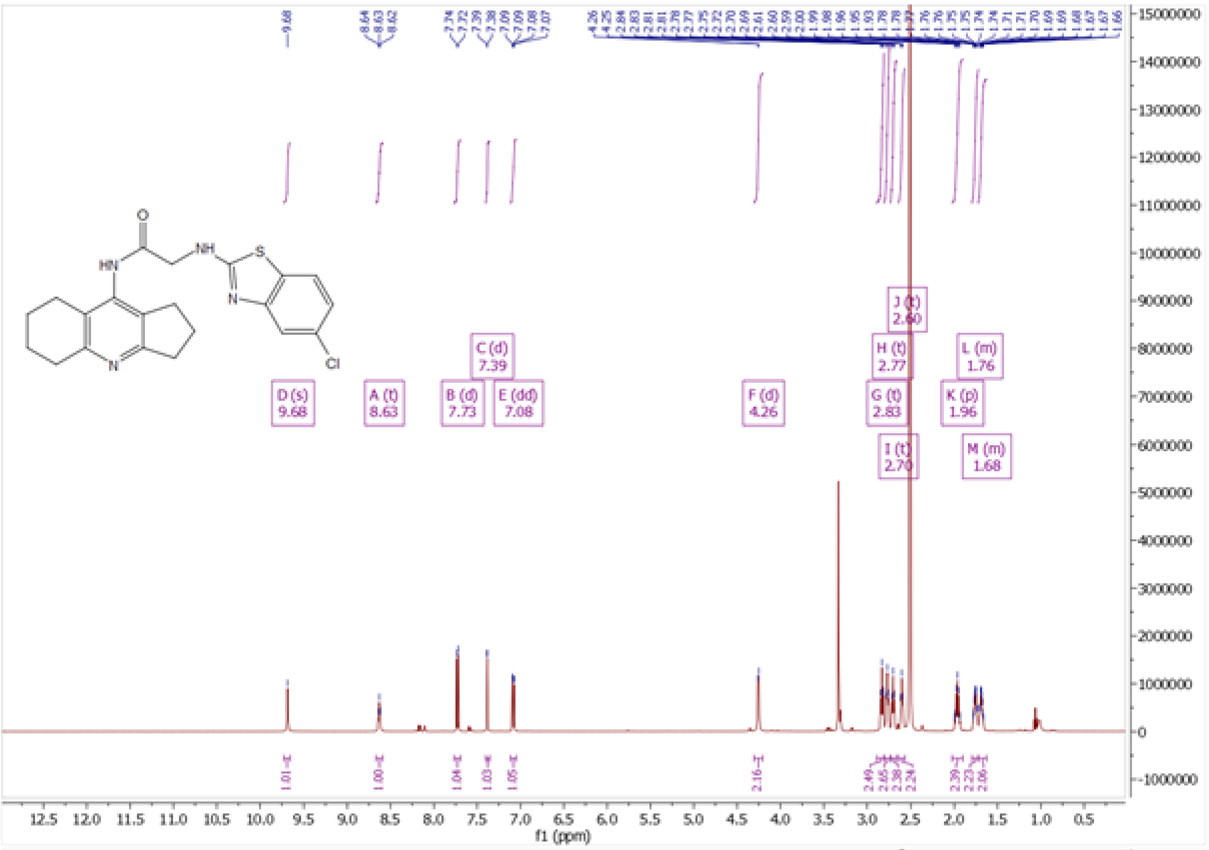

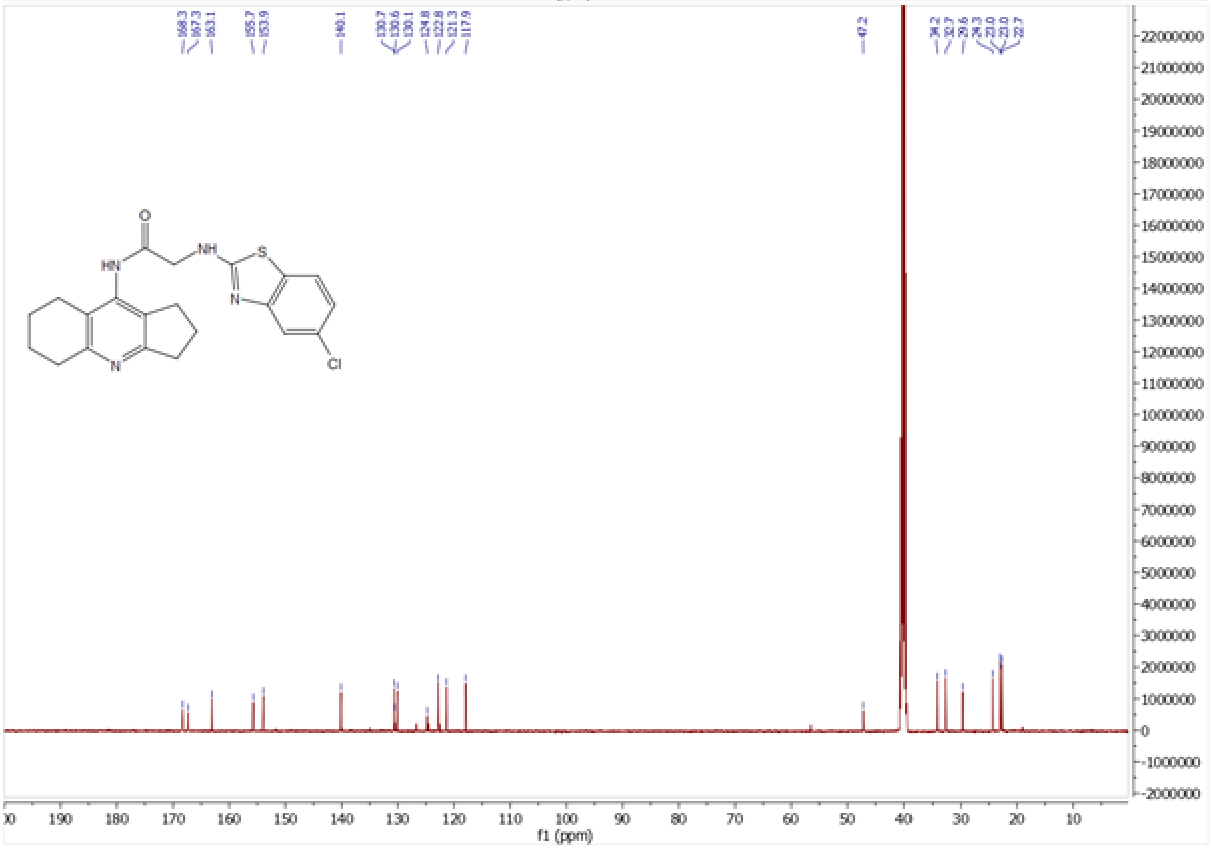

*Figure S19. [1]H and [13]C NMR spectra of **7g***

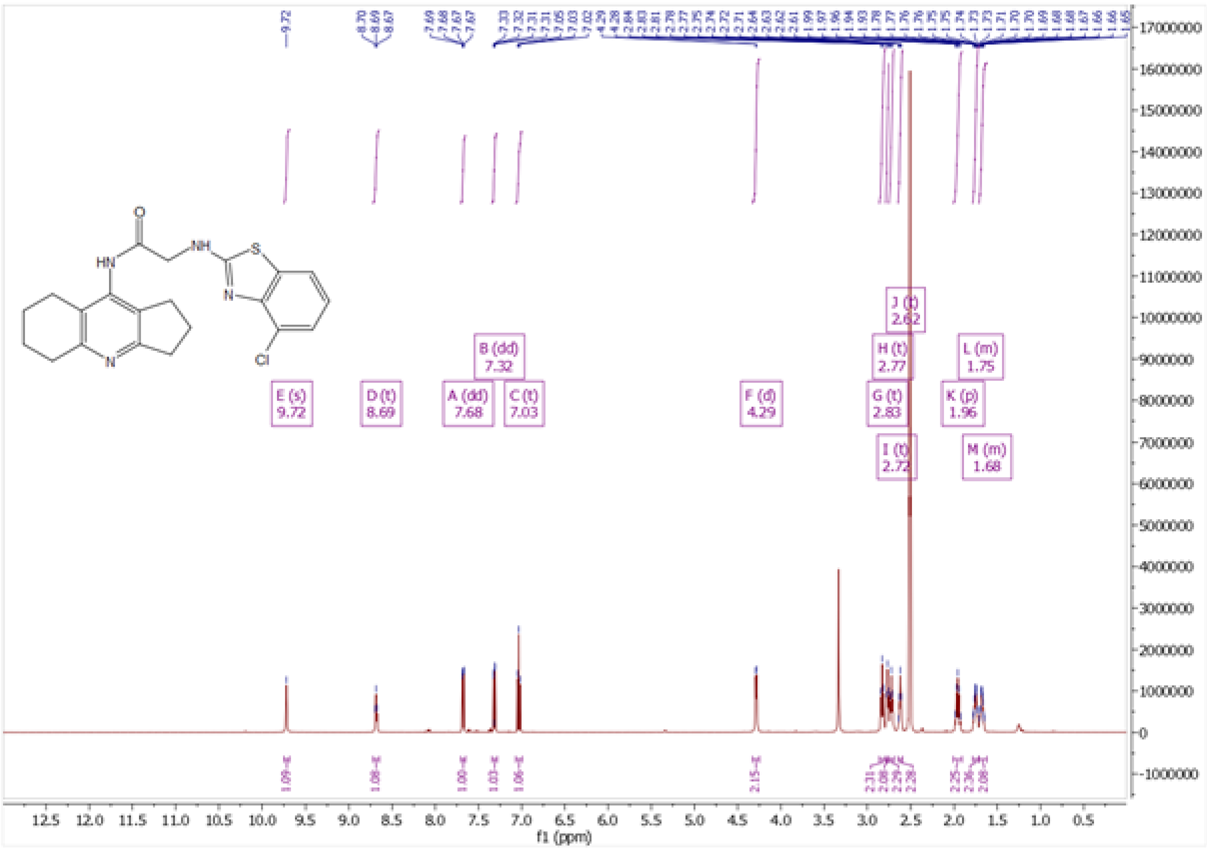

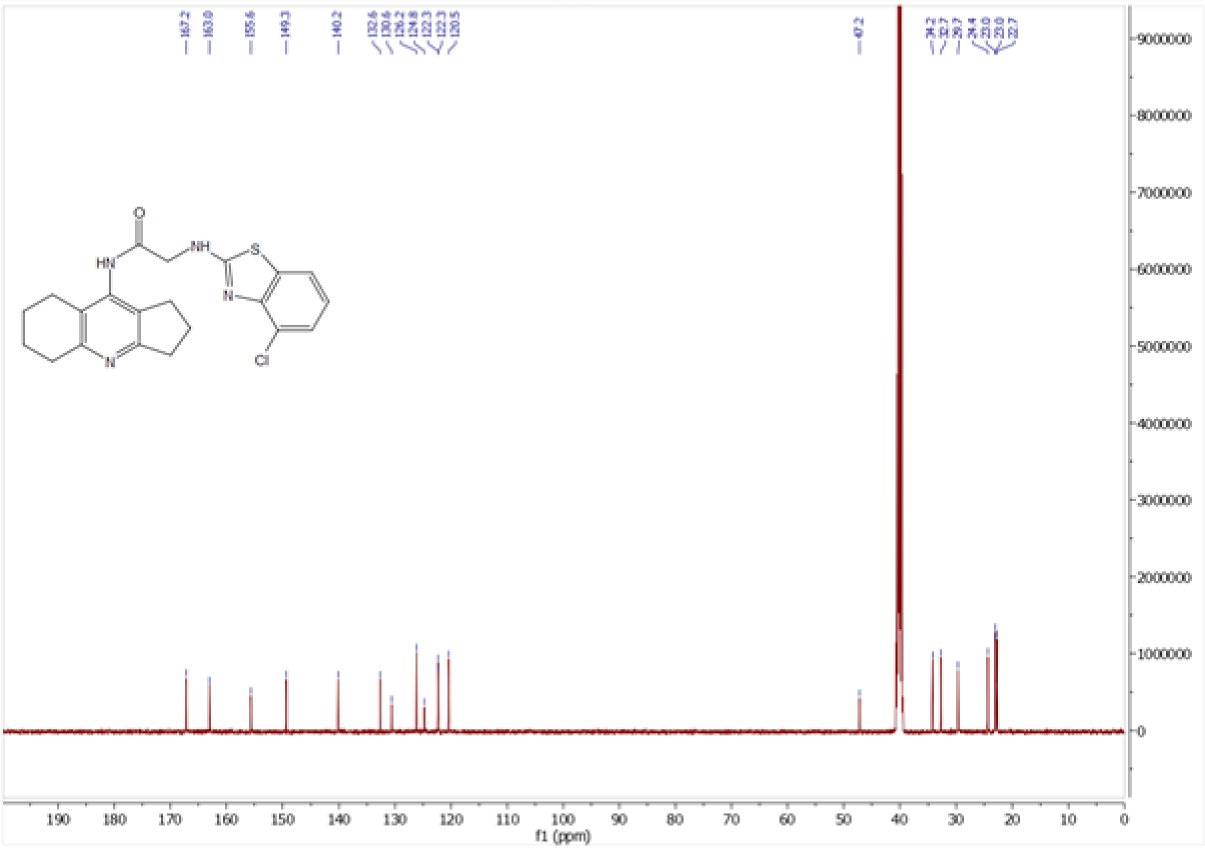

*Figure S20. $^{1}H$ and $^{13}C$ NMR spectra of **7h***

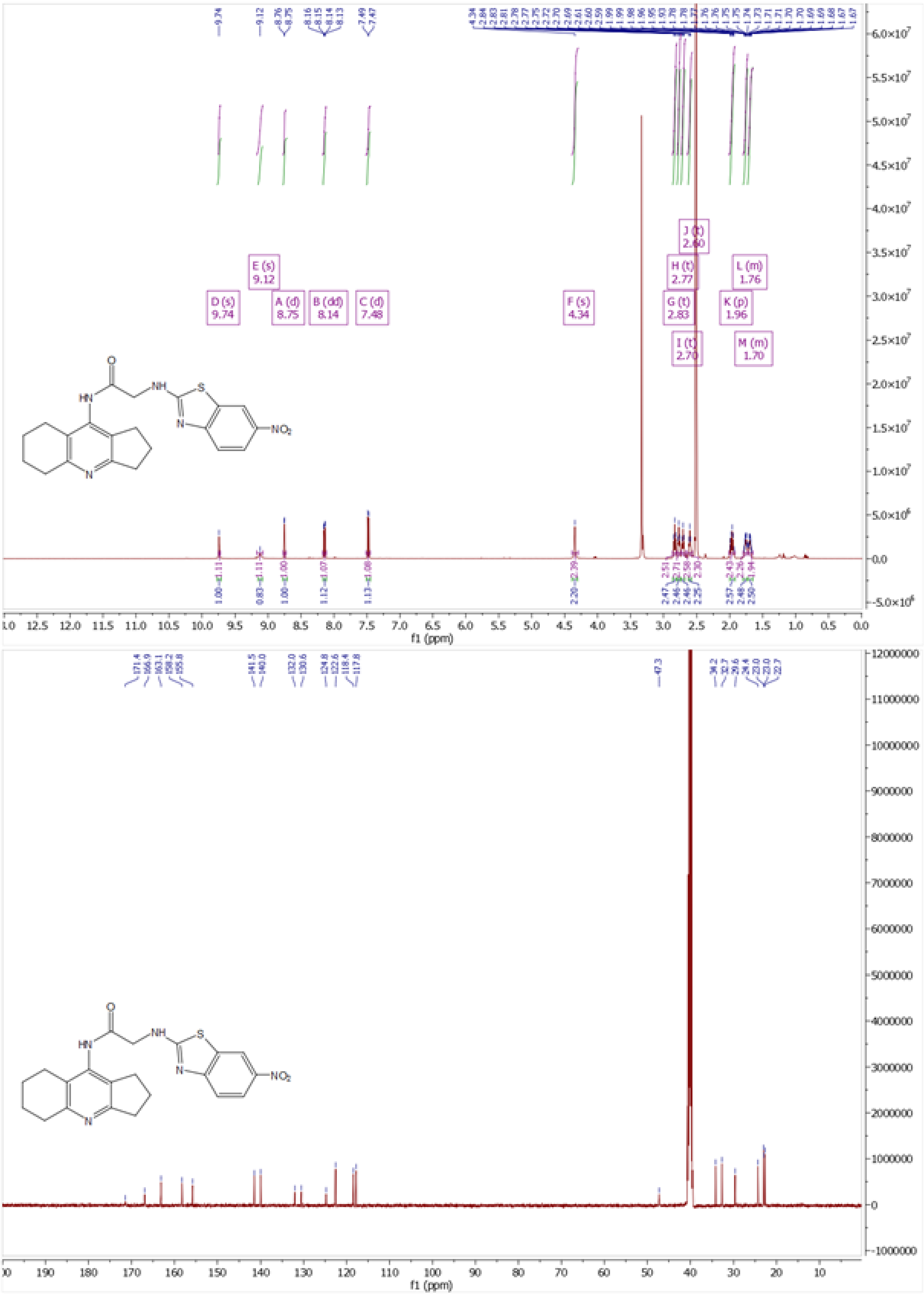

*Figure S21. $^{1}H$ and $^{13}C$ NMR spectra of **7i***

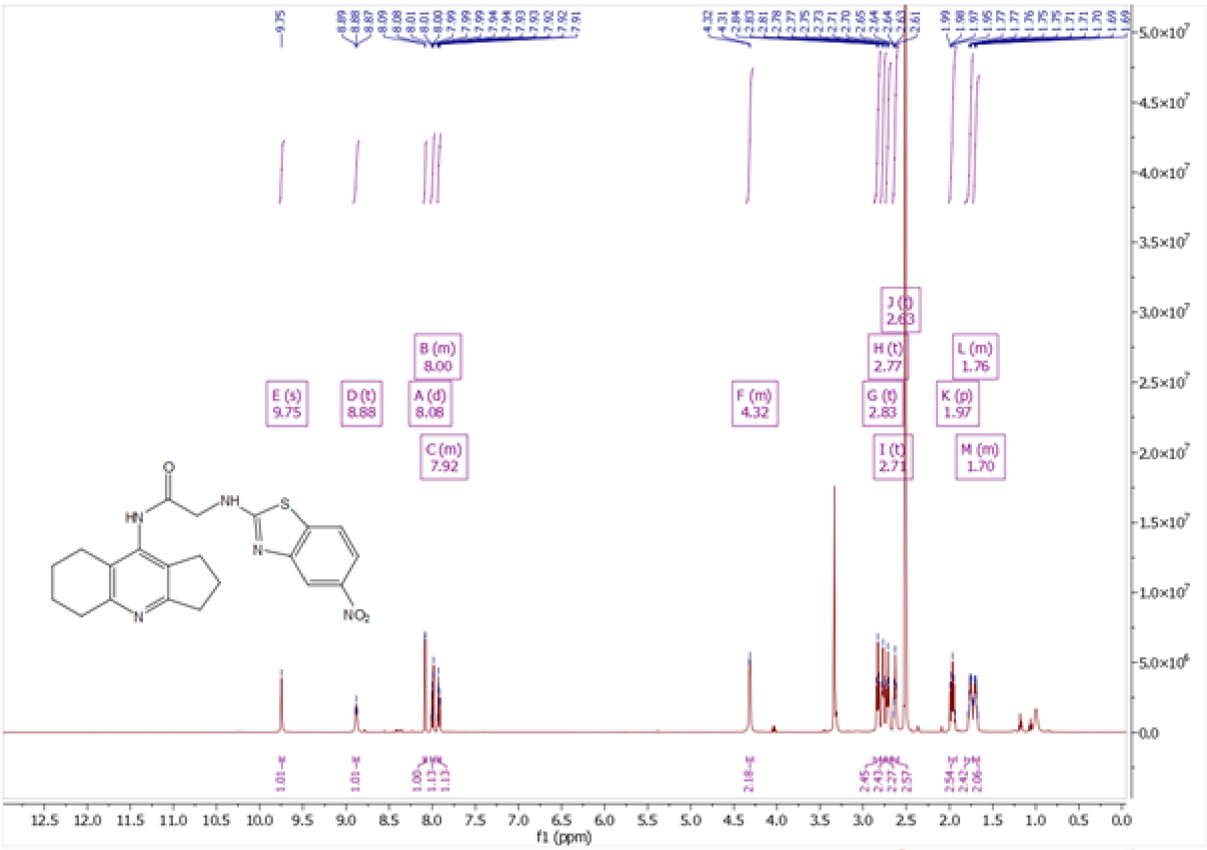

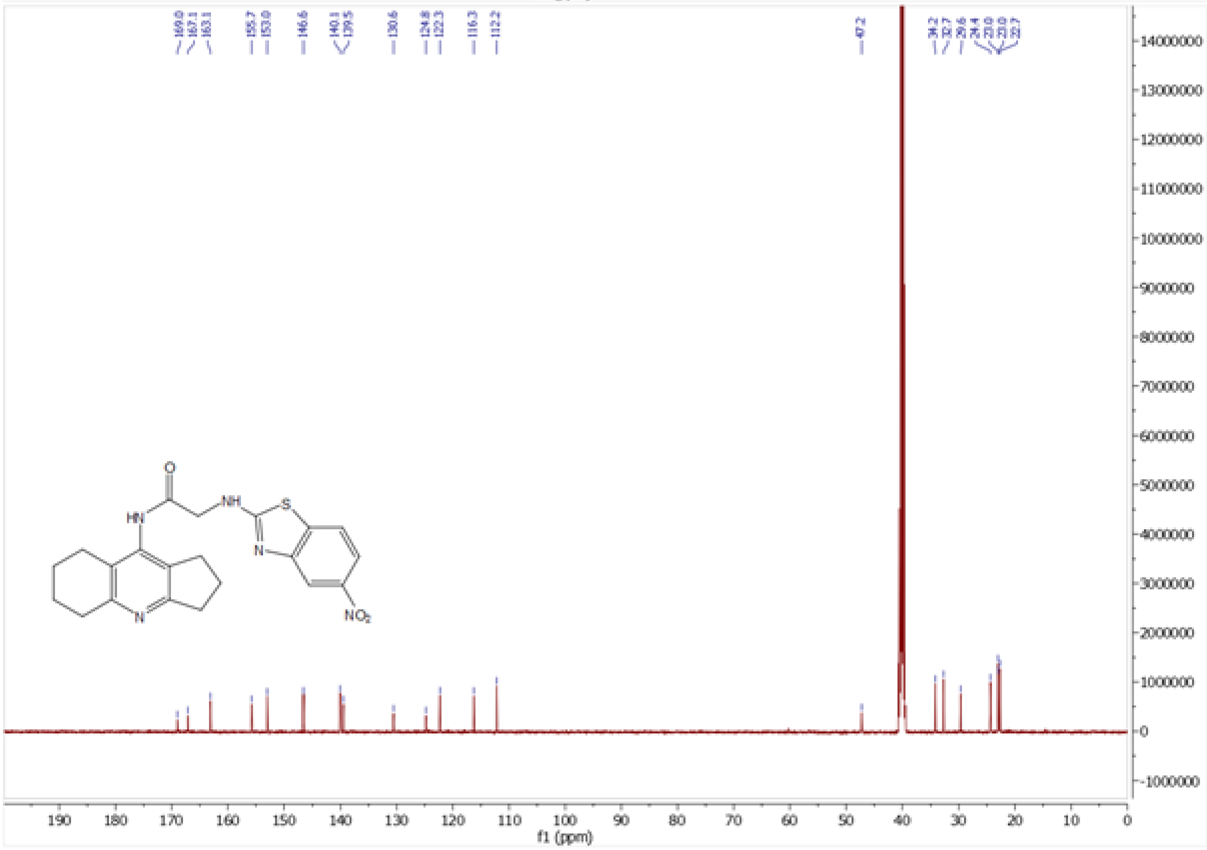

*Figure S22. $^{1}H$ and $^{13}C$ NMR spectra of* ***7j***

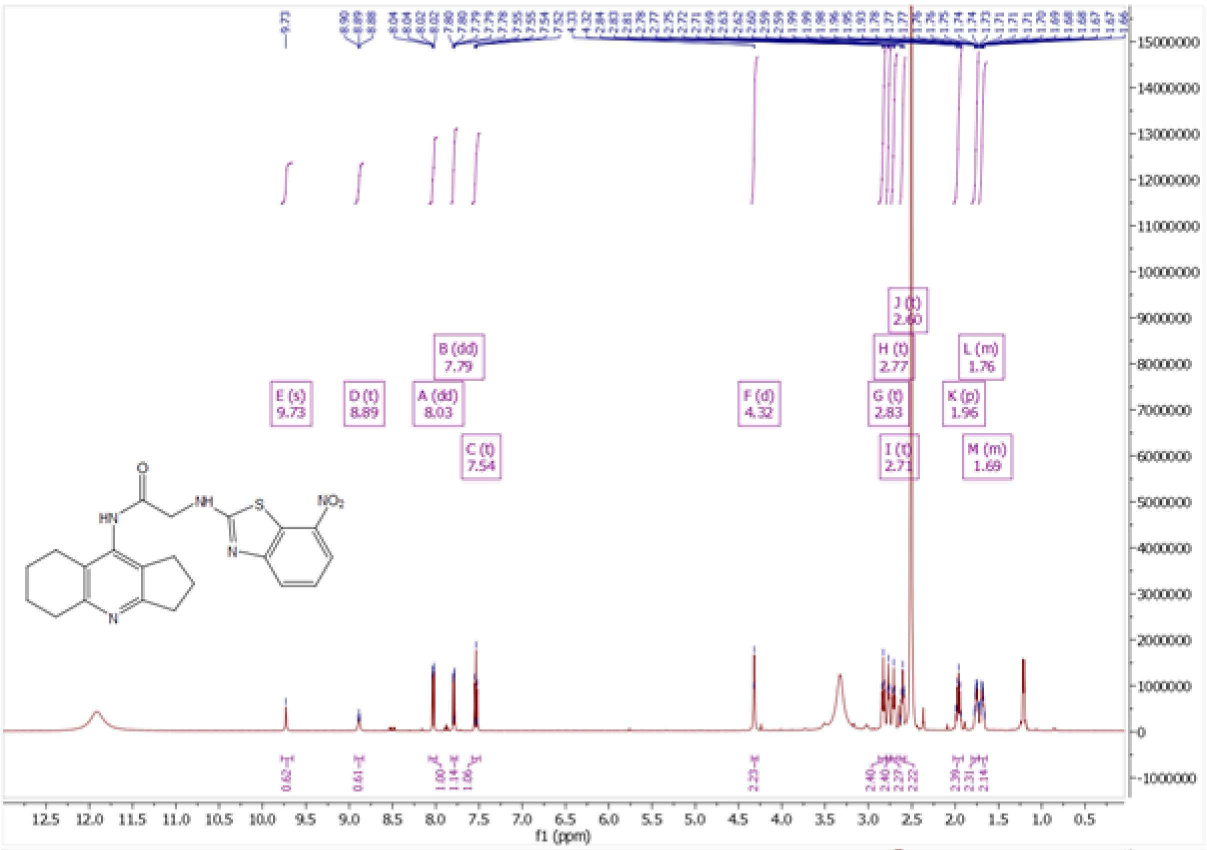

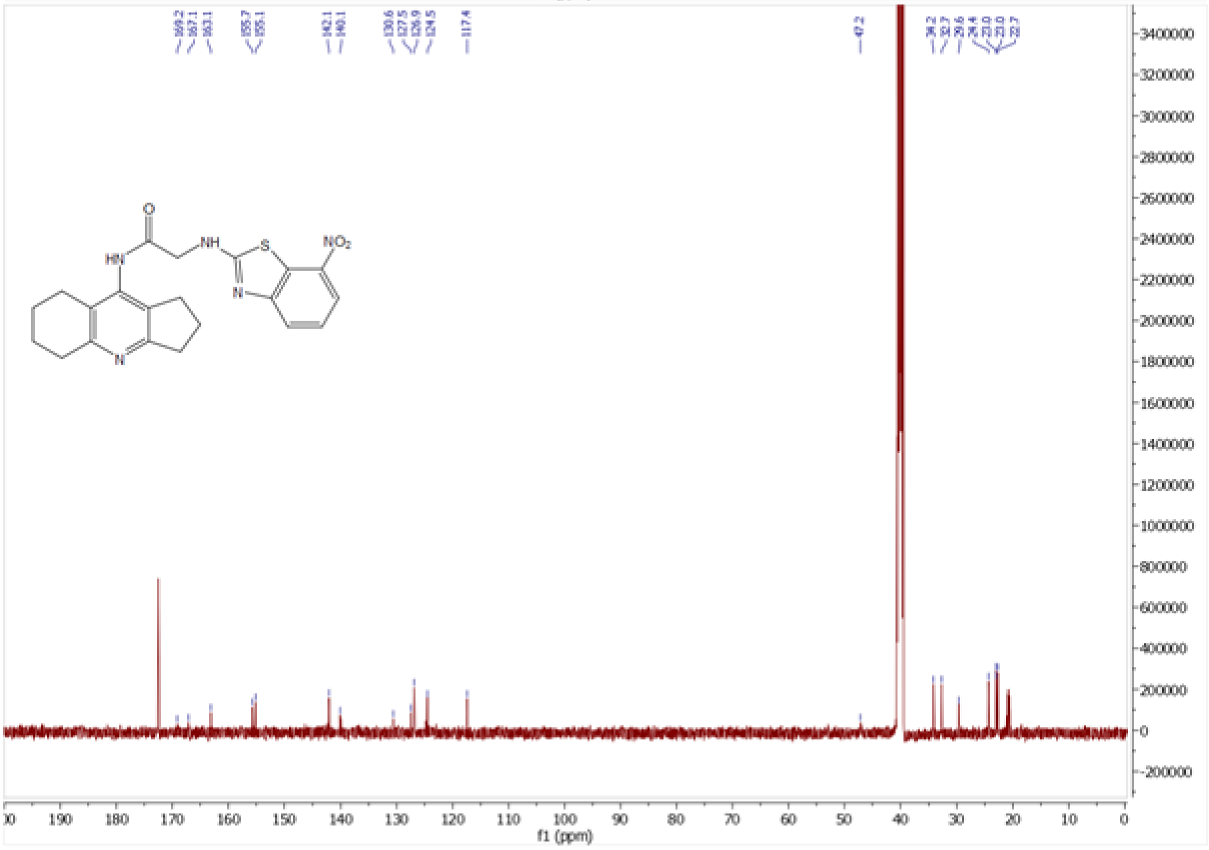

*Figure S23. $^{1}$H and $^{13}$C NMR spectra of **7k***

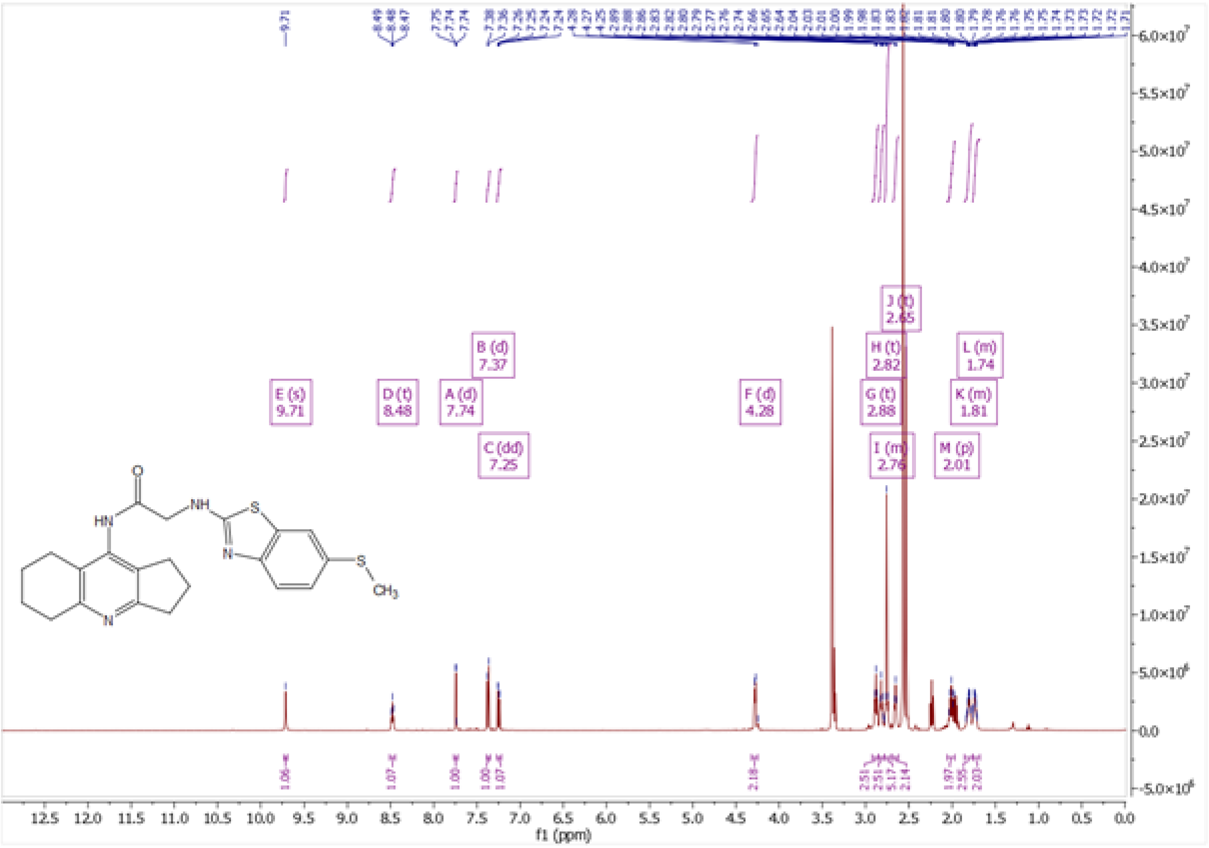

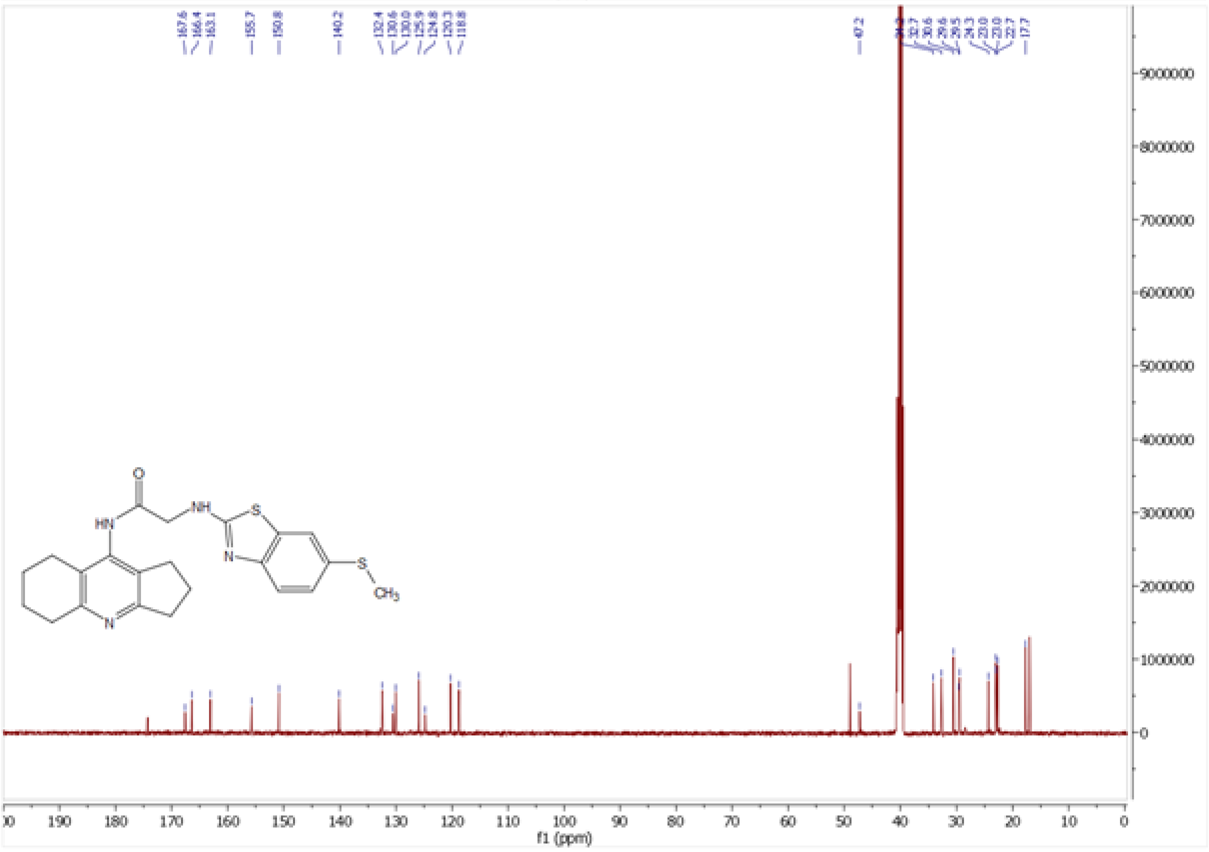

*Figure S24. ¹H and ¹³C NMR spectra of **7l***

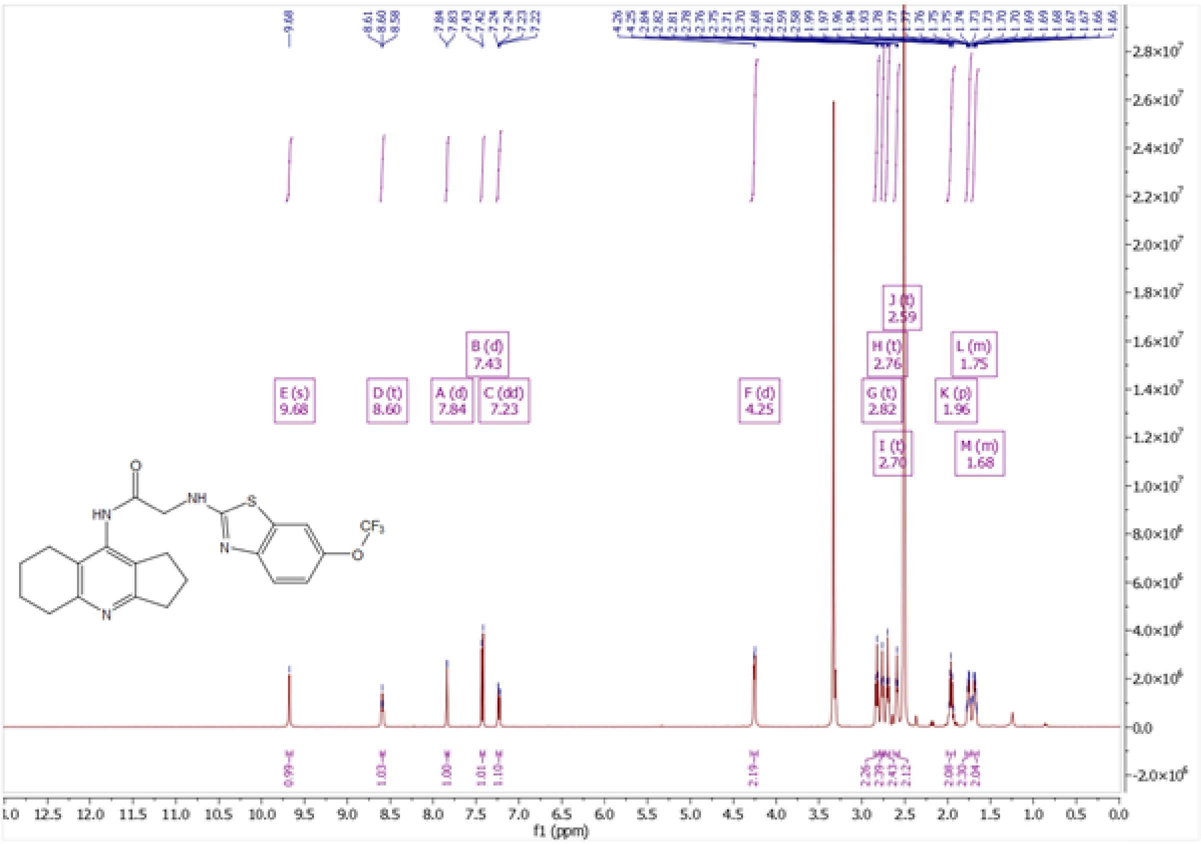

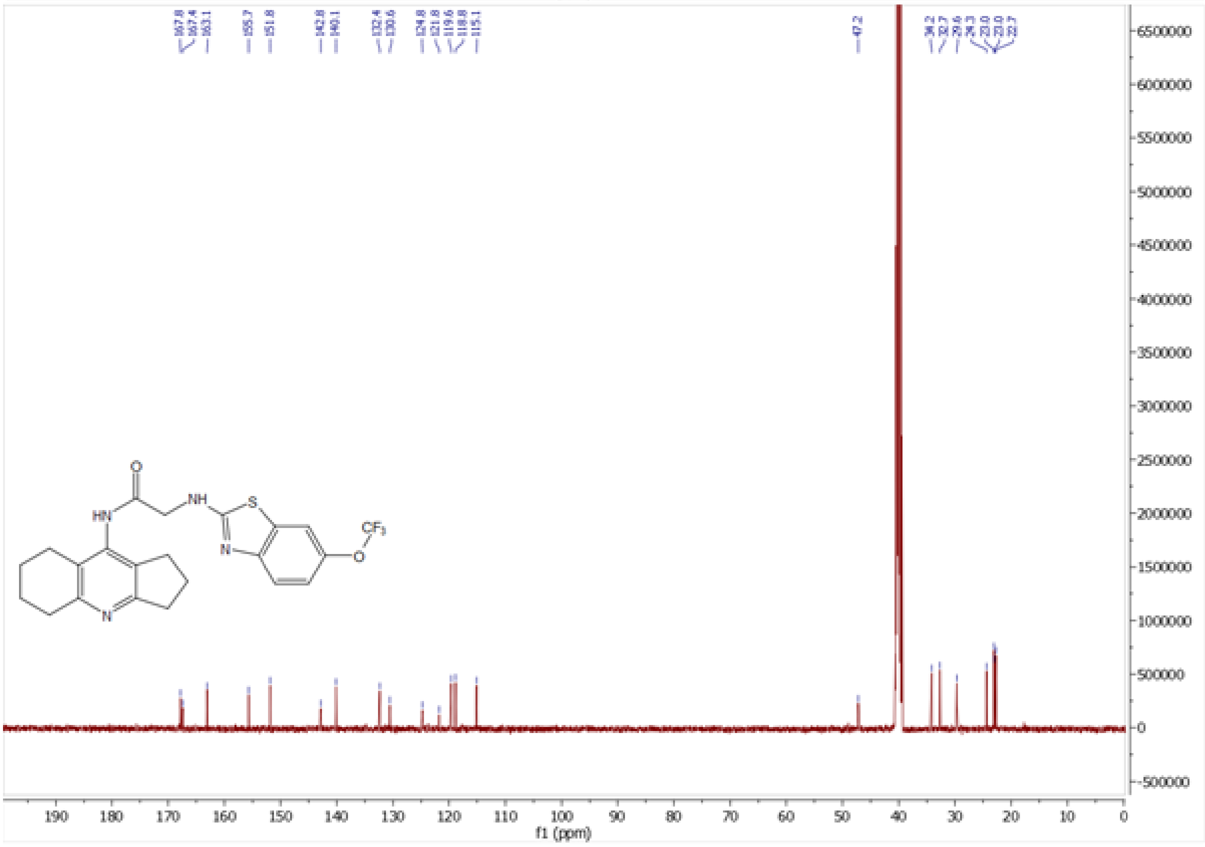

*Figure S25. $^{1}H$ and $^{13}C$ NMR spectra of **7m***

### 8. HPLC-MS spectra of prepared compounds

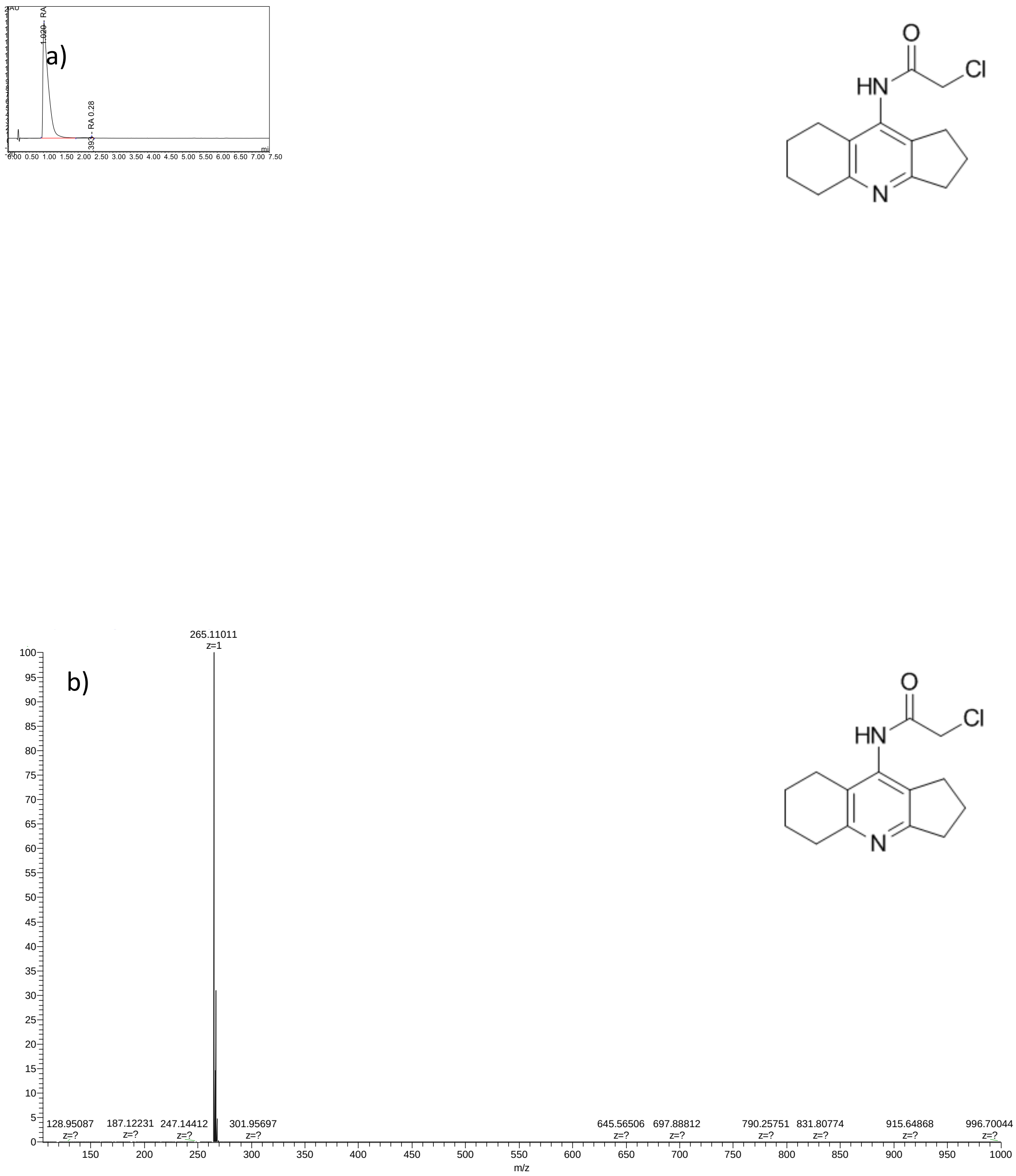

*Figure S26. HPLC-MS of compound **8** a) LC-UV chromatogram b) HRMS mass spectra*

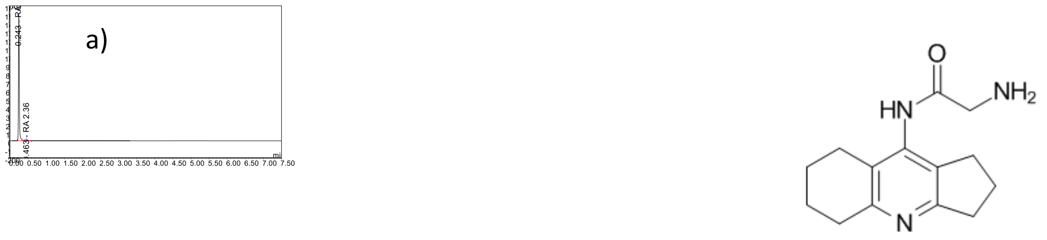

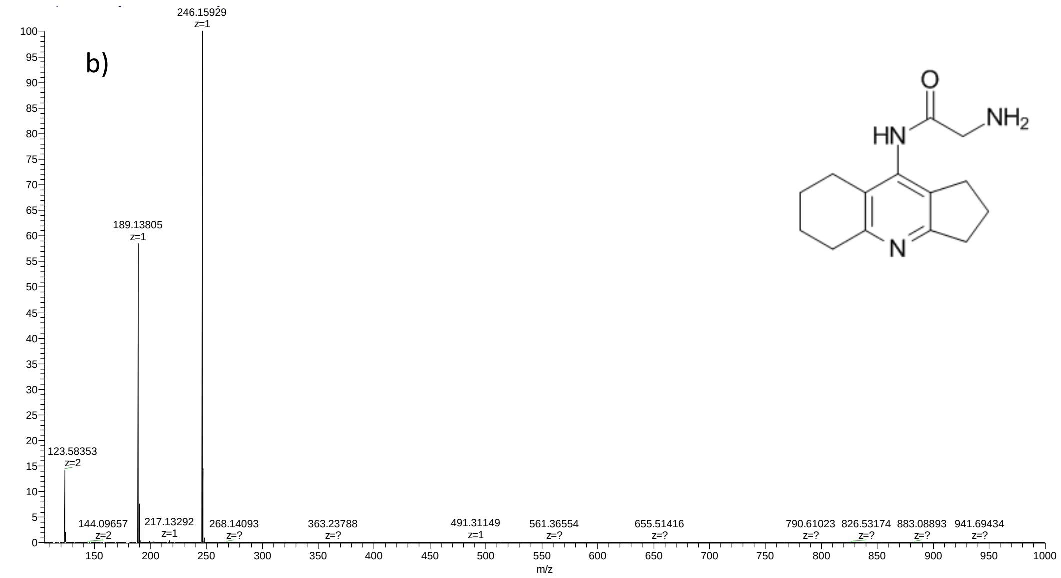

*Figure S27. HPLC-MS of compound **13** a) LC-UV chromatogram b) HRMS mass spectra*

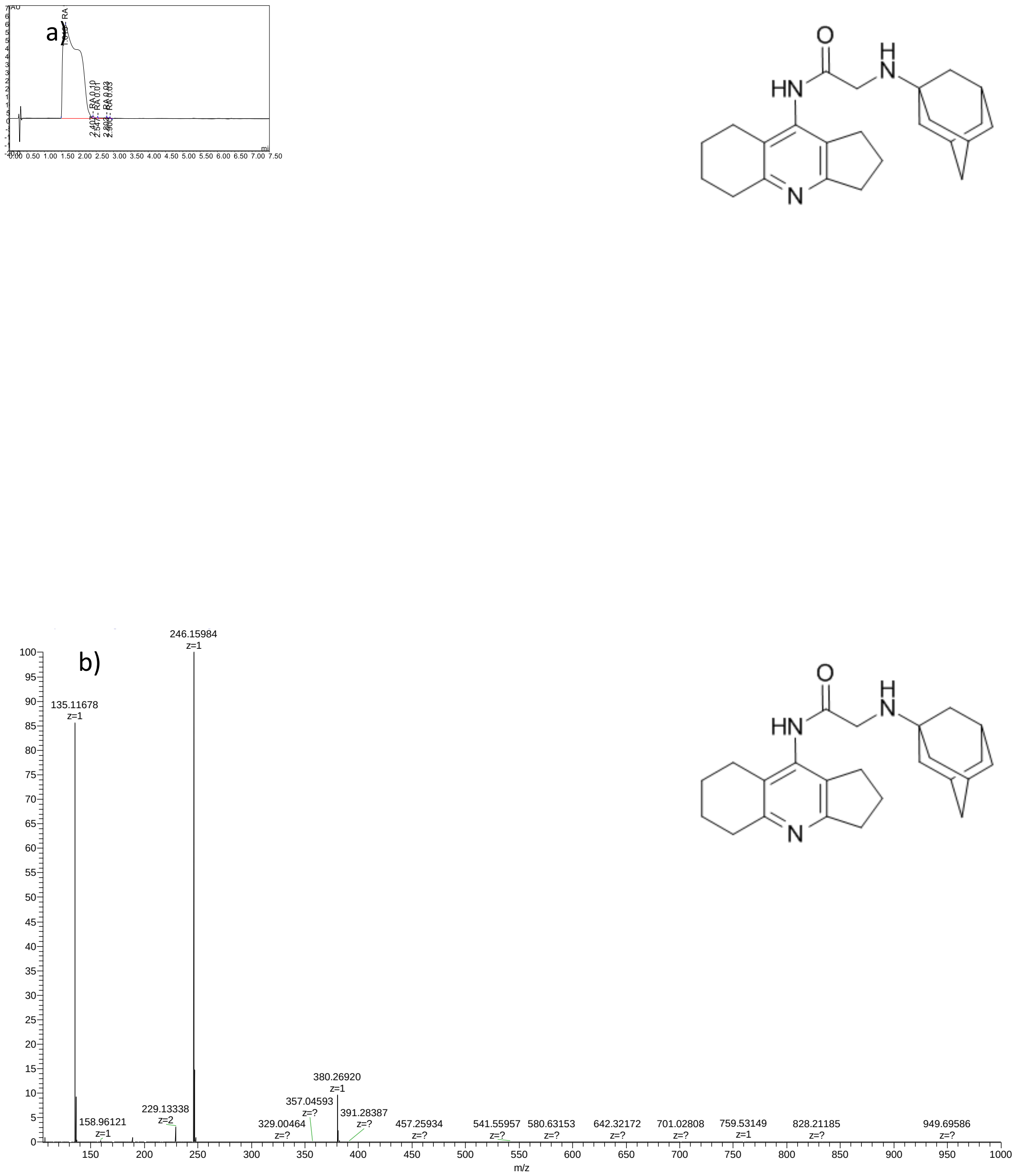

*Figure S28. HPLC-MS of compound **5a** a) LC-UV chromatogram b) HRMS mass spectra*

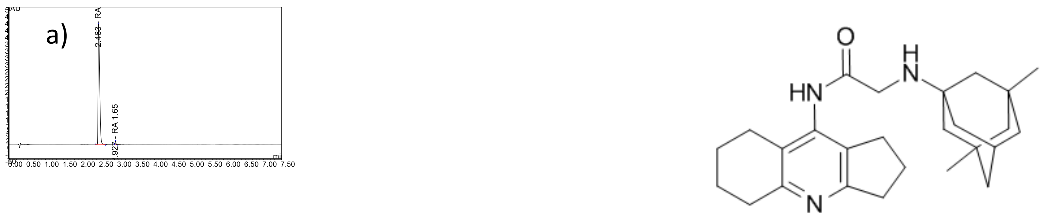

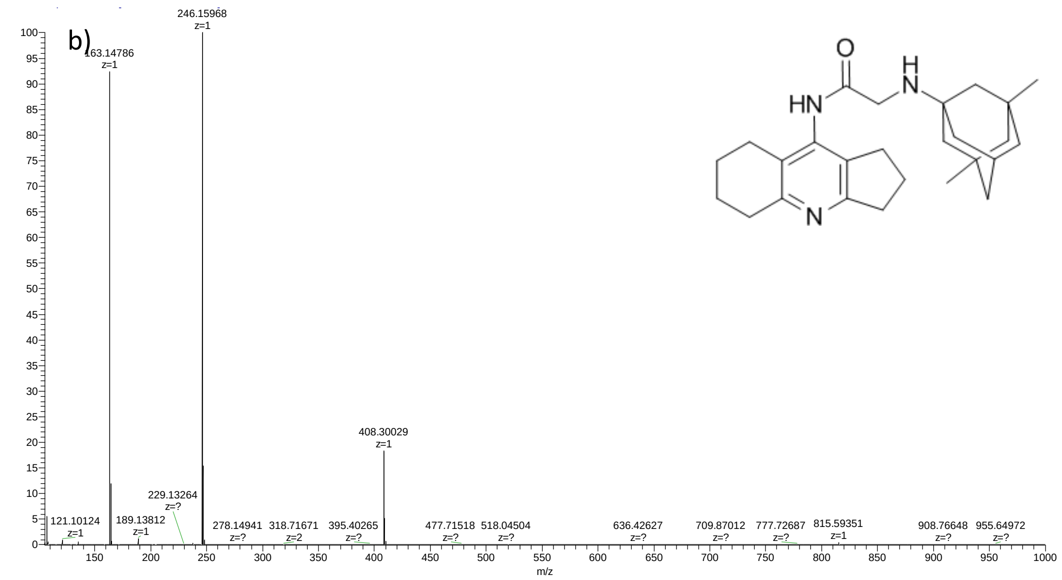

*Figure S 29. HPLC-MS of compound* ***5b*** *a) LC-UV chromatogram b) HRMS mass spectra*

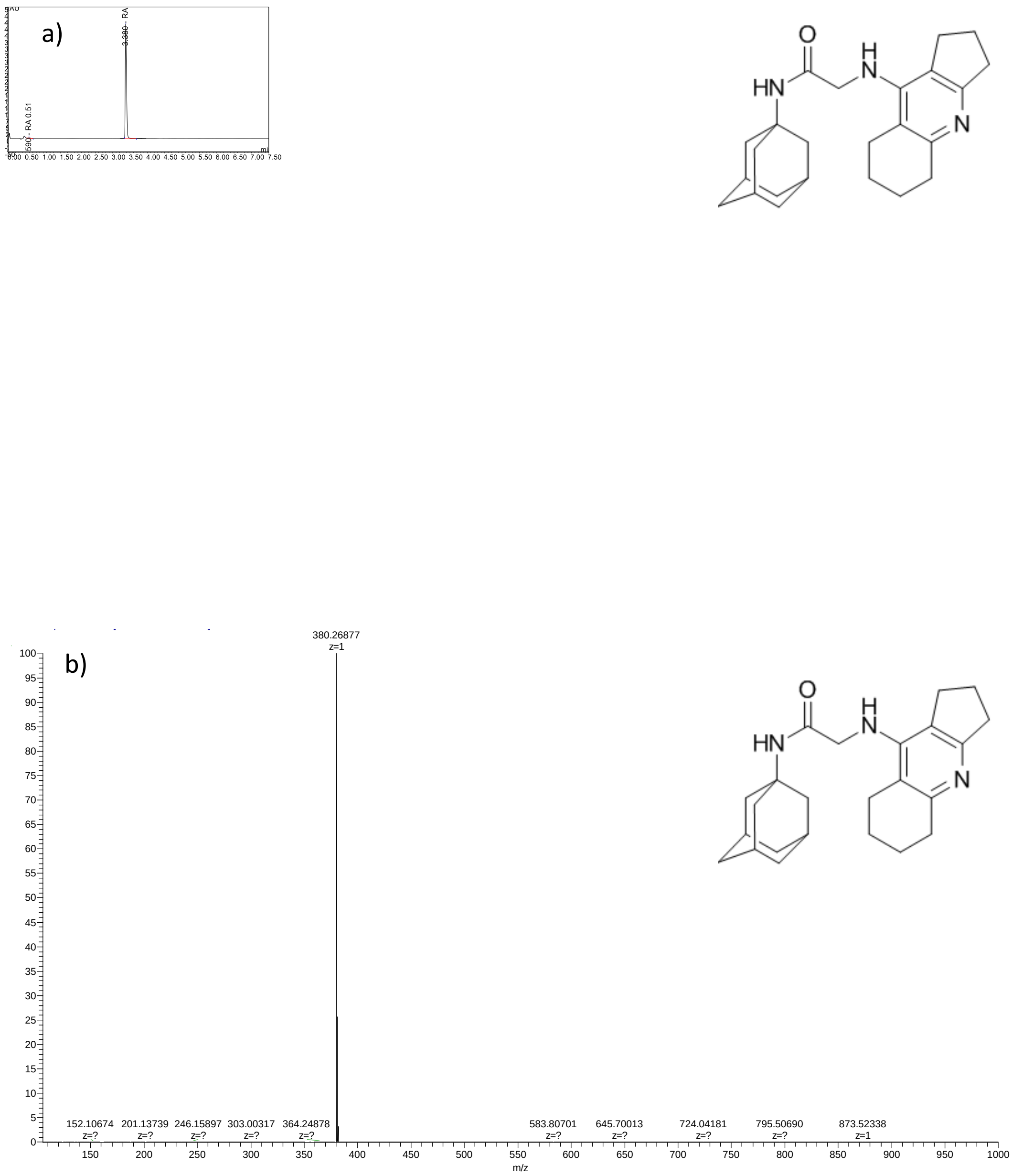

*Figure S 30. HPLC-MS of compound **5c** a) LC-UV chromatogram b) HRMS mass spectra*

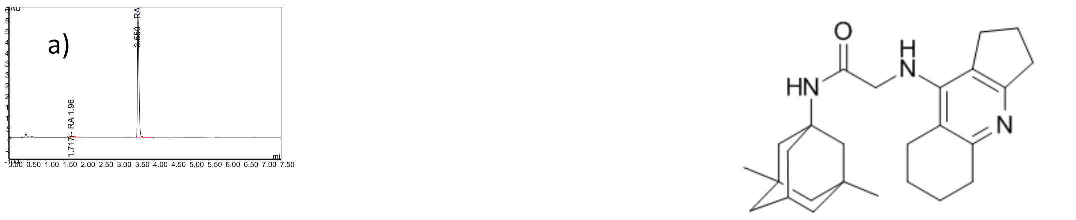

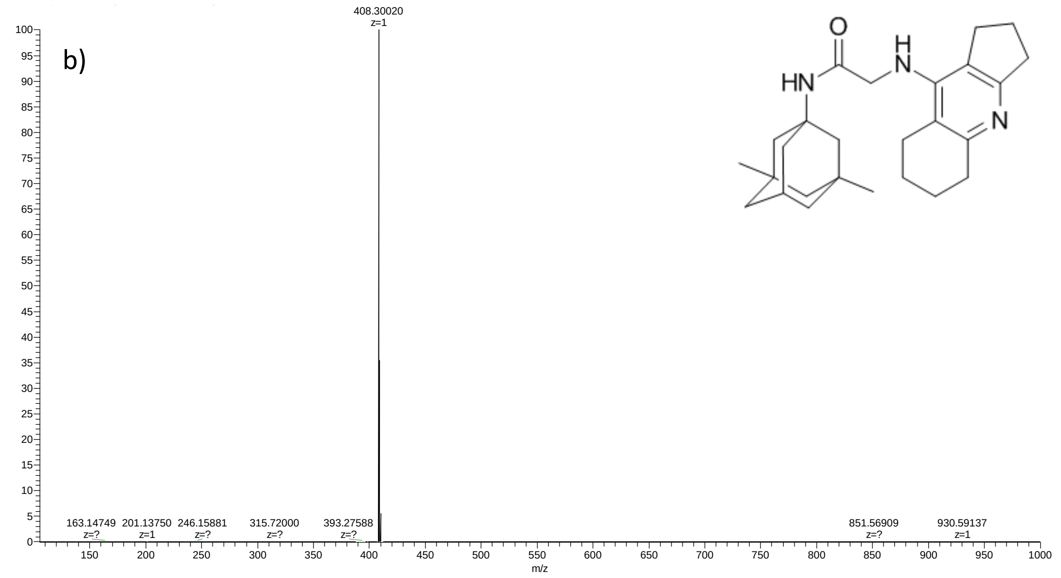

*Figure S 31. HPLC-MS of compound **5d** a) LC-UV chromatogram b) HRMS mass spectra*

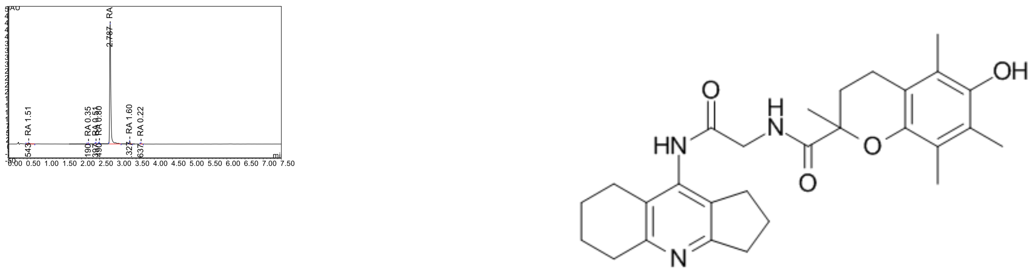

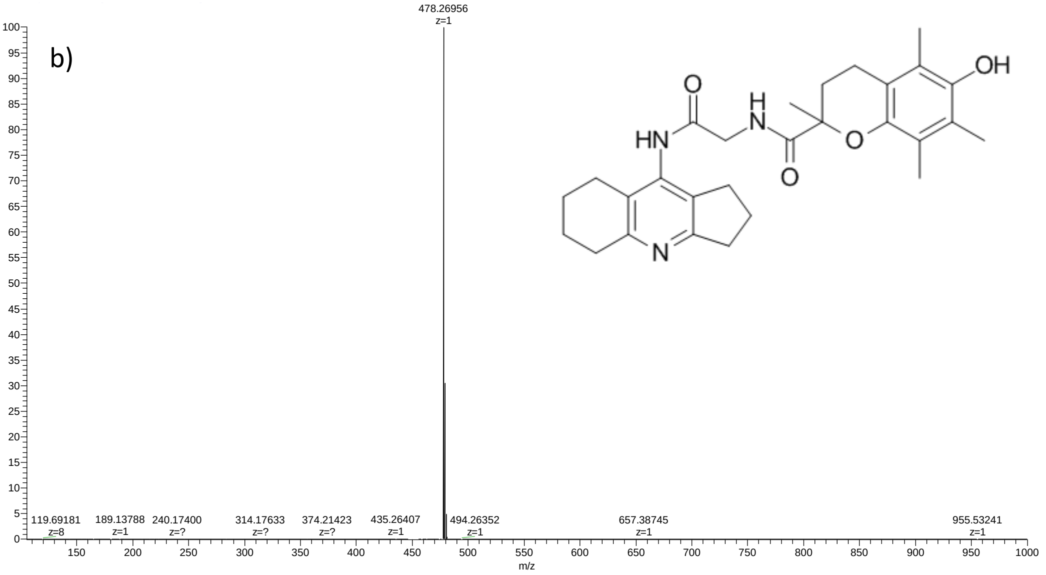

*Figure S 32. HPLC-MS of compound **6** a) LC-UV chromatogram b) HRMS mass spectra*

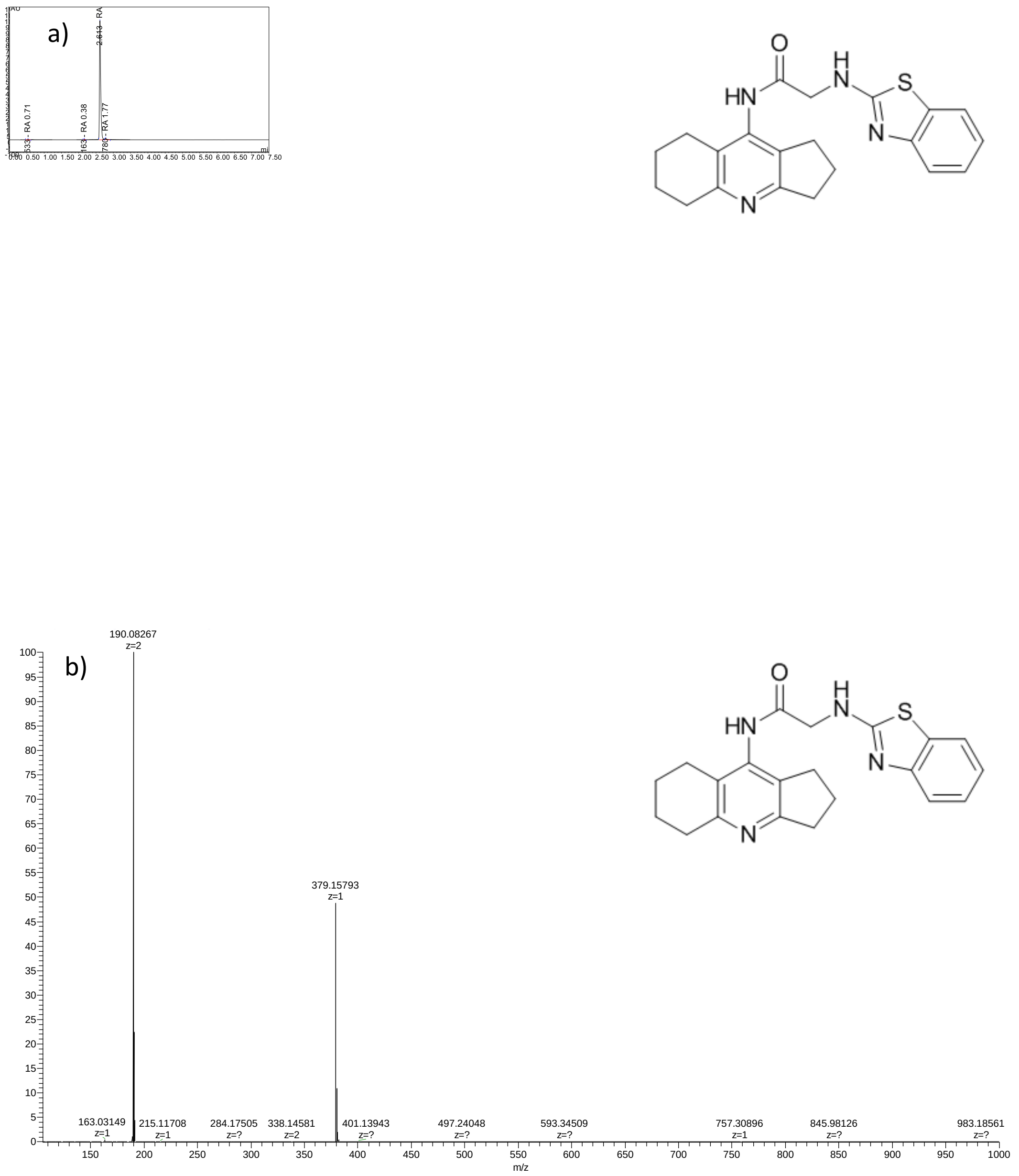

*Figure S 33. HPLC-MS of compound **7a** a) LC-UV chromatogram b) HRMS mass spectra*

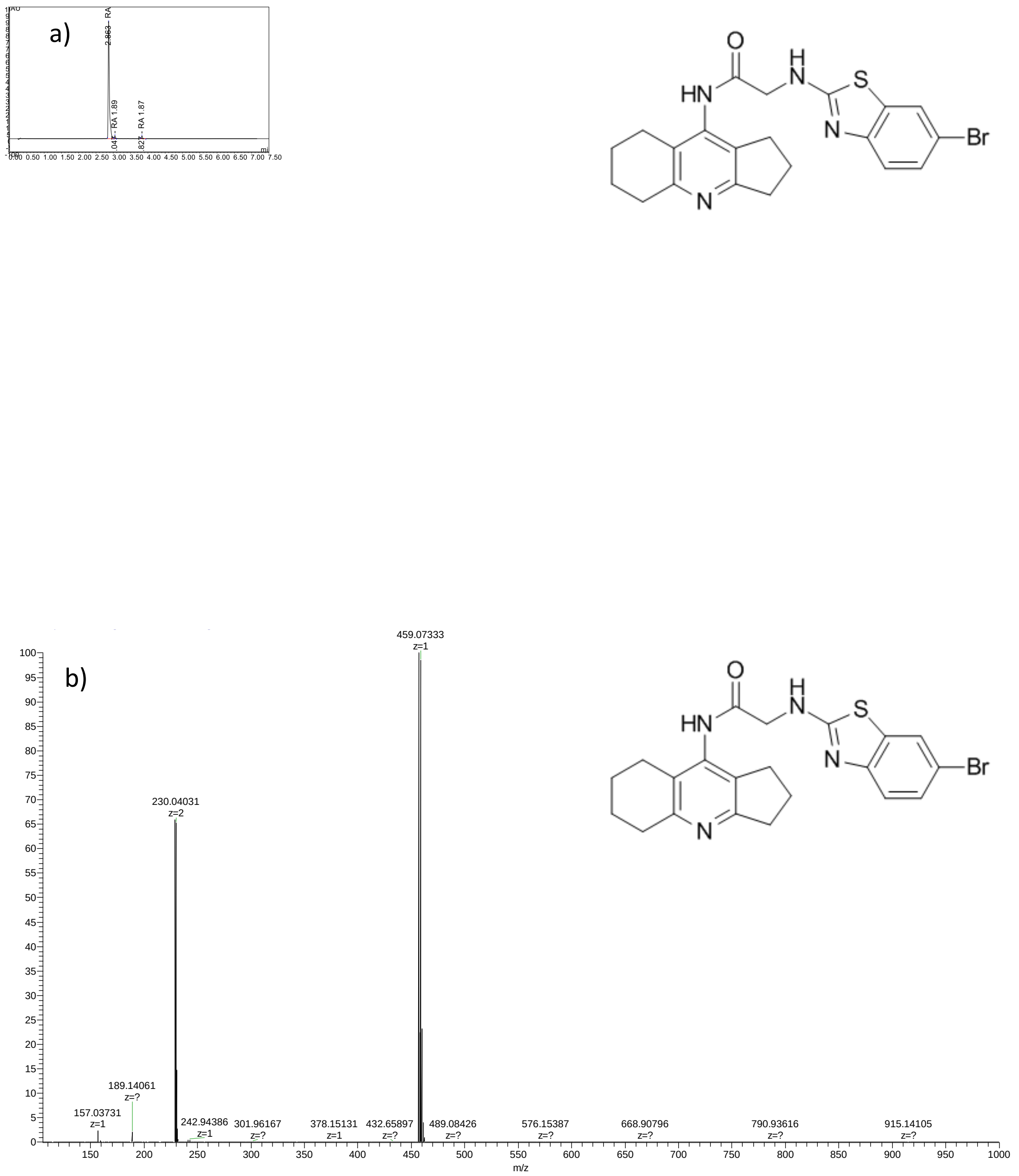

*Figure S 34. HPLC-MS of compound* ***7b*** *a) LC-UV chromatogram b) HRMS mass spectra*

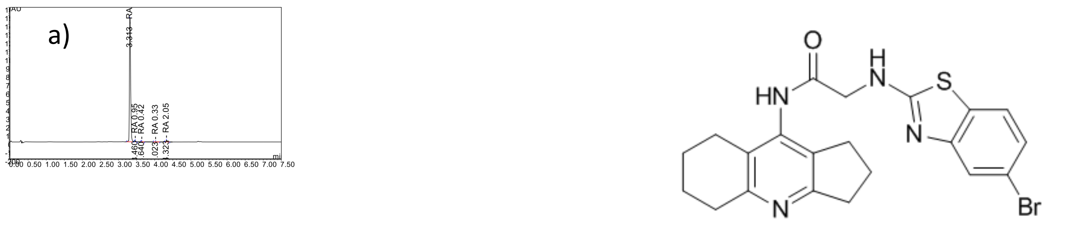

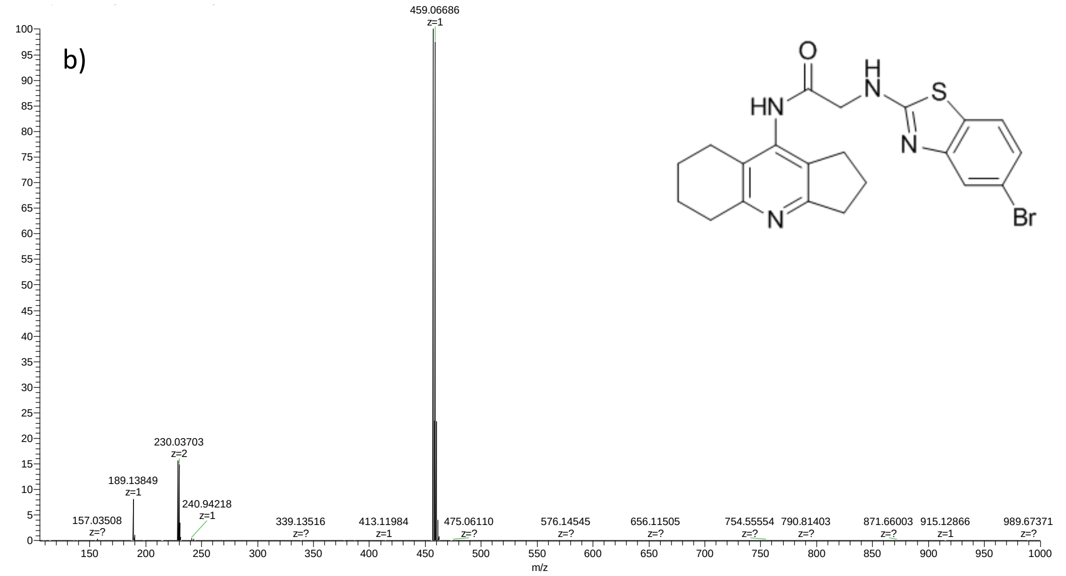

*Figure S 35. HPLC-MS of compound **7c** a) LC-UV chromatogram b) HRMS mass spectra*

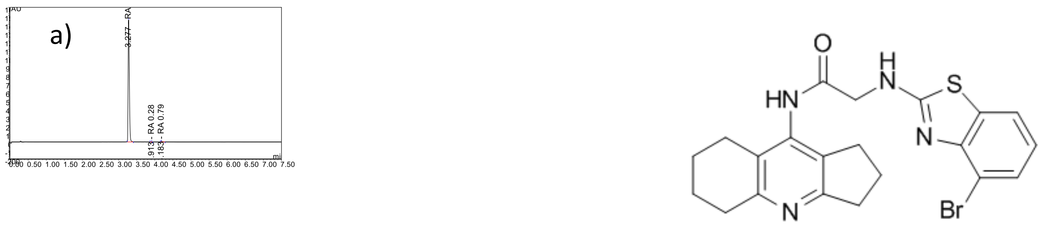

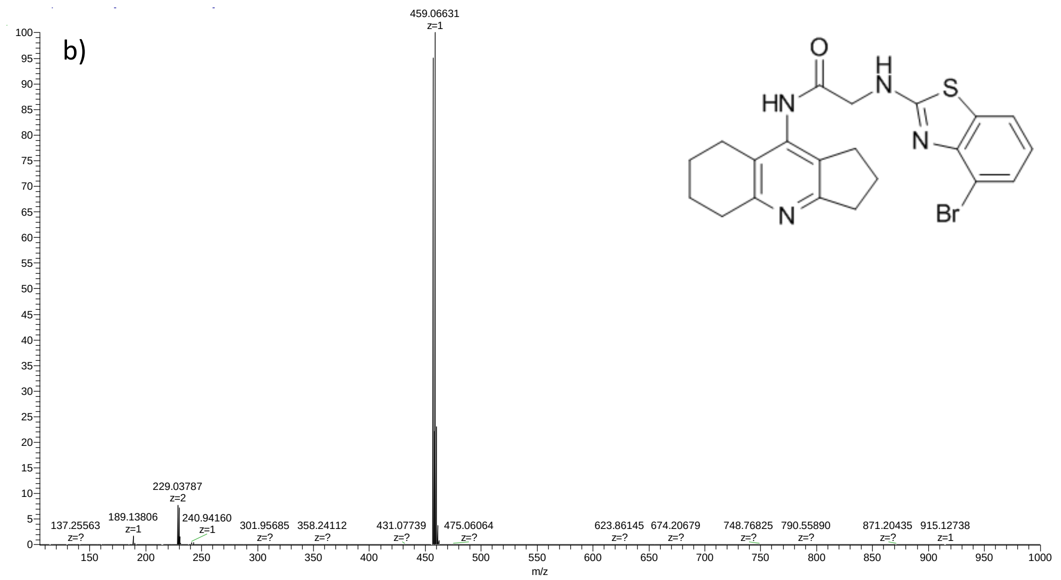

*Figure S 36. HPLC-MS of compound **7d** a) LC-UV chromatogram b) HRMS mass spectra*

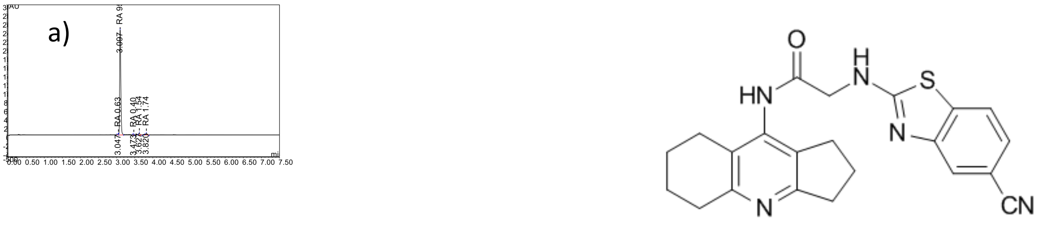

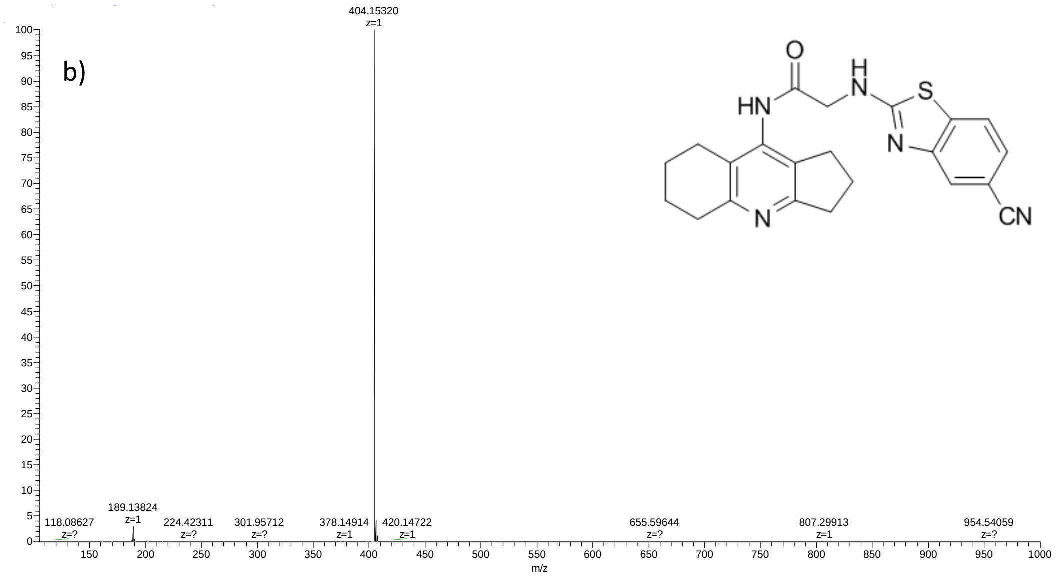

*Figure S 37. HPLC-MS of compound* ***7e*** *a) LC-UV chromatogram b) HRMS mass spectra*

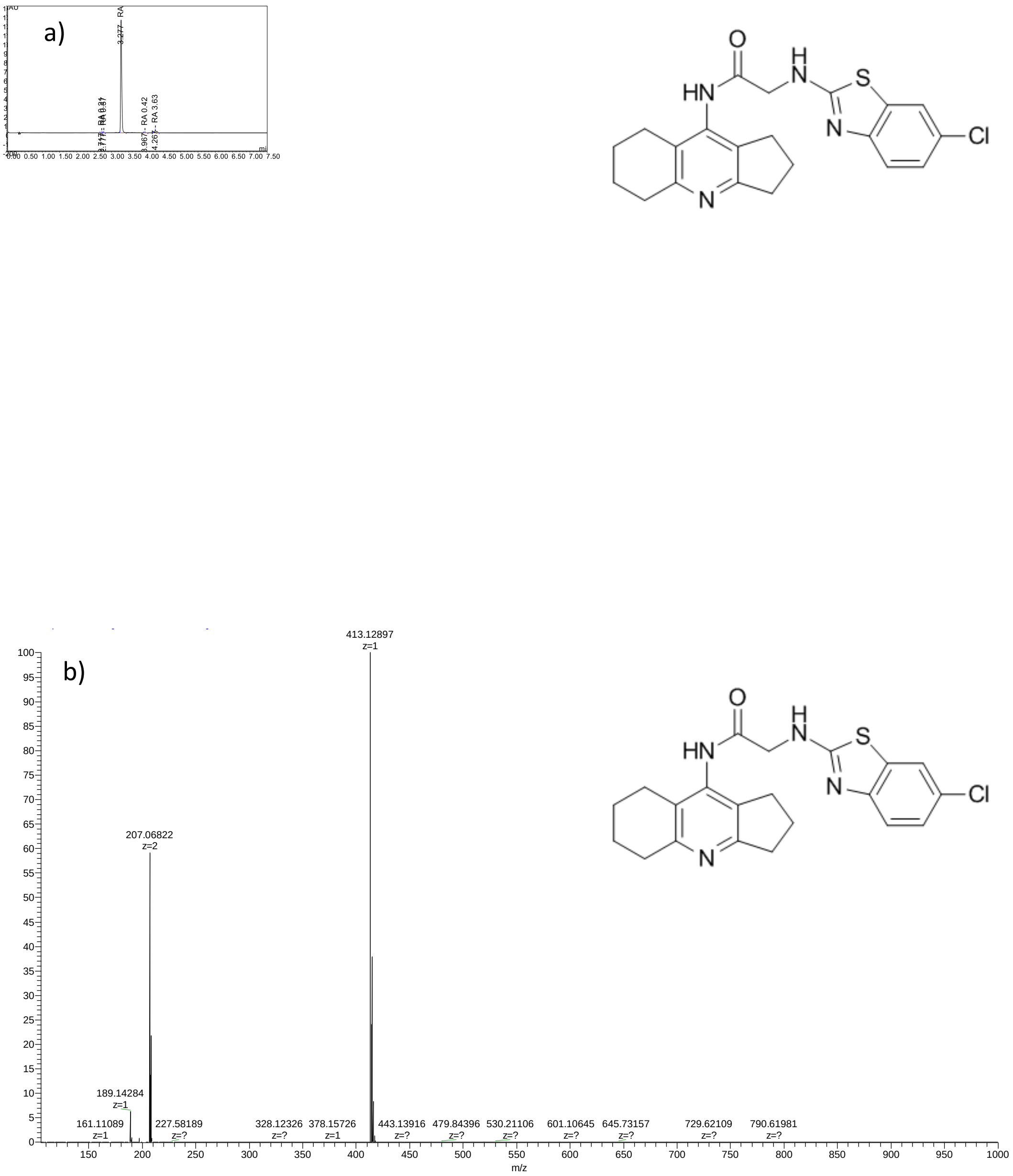

*Figure S 38. HPLC-MS of compound* ***7f*** *a) LC-UV chromatogram b) HRMS mass spectra*

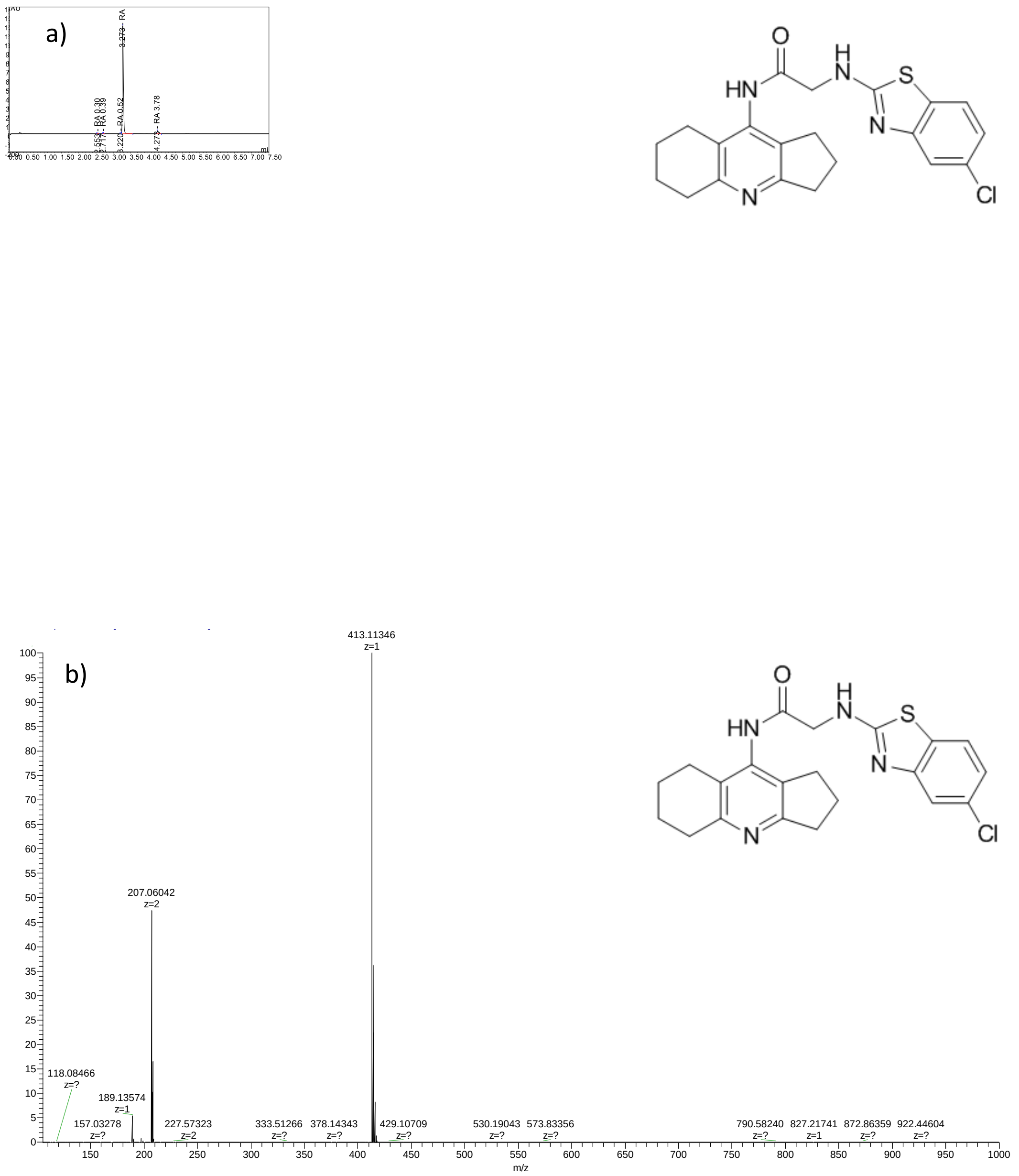

*Figure S 39. HPLC-MS of compound **7g** a) LC-UV chromatogram b) HRMS mass spectra*

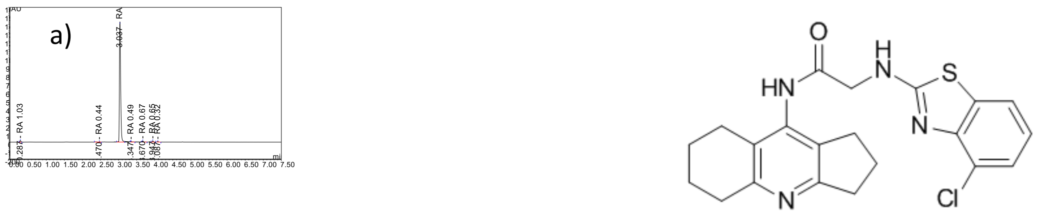

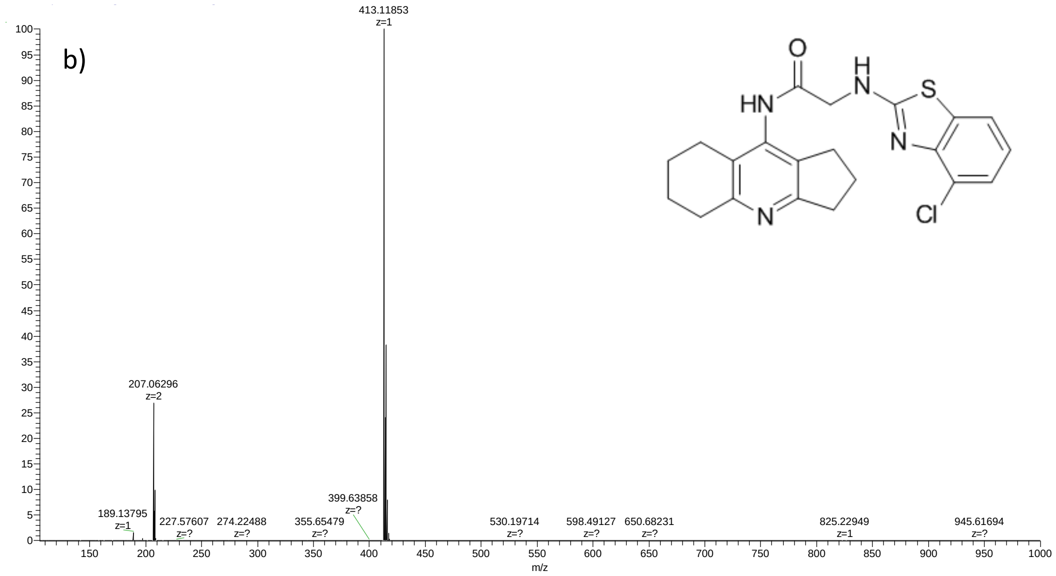

*Figure S 40. HPLC-MS of compound **7h** a) LC-UV chromatogram b) HRMS mass spectra*

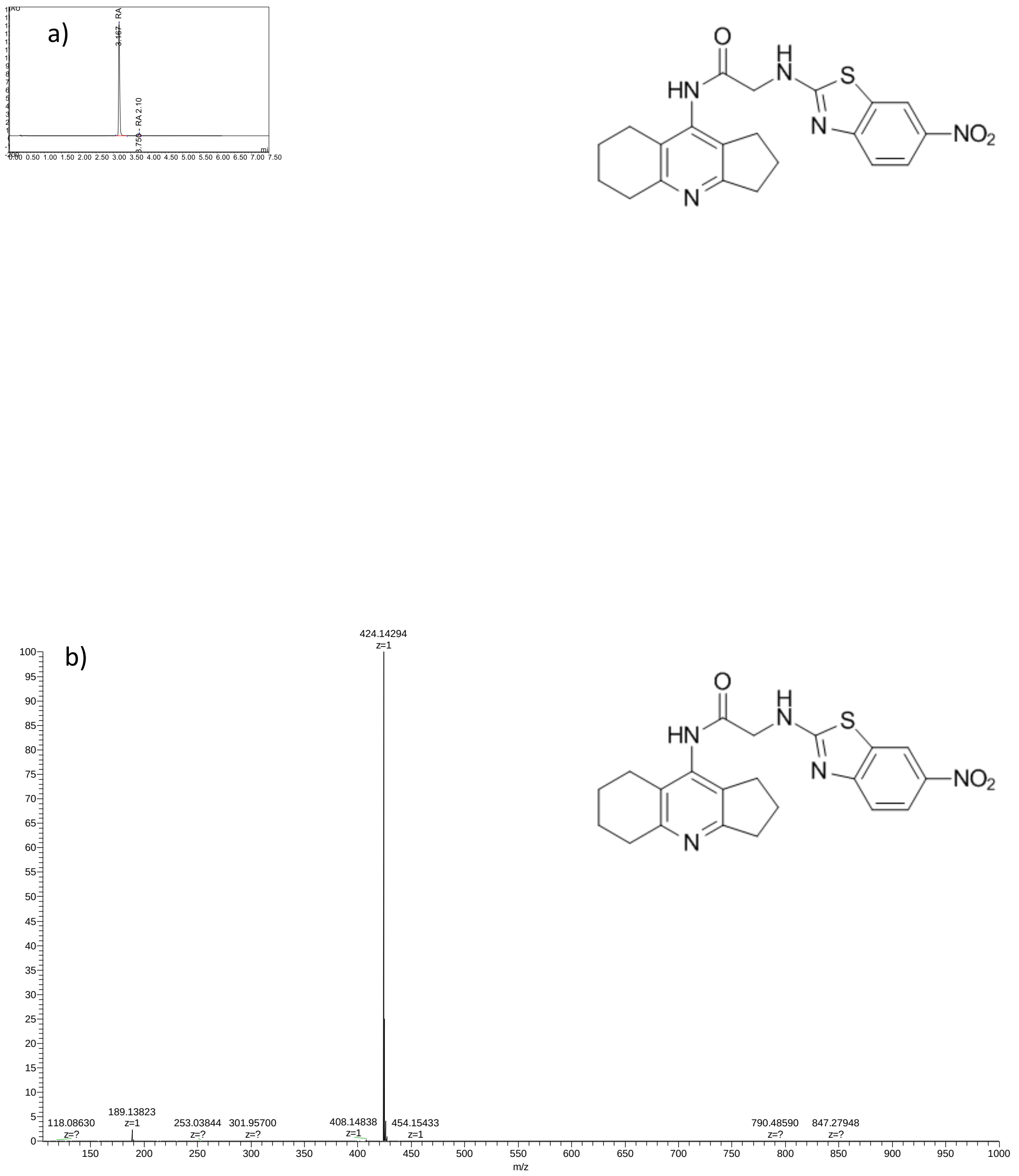

*Figure S 41. HPLC-MS of compound **7i** a) LC-UV chromatogram b) HRMS mass spectra*

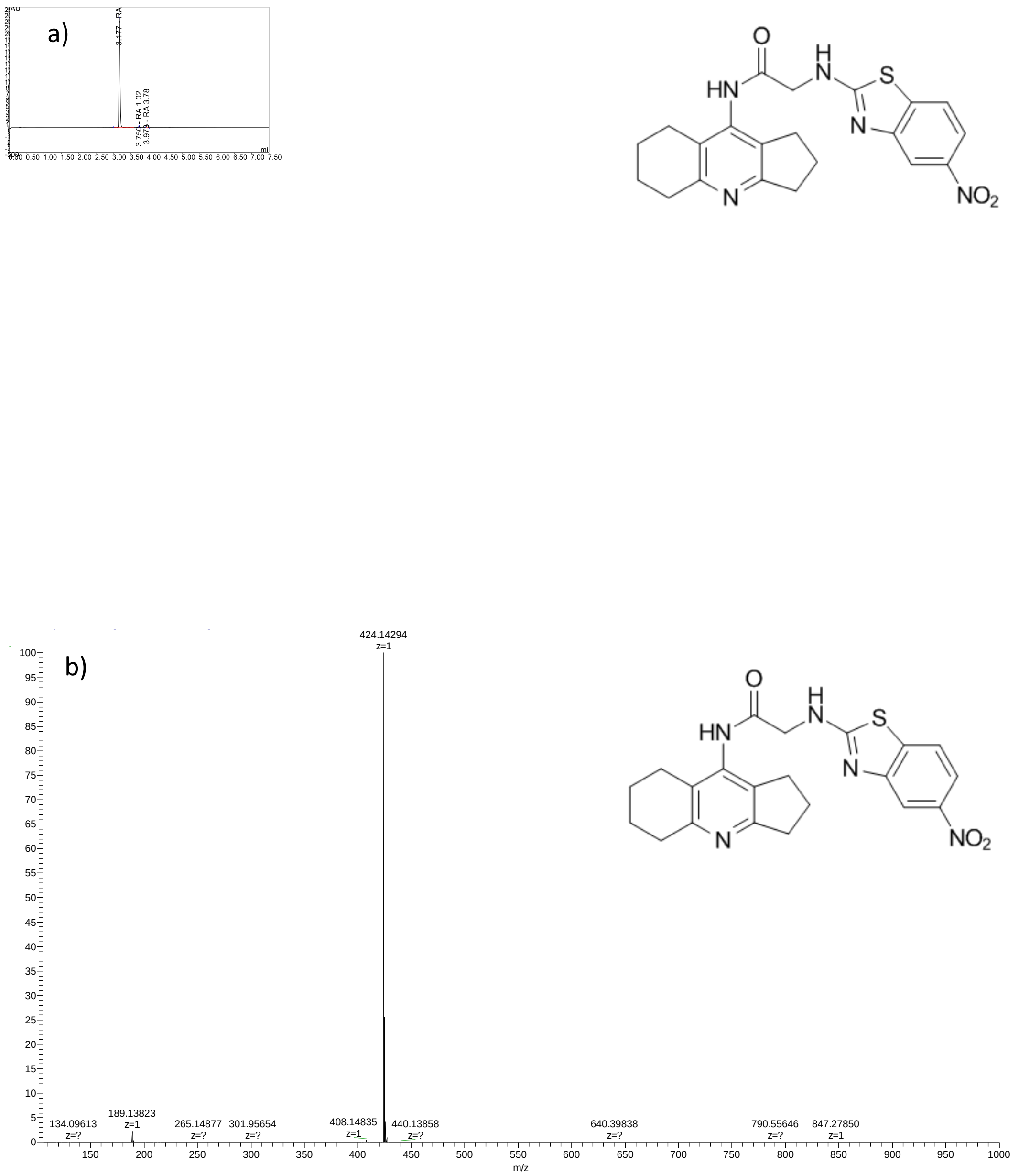

*Figure S 42. HPLC-MS of compound **7j** a) LC-UV chromatogram b) HRMS mass spectra*

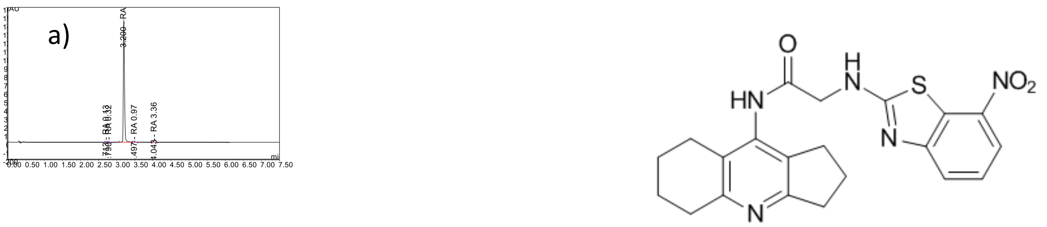

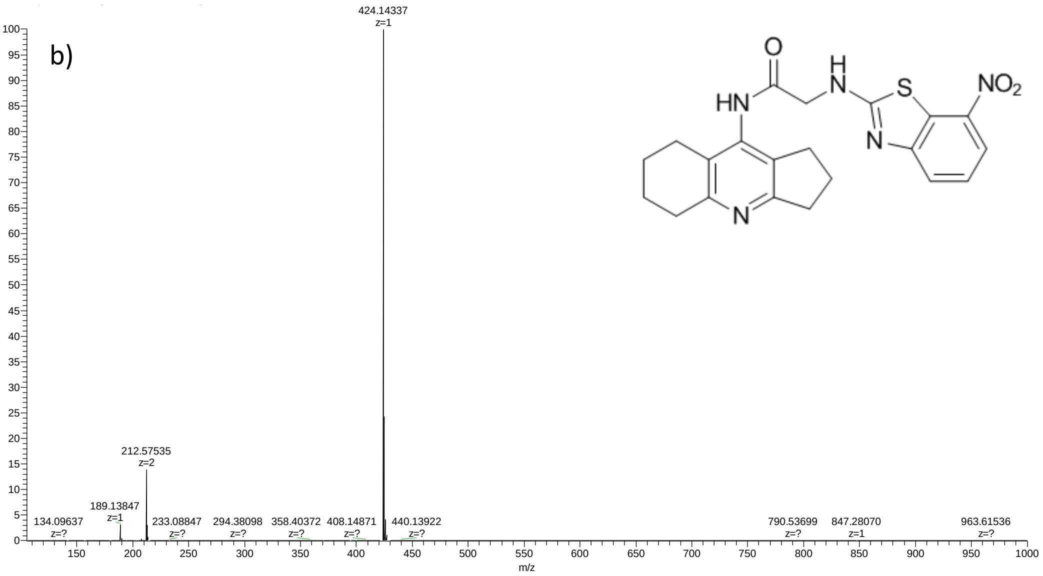

*Figure S 43. HPLC-MS of compound **7k** a) LC-UV chromatogram b) HRMS mass spectra*

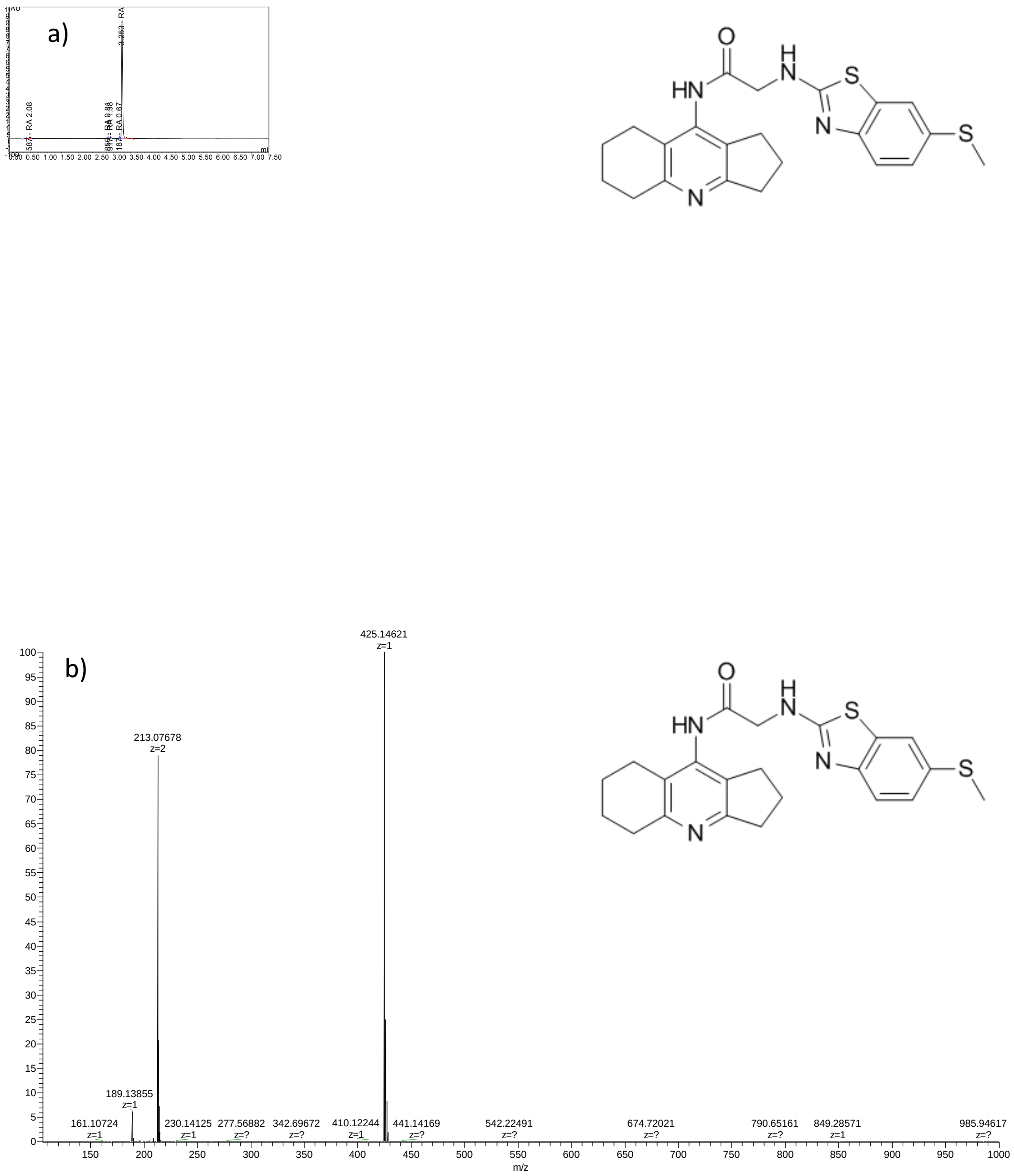

*Figure S 44. HPLC-MS of compound **7l** a) LC-UV chromatogram b) HRMS mass spectra*

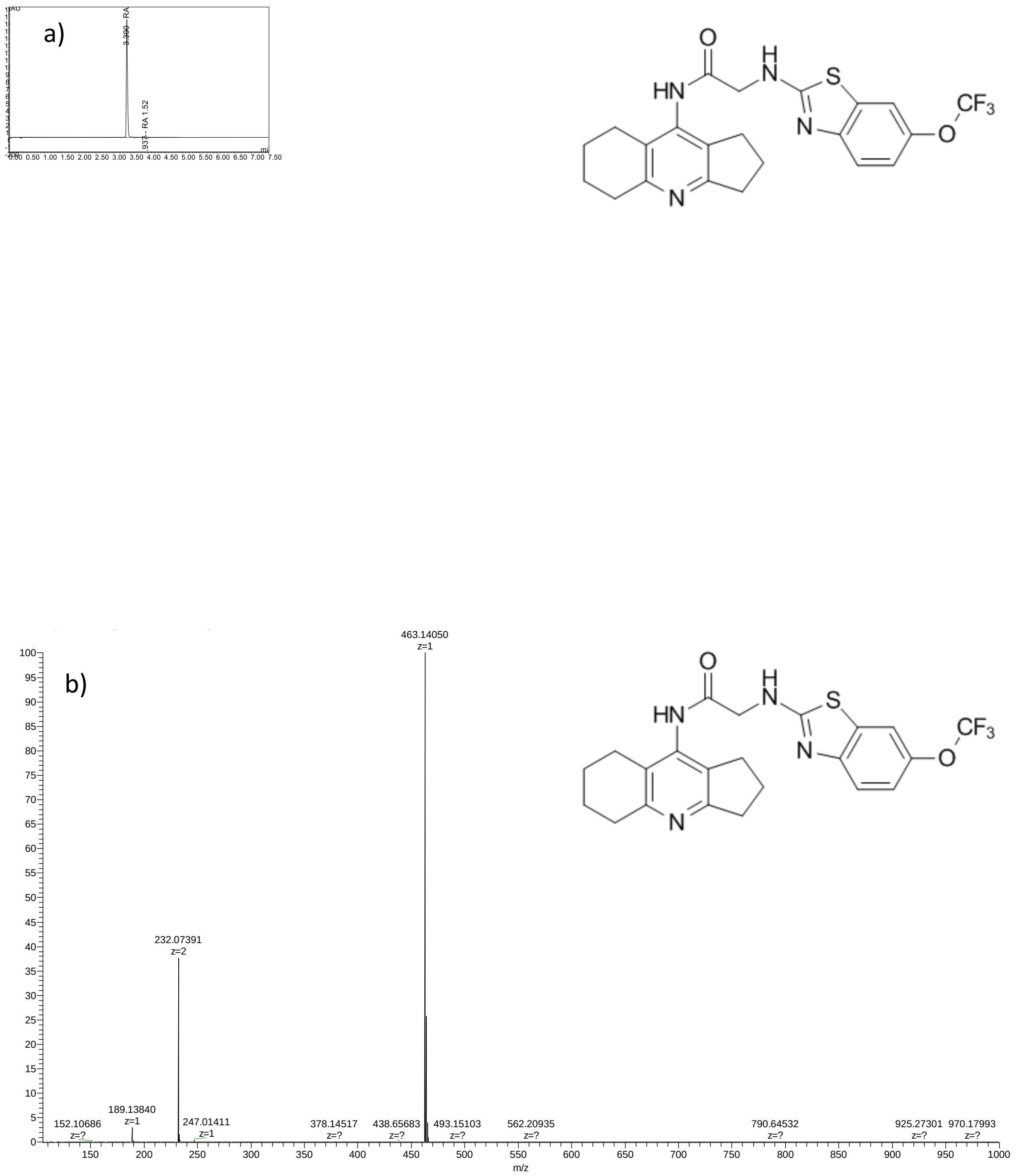

*Figure S 45. HPLC-MS of compound **7m** a) LC-UV chromatogram b) HRMS mass spectra*

### 9. Supporting data for *in silico* studies

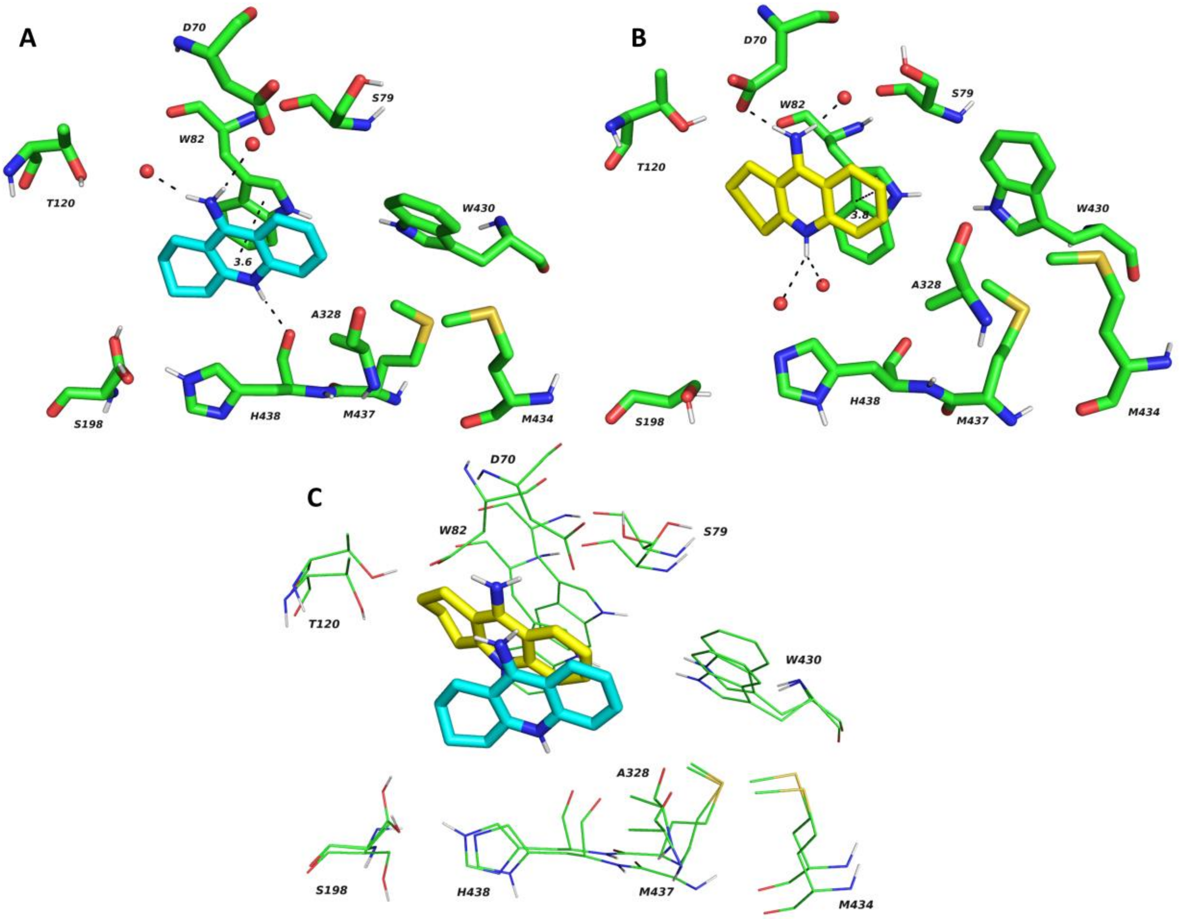

Figure S 46. (A) The crystal structure of THA (blue) bound to BChE active site (PDB ID: 4BDS) [16]. (B) Top-scored docking pose of tacrine (yellow) in the BChE active. (C) Superimposed structure of crystal and docked tacrine molecule with adjacent amino acid residues. The figure was created with The PyMOL Molecular Graphics System, v. 2.5.2.

[5] E. Nepovimova, J. Korabecny, R. Dolezal, K. Babkova, A. Ondrejicek, D. Jun, V. Sepsova, A. Horova, M. Hrabinova, O. Soukup, N. Bukum, P. Jost, L. Muckova, J. Kassa, D. Malinak, M. Andrs, K. Kuca, Tacrine–Trolox Hybrids: A Novel Class of Centrally Active, Nonhepatotoxic Multi-

Target-Directed Ligands Exerting Anticholinesterase and Antioxidant Activities with Low In Vivo Toxicity, J. Med. Chem. 58 (2015) 8985–9003. https://doi.org/10.1021/acs.jmedchem.5b01325.
[6] G. Karabanovich, J. Roh, O. Soukup, I. Pávková, M. Pasdiorová, V. Tambor, J. Stolaříková, M. Vejsová, K. Vávrová, V. Klimešová, A. Hrabálek, Tetrazole regioisomers in the development of nitro group-containing antitubercular agents, MedChemComm. 6 (2015) 174–181. https://doi.org/10.1039/C4MD00301B.
[7] L. Di, E.H. Kerns, K. Fan, O.J. McConnell, G.T. Carter, High throughput artificial membrane permeability assay for blood-brain barrier, Eur. J. Med. Chem. 38 (2003) 223–232. https://doi.org/10.1016/s0223-5234(03)00012-6.
[8] E. Mezeiova, J. Janockova, R. Andrys, O. Soukup, T. Kobrlova, L. Muckova, J. Pejchal, M. Simunkova, J. Handl, P. Micankova, J. Capek, T. Rousar, M. Hrabinova, E. Nepovimova, J.L. Marco-Contelles, M. Valko, J. Korabecny, 2-Propargylamino-naphthoquinone derivatives as multipotent agents for the treatment of Alzheimer's disease, Eur. J. Med. Chem. 211 (2021) 113112. https://doi.org/10.1016/j.ejmech.2020.113112.
[9] M. Gupta, H.J. Lee, C.J. Barden, D.F. Weaver, The Blood–Brain Barrier (BBB) Score, J. Med. Chem. 62 (2019) 9824–9836. https://doi.org/10.1021/acs.jmedchem.9b01220.
[10] A. Daina, O. Michielin, V. Zoete, SwissADME: a free web tool to evaluate pharmacokinetics, drug-likeness and medicinal chemistry friendliness of small molecules, Sci. Rep. 7 (2017) 42717. https://doi.org/10.1038/srep42717.
[11] A. Daina, O. Michielin, V. Zoete, SwissTargetPrediction: updated data and new features for efficient prediction of protein targets of small molecules, Nucleic Acids Res. 47 (2019) W357–W364. https://doi.org/10.1093/nar/gkz382.
[12] C.A. Lipinski, Lead- and drug-like compounds: the rule-of-five revolution, Drug Discov. Today Technol. 1 (2004) 337–341. https://doi.org/10.1016/j.ddtec.2004.11.007.
[13] Y.C. Martin, A Bioavailability Score, J. Med. Chem. 48 (2005) 3164–3170. https://doi.org/10.1021/jm0492002.
[14] J.B. Baell, G.A. Holloway, New Substructure Filters for Removal of Pan Assay Interference Compounds (PAINS) from Screening Libraries and for Their Exclusion in Bioassays, J. Med. Chem. 53 (2010) 2719–2740. https://doi.org/10.1021/jm901137j.
[15] R. Brenk, A. Schipani, D. James, A. Krasowski, I.H. Gilbert, J. Frearson, P.G. Wyatt, Lessons Learnt from Assembling Screening Libraries for Drug Discovery for Neglected Diseases, ChemMedChem. 3 (2008) 435–444. https://doi.org/10.1002/cmdc.200700139.
[16] F. Nachon, E. Carletti, C. Ronco, M. Trovaslet, Y. Nicolet, L. Jean, P.-Y. Renard, Crystal structures of human cholinesterases in complex with huprine W and tacrine: elements of specificity for anti-Alzheimer's drugs targeting acetyl- and butyryl-cholinesterase, Biochem. J. 453 (2013) 393–399. https://doi.org/10.1042/BJ20130013.
